# Neurogliaform Cells Exhibit Laminar-specific Responses in the Visual Cortex and Modulate Behavioral State-dependent Cortical Activity

**DOI:** 10.1101/2024.06.05.597539

**Authors:** Shuhan Huang, Daniella Rizzo, Sherry Jingjing Wu, Qing Xu, Leena Ziane, Norah Alghamdi, David A. Stafford, Tanya L. Daigle, Bosiljka Tasic, Hongkui Zeng, Leena Ali Ibrahim, Gord Fishell

## Abstract

Neurogliaform cells are a distinct type of GABAergic cortical interneurons known for their “volume transmission” output property. However, their activity and function within cortical circuits remain unclear. Here, we developed two genetic tools to target these neurons and examine their function in the primary visual cortex. We found that the spontaneous activity of neurogliaform cells positively correlated with locomotion. Silencing these neurons increased spontaneous activity during locomotion and impaired visual responses in L2/3 pyramidal neurons. Furthermore, the contrast-dependent visual response of neurogliaform cells varies with their laminar location and is constrained by their morphology and input connectivity. These findings demonstrate the importance of neurogliaform cells in regulating cortical behavioral state-dependent spontaneous activity and indicate that their functional engagement during visual stimuli is influenced by their laminar positioning and connectivity.

## INTRODUCTION

GABAergic cortical interneurons (cINs) play a crucial role in regulating cortical function through inhibition. Despite their relatively small numbers, these neurons are vital for controlling the activity of neuronal circuits, and their dysfunction can lead to a variety of neurological disorders. Over the past two decades, it has become recognized that they are comprised by four major cardinal types, which can be distinguished based on their expression of the molecular markers: parvalbumin (PVALB), somatostatin (SST), vasoactive-intestinal peptide (VIP), and lysosome-associated membrane protein 5 (LAMP5). Recent studies have extensively characterized their range of diversity, which is formed during development and results in each class possessing unique gene expression, morphologies, physiological properties, connectivities, and *in vivo* function^1–5^. Among the four cardinal types, LAMP5+ cINs have been the least investigated. This reflects the historic lack of genetic tools to specifically target them. Despite being initially discovered by Ramón y Cajal over 125 years ago^6^, their contribution to cortical function within neural circuits remains uncertain. In addition, it is unclear whether LAMP5+ cINs residing in different layers exhibit distinct functional activities.

Neurogliaform cells, also referred to as dwarf, spiderweb, or arachniform cells in the literature, are classified as the LAMP5+ cardinal type cINs and constitute the majority of LAMP5+ cINs^7–11^. Despite exhibiting heterogeneous morphology within the population in accordance with their laminar location (they are elongated in layer 1 (L1) while spherical in layers 2/3 (L2/3))^12^, neurogliaform cells share several commonalities in their dendritic and axonal organization^1,12–17^. They possess short, spherically-distributed and sparsely branched dendrites. Their axons are notably thin and form a highly ramified network within their cortical layer, occasionally extending into adjacent layers.

Neurogliaform cells exhibit a distinct late-spiking electrophysiological signature, characterized by a slow ramp of depolarization until the first spike occurs at rheobase^12,13,18,19^. Although their axonal varicosities contain synaptic vesicles, they seldom form conventional synaptic contacts and instead are thought to signal through “volume transmission” ^17,18,20^. Activation of neurogliaform cells induces slow and prolonged inhibitory postsynaptic potentials (IPSPs) in most adjacent neurons, mediated by extra-synaptically located GABAA and GABAB receptors^14,18,21^. Neurogliaform cells are electrically coupled to not only other neurogliaform cells but also other cIN types, suggesting they may monitor inhibitory network activity and regulate synchrony via gap junctions^22–24^.

Nevertheless, the *in vivo* activity and function of neurogliaform cells remain unclear. Given their “volume transmission” output property that results in them mediating prolonged and non-selective broad inhibition, we hypothesize that these neurons suppress baseline cortical activity when recruited, and contribute to maintaining a high signal-to-noise ratio (SNR) in cortical function. Additionally, their short dendritic structure suggests that laminar-specific inputs selectively target these cINs within their respective layers.

In this study, we investigated the functional consequence of removing LAMP5+ cINs and compared their activity and connectivity across different cortical layers. Our goal was to (1) understand how LAMP5+ cINs are recruited during various behavioral states, and (2) determine whether these cINs in different cortical layers might be recruited differently in specific contexts. We used the primary visual cortex (V1) in mice as a model to address these questions. Firstly, we developed and validated two new genetic strategies enabling selective targeting of LAMP5+ cINs. Secondly, we silenced LAMP5+ cINs, which led to increased spontaneous activity during locomotion, increased synchronization, and impaired visual responses in L2/3 pyramidal neurons (PYN^L23^) in V1. We also found that LAMP5+ cINs exhibited spontaneous activity that positively correlated with the animal’s active behavioral states. Finally, we observed disparities in visually evoked responses between LAMP5+ cINs in L1 (LAMP5^L1^) and L2/3 (LAMP5^L23^). Specifically, while LAMP5^L23^ showed a preference for high-contrast visual stimuli during locomotion, LAMP5^L1^ preferred low-contrast visual stimuli. We investigated whether the circuit connectivity of LAMP5+ cINs in V1 explains these differences and found that the inputs to these neurons vary depending on their laminar location. In summary, our study introduced new tools for accessing this underexplored population, and revealed the recruitment and function of LAMP5+ cINs in cortical circuits.

## RESULTS

### Genetic Targeting of LAMP5+ cINs in Mouse V1

Single cell RNA sequencing and spatial transcriptomic data^10,25^ have revealed that the majority of LAMP5+ cINs express NPY **(Extended Data 1a1)**, confirming their identity as neurogliaform cells described in previous literature^6,11–13,18^. Interestingly, among all cINs, these neurons also exhibited the highest expression of GAD2^26,27^ **(Extended Data 1a2)**, enabling them to exert powerful inhibition^14,18^. While many LAMP5+ cINs are located in L1, we estimate that ∼60% of this cardinal type could reside in L2-6 **(Figure 1a)**. To date, *in vivo* studies of LAMP5+ cINs have primarily focused on those within L1^28–32^, however, considerably less attention has been given to those in the other cortical layers^33^. This is largely because we currently lack genetic tools to access LAMP5+ cINs in all layers, including layers 2-6 (LAMP5^L2-6^).

**Figure 1.**
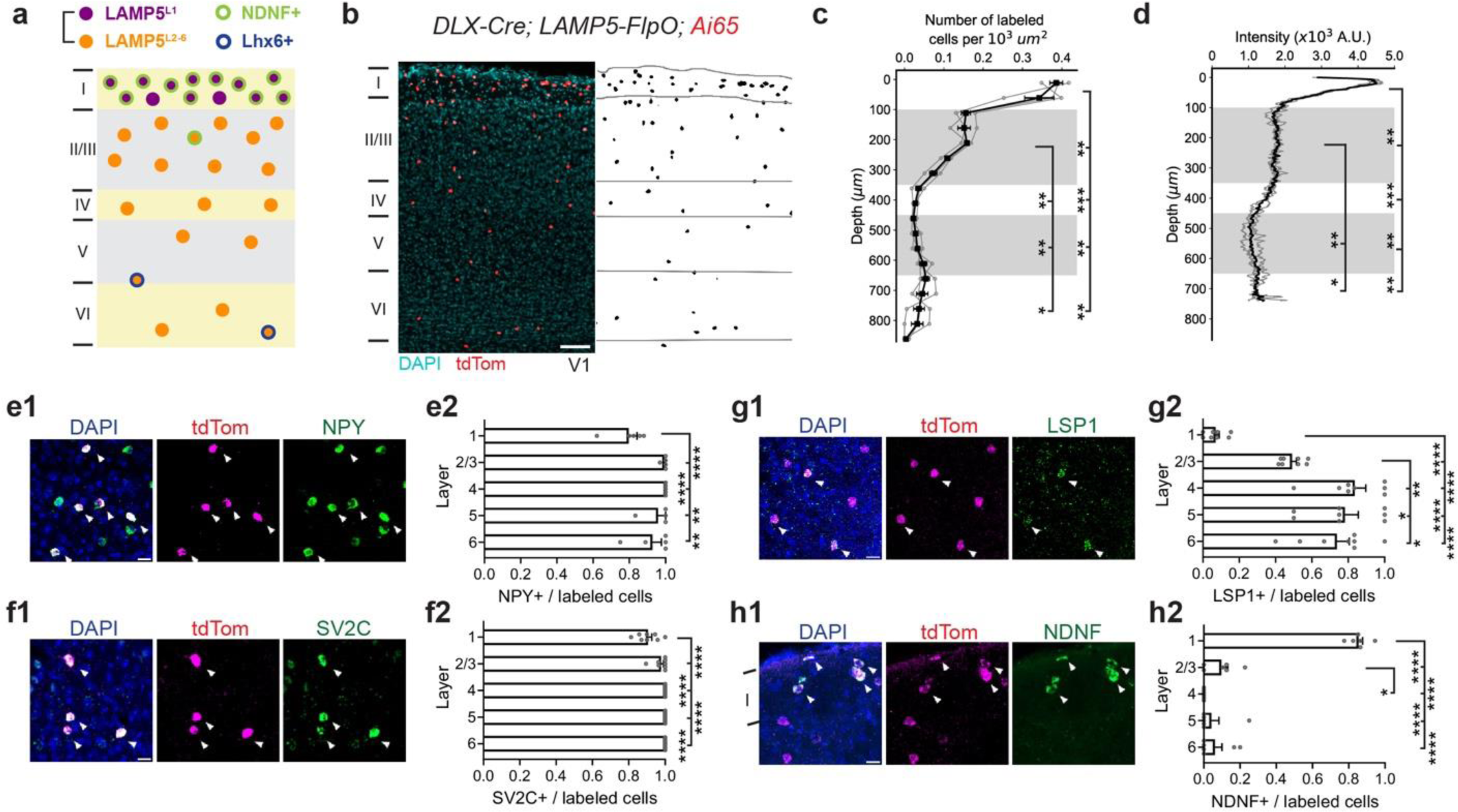
D*LX-Cre; LAMP5-FlpO* targets all LAMP5+ cINs. (a) Schematic illustration of the distribution of LAMP5+ cINs in V1, indicating their various subtypes. (b) Example image of *DLX-Cre; LAMP5-FlpO; Ai65(RCFL-tdT)* genetic labeling in V1 showing the distribution of LAMP5+ cINs (left) and segmented cell bodies (right). Scale bar = 100 μm. (c) Quantification for the number of labeled cells per 10^3^ μm^2^ and (d) the intensity of labeled neurites in 10^3^ arbitrary units (A.U.) across cortical layers. Gray indicates L2/3 and L5. The 40 μm thick coronal V1 sections were divided into 50 μm vertical bands for cell density measu rement. Each gray line represents data from an individual animal. (e-h) Representative RNAscope assays images in V1 taken with confocal microscopy and bar plots quantifying the ratio of marker gene-positive cells to all tdTom labeled cells in each cortical layer. Each panel displays an overlay figure (left) with nuclei dye DAPI (in blue), tdTom-labeled cells from *DLX-Cre; LAMP5-FlpO; Ai65(RCFL-tdT)* (middle, in red), and the marker gene expression detected by RNAscope assay (right, in green). Scale bar = 20 μm. Each dot represents data from one coronal V1 section. Arrow indicates tdTom and marker gene co-localized cells. (e1-e2) NPY; (f1-f2) SV2C; (g1-g2) LSP1; (h1-h2) NDNF. While NPY is a classical marker for identifying neurogliaform cells in rodents, it is not exclusively restricted to LAMP5+ cINs. Moreover, although LSP1 serves as a specific marker gene for the Lamp5/Lsp1 cluster, it is expressed in only half of the neurons within this population. Data from N = 3 animals. Error bar represents SEM. Repeated measures ANOVA (c-d, g2-h2) or mixed-effects model (e2) followed by uncorrected Fisher’s LSD were used for testing statistical significance. See supplementary data 1 - Table 1 for statistics.

To address this issue, we developed and validated two genetic labeling strategies to label all LAMP5+ cINs within the mouse V1. In the first strategy, *DLX-Cre; LAMP5-FlpO* alleles were bred with an intersectional reporter *Ai65(RCFL-tdT*) **(Figure 1b)**. While LAMP5 is primarily restricted to LAMP5+ cINs, it is also expressed within a subset of excitatory neurons. The use of the *DLX-Cre* allele restricted the expression to cINs, and the *LAMP5-FlpO* allele limited it to the LAMP5+ cIN population. Our genetic labeling strategy demonstrated that LAMP5+ cINs are primarily enriched within L1 and L2/3 (∼42.5% in L1 and ∼38% in L2/3) **(Figure 1b-d)**^10,12^. Notably, although the level of neurite fluorescence was equivalent across L2/3 **(Figure 1d)**, the density of labeled cell bodies was higher in L2 than in L3 **(Figure 1c)**. By contrast, significantly fewer cells were observed in L4-6 (∼19.5%), accompanied by the lowest intensity of neurite fluorescent signals in L5 **(Figure 1c,d, Extended Data 1b)**.

To validate the specificity of this targeting strategy, we examined the colocalization between the genetically labeled neurons and marker genes we identified for LAMP5+ cINs using RNAscope. Over 95% of the labeled neurons were DLX1+ **(Extended Data 1d)**, confirming their identity as cINs. Moreover, most of the labeled neurons express NPY (80% in L1, 93-100% in L2-6, **Figure 1e)** and SV2C (90% in L1 and 98-100% in L2-6, **Figure 1f)**, markers enriched in neurogliaform cells. The distribution of two other markers, LSP1 (7% in L1, 49% in L2/3, 74-83% in L4-6, **Figure 1g)** and NDNF (85% in L1, 10% in L2/3, 6% in L4-6, **Figure 1h)**, was consistent with distinct LAMP5+ cIN subtypes in L2-6 and L1, respectively. These results aligned with the expected ratios from transcriptomic data. Additionally, almost all SV2C+ cells in L1-4 **(Extended Data 1c1)** and NDNF+ cells in L1-3 **(Extended Data 1c2)** were labeled, underscoring the robustness of this targeting strategy. To rule out nonspecific labeling in other cIN types, we assessed co-expression with LHX6 (detects PV+ and SST+ cINs), VIP and SNCG, and observed minimal overlap **(Extended Data 1e-g)**. Furthermore, the distribution of LAMP5+ cINs appears similar across different neocortical regions, including the primary somatosensory cortex (SSp) barrel field **(Extended Data 1h-j)** and the primary motor cortex (MOp) **(Extended Data 1k-m)**. Collectively, these results suggest that the *DLX-Cre; LAMP5-FlpO* offers specific and robust access to the LAMP5+ cIN population.

While the intersectional *DLX-Cre; LAMP5-FlpO* provides comprehensive genetic access to LAMP5+ cINs, the requirement for multiple alleles reduces its flexibility. Therefore, we devised an alternative strategy using a single allele, *JAM2-Cre*, in conjunction with local viral delivery of Cre-dependent AAV driven by Dlx enhancer^34^ **(Extended Data 2a)**. This approach allows for selective targeting of LAMP5+ cINs in adults (**Extended Data 2b**; see Method - Mouse). We assessed the specificity of this targeting using similar methods. Labeled neurons co-express GAD2 (100%, **Extended Data 2c)**, NPY (82% in L1, 81% in L2/3, 100% in L4-5, 86% in L6, **Extended Data 2d)**, SV2C (76-80%, **Extended Data 2e)**, LSP1 (4.5% in L1, 60-67% in L2-6, **Extended Data 2f)** and NDNF (82% in L1, **Extended Data 2g)**, suggesting this strategy closely mirrors the LAMP5+ cIN labeling achieved with *DLX-Cre; LAMP5-FlpO* approach. As expected, a large proportion of the targeted neurons exhibited late-spiking electrophysiological characteristics, a trait associated with neurogliaform cells **(Extended Data 2i)**^35^. These neurons also displayed typical neurogliaform cell morphology with short spherical dendrites and thin, highly ramified axons **(Extended Data 2j)**^12^. Off-target analysis revealed that this strategy achieves ∼85% specificity, with 13-16% VIP+ in L1-4, 10% PV+ in L4-5, 15% SNCG+ in L6 **(Extended Data 2k-n)**.

In summary, we have developed and validated two intersectional strategies to specifically target LAMP5+ cINs: one utilizing *DLX-Cre; LAMP5-FlpO* and another employing *JAM2-Cre* with a viral tool. Both strategies provide precise and reliable access to LAMP5+ cINs, predominantly located in L1 and L2/3 of the neocortex^12,25^. In subsequent experiments, we used both strategies concurrently, unless one offered significant advantages in terms of convenience or technical feasibility for specific experimental demands.

### LAMP5+ cIN Silencing Increased the Spontaneous Activity of PYN^L23^ in V1

The majority of LAMP5+ cINs are neurogliaform cells, whose activation results in broad and long-lasting inhibition *ex vivo* via “volume transmission” ^12,18,20,21,36^. Nevertheless, the *in vivo* functional contribution of this unique form of synaptic transmission from neurogliaform cells remains unclear.

To investigate the overall role of inhibition provided by LAMP5+ cINs within cortical circuits, we specifically blocked the synaptic transmission from LAMP5+ cINs by crossing the *DLX-Cre; LAMP5-FlpO* with the tetanus toxin effector *RC:PFtox^37^ (*hereafter referred to as *LAMP5::TOX)* **(Figure 2a)**. We then preferentially delivered GCaMP into L2/3 pyramidal neurons (PYN^L23^) by injecting AAV9.CaMKII.GCaMP6f.WPRE.SV40 into the V1 **(Figure 2a,b)**. We conducted two-photon calcium imaging to record the spontaneous activity of PYN^L23^ in V1 of awake, head-fixed, free-running mice, which were viewing a gray screen **(Figure 2a,b)**. As a control, we utilized Cre-negative and/or FlpO-negative mice *(LAMP5::CTR)* **(Figure 2a)**. Silencing LAMP5+ cINs resulted in increased spontaneous activity of PYN^L23^ **(Figure 2c)**. When cross-correlating the spontaneous activity among PYN^L23^ **(Figure 2d)**, we found an increase in both the proportion of significantly correlated neuronal pairs **(Figure 2e)** and the correlation level between neuronal pairs in *LAMP5::TOX* compared with *LAMP5::CTR* **(Figure 2f, Extended Data 3a)**. When we analyzed the activity of PYN^L23^ during different behavioral states (stationary or run), we observed a higher mean activity in *LAMP5::TOX* compared to *LAMP5::CTR* only when the mice were running **(Extended Data 3b)**. As a result, the difference in spontaneous activity of PYN^L23^ between the running and the stationary state was significantly higher in *LAMP5::TOX* **(Figure 2g)**.

**Figure 2.**
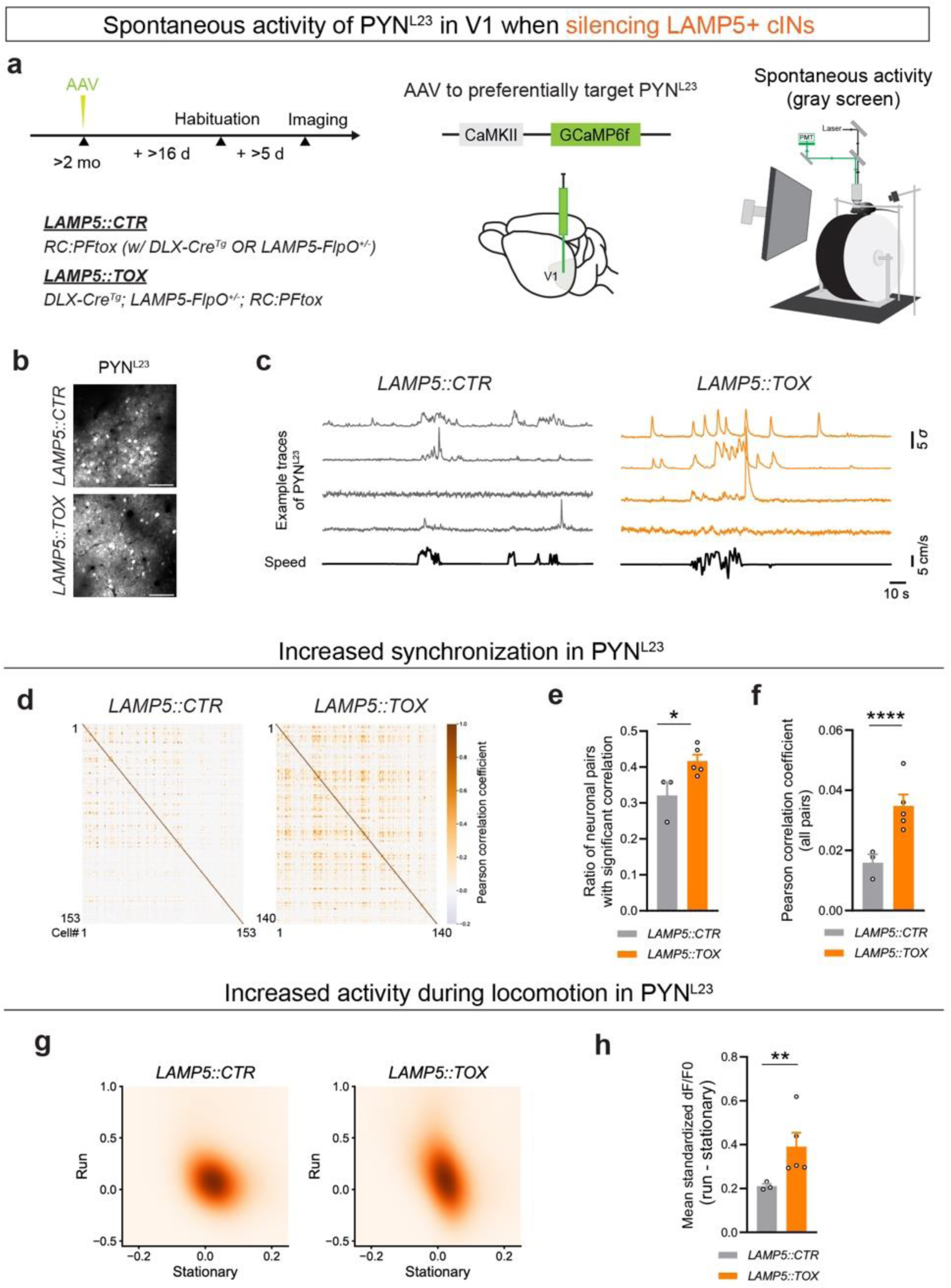
LAMP5+ cIN silencing results in increased spontaneous activity of PYN^L23^ in V1. (a) (left) The experimental animals (*LAMP5::TOX*) were generated by crossing the *DLX-Cre; LAMP5-FlpO* mouse model with *RC:PFtox*, specifically, *DLX-Cre^(Tg)^; LAMP5-FlpO^(+/-)^; RC:PFtox^(Flox/+, Frt/+)^*. Controls (*LAMP5::CTR*) were mice lacking Cre and/or FlpO (either one or both negative) while still carrying the *RC:PFtox^(Flox/+, Frt/+)^* genotype. No notable differences were observed among different control genotypes. (middle) To visualize PYN^L23^ activity, AAV9.CaMKII.GCaMP6f.WPRE.SV40 was injected into V1 during the cranial window implantation surgery to express GCaMP6f preferentially in PYN^L23^. Intrinsic imaging (refer to Methods) was used to confirm the V1 location in each animal before proceeding with *in vivo* two-photon calcium imaging experiments. (right) Illustration of the experimental setup during the spontaneous activity recordings. The animal was presented with a gray screen (uniform mean luminance) during two-photon imaging of the GCaMP signal. (b) Example images from the maximum intensity projection of the two-photon imaging experiments, showing GCaMP6f expression within L2/3 of V1 in *LAMP5::CTR* (upper) and *LAMP5::TOX* (bottom). Scale bar = 100 μm. (c) Example traces from spontaneous activity recordings in both *LAMP5::CTR* (left) and *LAMP5::TOX* (right). The top four traces in each set (gray for *LAMP5::CTR* and orange for *LAMP5::TOX*) represent the smoothed standardized dF/F0 activity of four randomly chosen neurons during a randomly selected time interval for representation. The bottom trace (in black) in each set shows the corresponding locomotion speed of the animal (measured in cm/s). (d) Example heatmap showing Pearson’s correlation coefficient of neuron pairs from a single imaging session using *LAMP5::CTR* (left) or *LAMP5::TOX* (right). (e) Bar plot showing ratio of significantly (p < 0.05) correlated pairs out of all neuron pairs with shuffling (refer to Methods) in *LAMP5::CTR* (gray) and *LAMP5::TOX* (orange). (f) Bar plot showing Pearson’s correlation coefficient of all neuron pairs in *LAMP5::CTR* (gray) and *LAMP5::TOX* (orange). (g) Averaged spontaneous standardized dF/F0 activity for each neuron during either stationary (speed ≤ 1 cm/s) or running (sp eed > 1 cm/s) period for *LAMP5::CTR* (left) and *LAMP5::TOX* (right) groups. The data are fit with a Gaussian kernel for visualization. (h) Bar plot showing the differences in mean standardized dF/F0 activity between running and stationary period, obtained by subtracting the latter from the former, for *LAMP5::CTR* (gray) and *LAMP5::TOX* (orange). Error bar represents SEM. Each dot represents the (averaged) result from an individual animal. Mann-Whitney test (e) and hierarchical bootstrap (f,h-i) were used for testing statistical significance. See supplementary data 1 - Table 2 for statistics.

VIP+ cINs are most prevalent in L2/3 and their spontaneous activities are highly correlated with locomotion speed^38^. However, unlike the results with LAMP5+ cIN silencing, VIP+ cIN silencing (**Extended Data 3c-e)** did not affect the highly desynchronized property of PYN^L23^ activity (**Extended Data 3f-i)**. We also did not observe any significant changes in the spontaneous activity of PYN^L23^ during different behavioral states with VIP+ cIN silencing (**Extended Data 3j-l)**.

Collectively, these findings suggest that LAMP5+ cINs play a critical role in desynchronizing and maintaining sparse spontaneous activity of PYN^L23^ in V1, particularly during active behavioral state.

### LAMP5+ cIN Silencing Results in Impaired Visual Response of PYN^L23^ in V1

To explore the potential impact of the increased spontaneous activity in *LAMP5::TOX* on the visual response properties of PYN^L23^ in V1, we exposed the mice to full-field visual stimuli consisting of moving gratings at 12 different orientations (0° to 330°, in 30° increments), while simultaneously recording the activity of PYN^L23^ in V1 **(Figure 3a,b)**. Interestingly, while the proportion of responsive neurons remained unchanged **(Extended Data 4a)**, we found that the average peak response at the preferred orientation was diminished in *LAMP5::TOX* compared to *LAMP5::CTR* **(Figure 3c,d, Extended Data 4b)**. Given the visual responses were calculated by subtracting the baseline activity from the mean activity during visual stimulus, this reduction could be due to the elevated baseline activity in *LAMP5::TOX*, which was significantly higher **(Extended Data 4c)**. Furthermore, the activity during the visual stimulus period was also reduced in *LAMP5::TOX* **(Extended Data 4c)**. These findings suggest that both the elevated baseline and reduced stimulus period activities contributed to the diminished visual responses in *LAMP5::TOX*.

**Figure 3.**
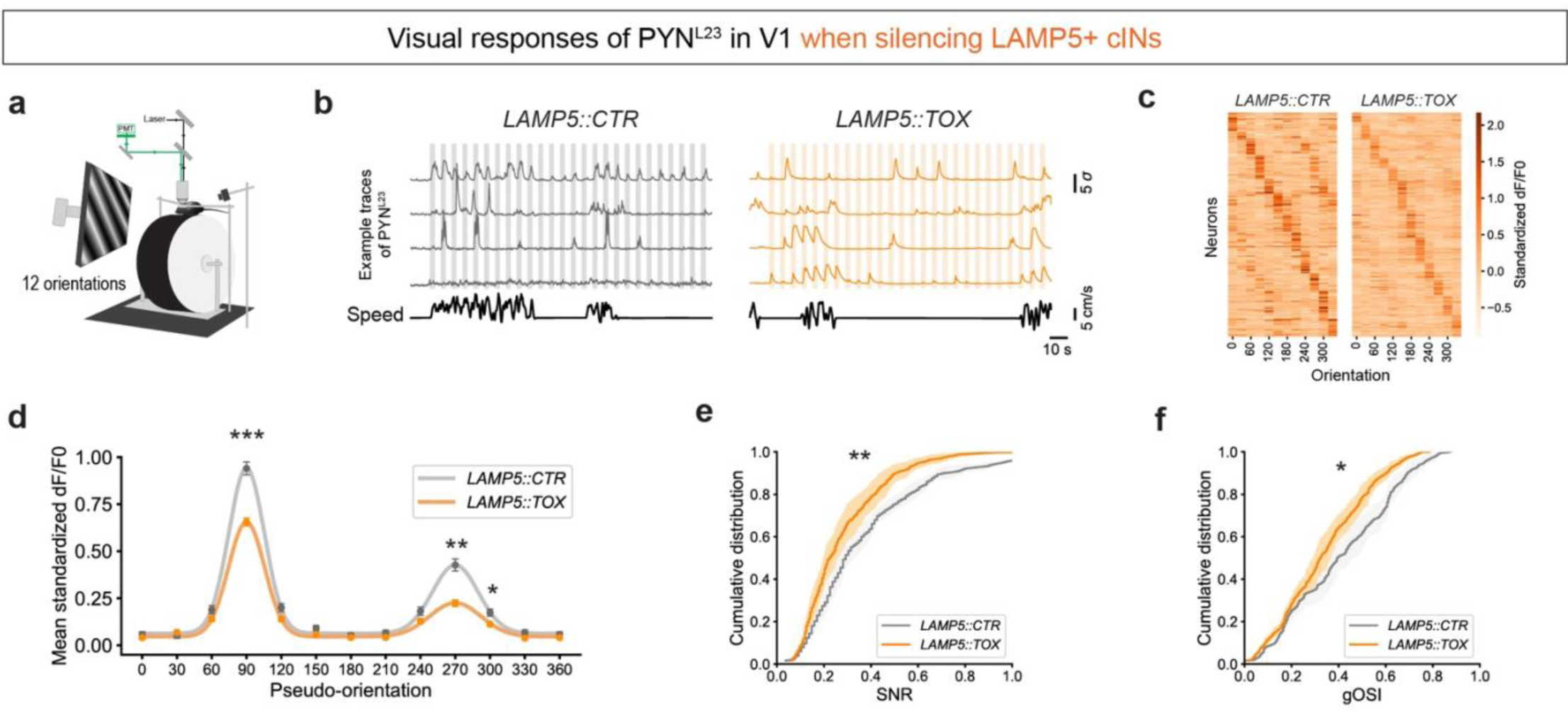
LAMP5+ cIN silencing results in impaired visual response properties of PYN^L23^ in V1. (a) Illustration of the experimental setup during the recording of visual responses. The animal was presented with moving gratings on the screen, while the GCaMP signal in PYN^L23^ was imaged with two-photon microscopy. Each trial was 6s, consisting of 2s full-field moving gratings with orientations displayed randomly, a fixed contrast (80%), spatial frequency (0.04 cpd) and temporal frequency (1 Hz). 4s inter-stimulus-intervals of gray screen (mean luminance) was presented. 12 orientations were examined: 0°, 30°, 60°, 90°, 120°, 150°, 180°, 210°, 240°, 270°, 300°, 330°. (b) Example traces of PYN^L23^ responses from the *LAMP5::CTR* (gray) or *LAMP5::TOX* (orange) mice. The top four traces in each set represent the standardized dF/F0 activity (smoothed for representation) of four randomly chosen neurons during a randomly selected time interval. The bottom trace (in black) in each set showed the corresponding locomotion speed of the animal (measured in cm/s). Colored fill (gray or orange) indicates the presence of moving gratings. (c) Heatmap showing averaged visual responses to various tested orientations (columns) of all recorded neurons (rows) sorted by their preferred orientation for *LAMP5::CTR* (left) and *LAMP5::TOX* (right). (d) Averaged visual responses to orientations (pseudo-90° indicates the preferred orientation for a neuron) in *LAMP5::CTR* (gray) and *LAMP5::TOX* (orange). (e) Cumulative ratio of signal-noise-ratio (SNR) for *LAMP5::CTR* (gray) and *LAMP5::TOX* (orange). (f) Cumulative ratio of signal correlation of visual responses for *LAMP5::CTR* (gray) and *LAMP5::TOX* (orange). (g) Cumulative ratio of global orientation selective index (gOSI) for *LAMP5::CTR* (gray) and *LAMP5::TOX* (orange). Error bar represents SEM. Hierarchical bootstrap was used for testing statistical significance. See supplementary data 1 - Table 3 for statistics.

Consequently, visual responses in PYN^L23^ of *LAMP5::TOX* exhibited a decreased SNR compared to *LAMP5::CTR* **(Figure 3e).** In addition, *LAMP5::TOX* showed a decrease in the global orientation selectivity index (gOSI) **(Figure 3f)**, suggesting a broader visual tuning curve in PYN^L23^ in *LAMP5::TOX*. No significant changes were found in signal correlation **(Extended Data 4d)**, noise correlation **(Extended Data 4e)** or the distribution of preferred orientation **(Extended Data 4f)**. We observed a tendency toward an increase in the direction selective index (DSI) in *LAMP5::TOX* (**Extended Data 4g)**.

Thus, when LAMP5+ cINs are silenced, visual responses in PYN^L23^ exhibit reduced sharpness in orientation tuning and decreased SNR. These results suggest that LAMP5+ cINs play a pivotal role in maintaining PYN^L23^’s sensitivity to visual stimuli in V1.

### Spontaneous Activity of LAMP5+ cINs Correlates with Behavioral States

Silencing LAMP5+ cINs led to increased spontaneous activity in PYN^L23^ during locomotion **(Figure 2)**, suggesting that these cINs may be recruited during active behavioral states. To understand how LAMP5+ cINs are engaged in different behavioral states, we performed *in vivo* two-photon calcium imaging and recorded the spontaneous activity of these cINs in V1 of awake behaving mice, while presenting them with a gray screen **(Figure 4a)**. The activity of LAMP5^L23^ and LAMP5^L1^, where most of these cINs reside, were monitored via GCaMP7s, which was specifically expressed in these cINs using the *DLX-Cre; LAMP5-FlpO; Ai195* mouse model **(Figure 4b,c)**.

**Figure 4.**
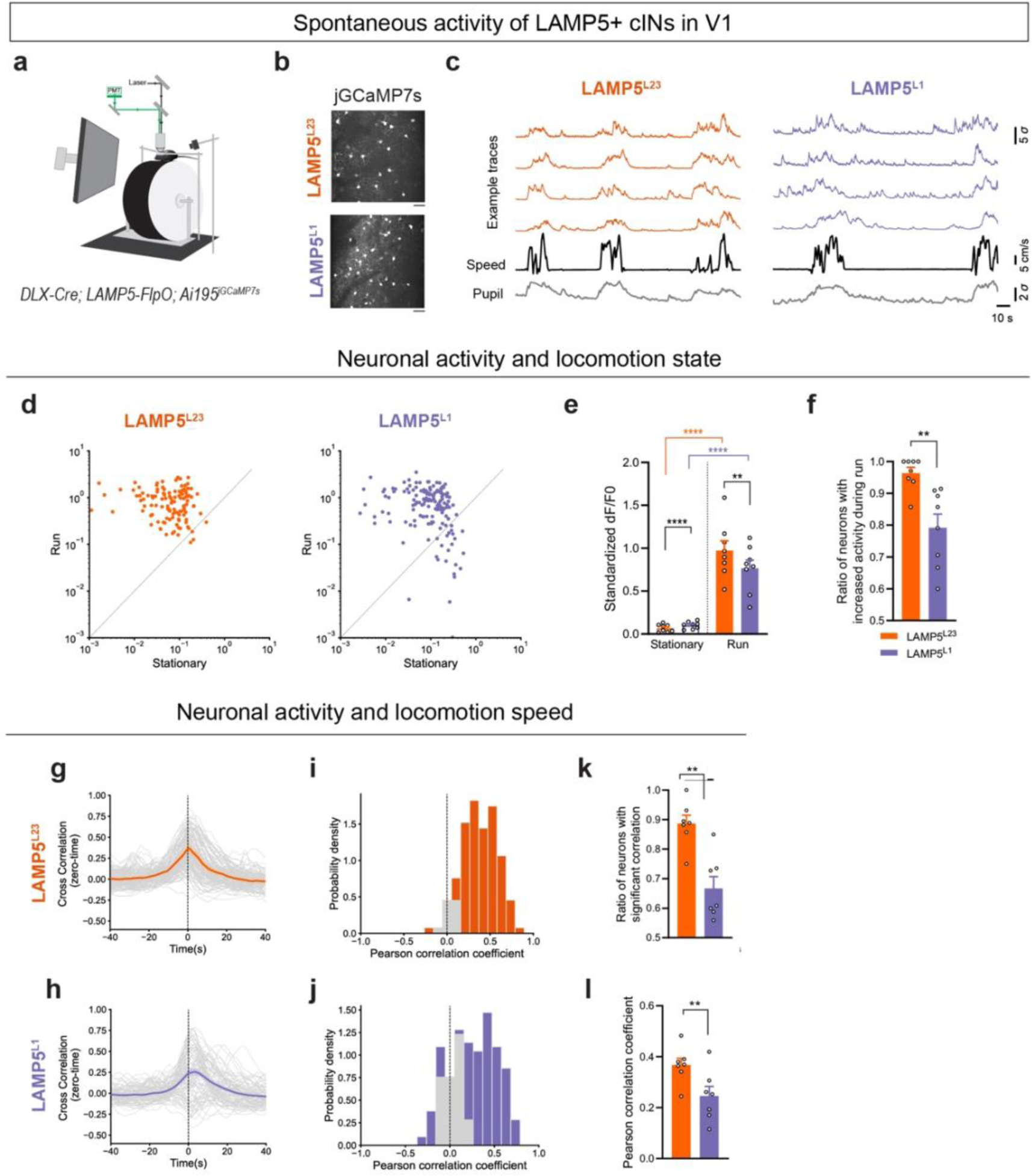
Spontaneous activity of LAMP5+ cINs is correlated with locomotion speed. (a) Illustration of the experimental setup during the spontaneous activity recordings. The animal was presented with a gray screen (mean luminance) while the GCaMP signal was imaged with two-photon microscopy. (b) A representative mean projection image from a two-photon recording session in L2/3 (upper) or L1 (bottom) LAMP5+ cINs in V1 of *DLX-Cre; LAMP5-FlpO; Ai195^jGCaMP7s^*. (c) Example traces from spontaneous activity recordings in LAMP5^L23^ (left, in orange) and LAMP5^L1^ (right, in purple). The top four traces represent the standardized dF/F0 activity of four randomly chosen neurons during a randomly selected time interval. The activity traces shown here are smoothed for representation. The black trace indicates the animal’s locomotion speed (in cm/s), and the gray trace shows the z-scored pupil size, both measured concurrently. (d) Scatter plot showing the mean standardized dF/F0 during stationary and running periods for LAMP5^L23^ (left, in orange) and LAMP5^L1^ (right, in purple). Each dot represents a neuron. (e) Bar plot showing the mean standardized dF/F0 during stationary (white bar) and run (color filled bar) periods for LAMP5^L23^ (orange) and LAMP5^L1^ (purple). (f) Bar plot showing ratio of neurons with increased activity during running periods. (g-h) Zero-time cross-correlation analysis of neuronal activity between (g) LAMP5^L23^ (orange) or (h) LAMP5^L1^ (purple), and locomotion speed. (i-j) Histogram of Pearson’s correlation coefficient (same as cross-correlation value at zero-time) of (i) LAMP5^L23^ (orange) or (j) LAMP5^L1^ (purple) activity and locomotion speed. Gray bars represent pairs with no significant correlation and colored bars represent pairs with significant correlation, determined by comparing against shuffled time series for each pair for 1000 times (refer to Methods). (k) Bar plot showing ratio of neurons with significant correlation between the spontaneous activity and locomotion speed for LAMP5^L23^ (orange) or LAMP5^L1^ (purple). (l) Bar plot showing Pearson’s correlation coefficient of the spontaneous activity and locomotion speed for LAMP5^L23^ (orange) or LAMP5^L1^ (purple). Error bar represents SEM. Each dot in the bar plots represents data from an individual mouse. Mann-Whitney test (f,k), mixed linear model regression (e) and hierarchical bootstrap (l) were used for testing statistical significance. See supplementary data 1 - Table 4 for statistics.

Notably, both LAMP5^L23^ and LAMP5^L1^ showed significantly increased spontaneous activities while the mice were running **(Figure 4d-f)**. We then investigated the temporal relationship between the neural activity of LAMP5+ cINs and locomotion speed of the mice by performing the zero-time cross-correlation analysis. We found that the spontaneous activity of both LAMP5^L23^ and LAMP5^L1^ were highly correlated with the locomotion speed **(Figure 4g-l)**. Together, these results suggested that LAMP5+ cINs were recruited during active behavioral states.

Moreover, a close comparison between the spontaneous activity of LAMP5^L23^ and LAMP5^L1^ revealed that the activity of LAMP5^L1^ exhibited greater heterogeneity. While nearly all LAMP5^L23^ showed increased activity during running compared to the stationary period, only 80% of LAMP5^L1^ exhibited a similar increase (**Figure 4f)**. Additionally, the zero-time cross-correlation analysis between neuronal activity and locomotion speed showed that LAMP5^L23^ exhibited a higher proportion of neurons that had significant correlation (p < 0.05 in the shuffle test, **Figure 4k)** and higher Pearson’s correlation coefficients (same as the correlation value at zero-time, **Figure 4l)**, indicating a stronger correlation between LAMP5^L23^ activity and locomotion speed. These results suggested that although both LAMP5^L23^ and LAMP5^L1^ activities are highly correlated with locomotion speed, LAMP5^L23^ activity were more homogeneous and better correlated with locomotion in time.

We also examined the synchronization of activity among LAMP5+ cINs **(Extended Data 5a,b)**, and found that the neuronal pairs in LAMP5^L23^ exhibited a higher proportion of significant synchronization **(Extended Data 5c)** and higher Pearson’s correlation coefficients **(Extended Data 5d),** compared to LAMP5^L1^. Most pairs in LAMP5^L23^ showed positive correlation, while a small percentage of LAMP5^L1^ pairs exhibited negative correlation within the population **(Extended Data 5b)**. Together, these results suggested that LAMP5+ cINs may exhibit some degree of lateral inhibition within L1, which is less prevalent in L2/3^16,31^.

Finally, to further validate these results, we repeated these experiments with mice viewing a dark screen **(Extended Data 5e-k)**, yielding similar results. Secondly, we replicated the experiments with an alternative strategy *JAM2-Cre; VIP-FlpO; Ai65F(RCF-tdT)*, and injected AAV9.Dlx.DIO.jGCaMP8m into V1. We recorded spontaneous activity from JAM2+ neurons in L2/3 (JAM2^L23^) and L1 (JAM2^L1^), manually excluding off-targeted VIP+ cINs in L2/3 of *JAM2-Cre* from the analysis **(Extended Data 2m, Extended Data 6a)**. The results from these experiments were consistent with those observed in *DLX-Cre; LAMP5-FlpO; Ai195* **(Extended Data 6)**. Lastly, we performed parallel experiments investigating the spontaneous activity of VIP+ and SST+ cINs in relation to locomotion. Consistent with previous reports^38–40^, these results showed a strong correlation between VIP+ cIN activity and locomotion speed, while the activity of only a subset of SST+ cINs was found to be modulated by locomotion **(Extended Data 7)**.

In conclusion, our experiments reveal that the spontaneous activities of LAMP5+ cINs increase during locomotion, strongly correlate with locomotion speed and are highly synchronized. In conjunction with our previous findings **(Figure 2)**, this suggests that LAMP5+ cINs hyperpolarize the baseline activity of PYN^L23^ and may contribute to sparse cortical activity in L2/3, especially during active behavioral states. Furthermore, we demonstrate that LAMP5^L23^ exhibit a higher degree of homogeneity in their responses and correlation with locomotion speed, whereas LAMP5^L1^ show more heterogeneity. This may arise from some degree of lateral inhibition within L1^16,31^, attributed to higher density of these neurons in L1, their elongated axonal morphology, combined with the non-selective inhibition properties of LAMP5+ cINs.

### Visual Responses and Their Modulation by Locomotion are Distinct in LAMP5^L23^ and LAMP5^L1^

Having established the important role played by LAMP5+ cINs in regulating cortical activity, we shifted our focus to understanding their recruitment in V1 *in vivo* during visual stimulation. Specifically, we aimed to investigate whether these neurons respond differently to visual stimuli based on their laminar position, and how these responses may change under different behavioral states.

We first tested the orientation tuning of LAMP5+ cINs by presenting *DLX-Cre; LAMP5-FlpO; Ai195* mice with full-field moving gratings of 12 different orientations **(Extended Data 8a)**. Similar to other cINs, LAMP5+ cINs showed a broad orientation tuning curve, as indicated by low gOSI **(Extended Data 8b-c)**. Additionally, we found that both LAMP5^L23^ and LAMP5^L1^ showed minor visual responses during the stationary period, while locomotion significantly increased their visual responses **(Extended Data 8d)**.

Prior research has indicated the visual responses of cINs can be modulated by stimulus contrast. For example, SST+ cINs exhibit stronger responses to high-contrast stimuli, whereas VIP+ cINs are more responsive to low-contrast visual gratings in V1, with PYN^L23^ displaying mixed preferences^41^ (we confirmed these results in **Figure 5b-c, Extended Data 9a-c)**. Here, we investigated how stimulus contrast influences the visual responses of LAMP5+ cINs. To this end, we conducted two-photon calcium imaging in V1 of awake *DLX-Cre; LAMP5-FlpO; Ai195* mice, and presented them with moving gratings of different contrasts and orientations in each trial. Each trial began with a 1s gray screen baseline, followed by 2s of moving gratings, with “blank” trials maintaining a gray screen (equal to 0% contrast) throughout the 3s duration **(Figure 5a)**.

**Figure 5.**
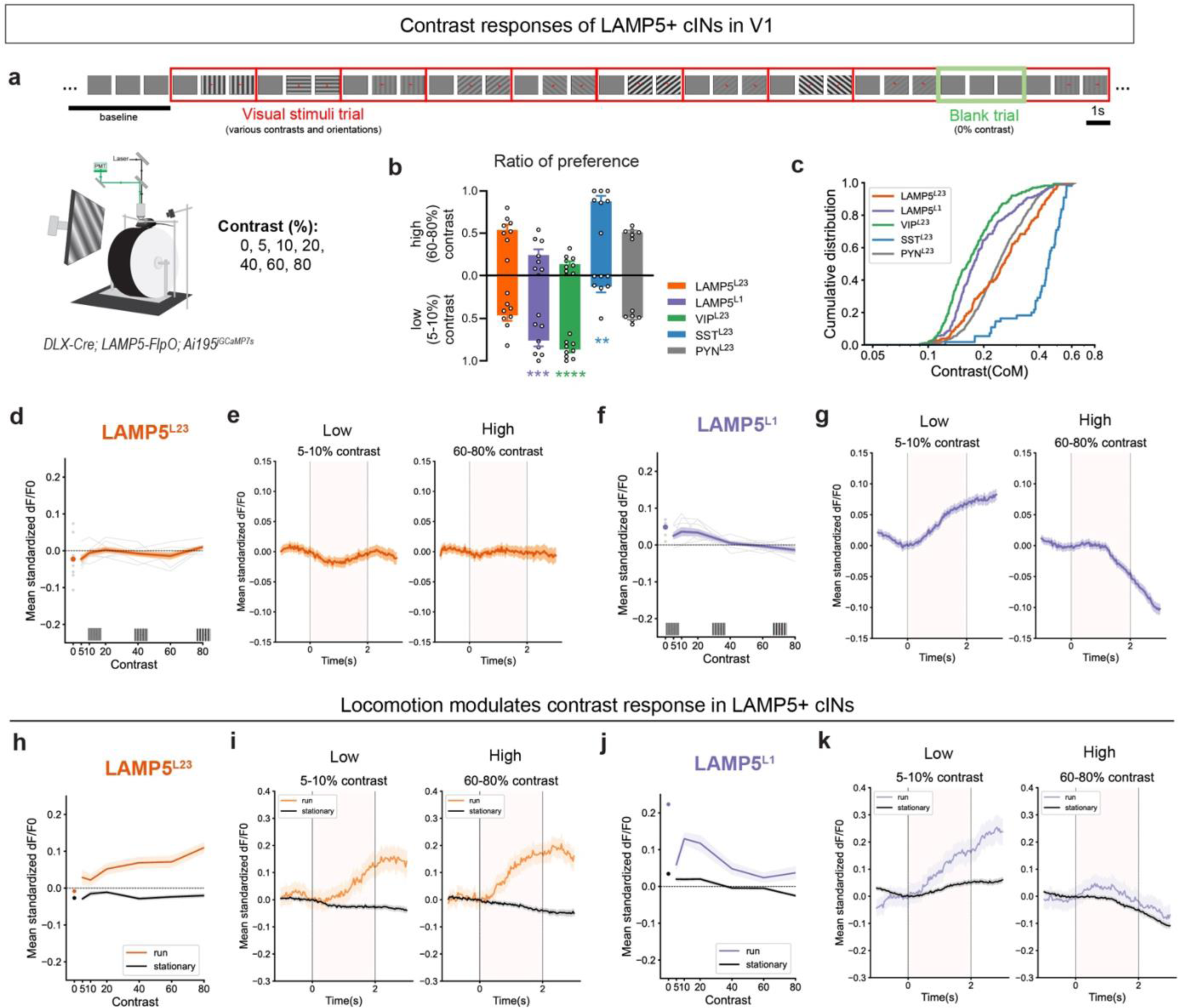
Visual responses and their modulation by locomotion are distinct in LAMP5^L23^ and LAMP5^L1^. (a) Illustration of the experimental trial design. After a baseline of 10 s with the gray screen, mice were presented with random 3s visual stimuli trials which each had 2s of full-screen moving gratings with variable contrast and orientations in each trial. 1s of inter-stimulus-interval was used with the gray screen. Six contrast levels (5%, 10%, 20%, 40%, 60% and 80%) and eight orientations (every 45°) were included in the task and each combination was randomly repeated for 15 trials. In addition, we included blank trials (0% contrast) randomly. Each blank trial consisted of 3s of gray screen. (b) Ratio of high (60-80%) or low (5-10%) contrast preferring neurons in LAMP5^L23^ (orange), LAMP5^L1^ (purple), VIP^L23^ (green), SST^L23^ (blue) and PYN^L23^ (gray). Each dot represents data from an individual animal. VIP^L23^/SST^L23^ results were from *Germline-Cre; VIP-FlpO/SST-FlpO; Ai195* mice (*Ai195* is an intersectional reporter and this converts them into a Flp-reporter). PYN^L23^ results were from TOX-control animals with AAV.GCaMP6f. (c) Cumulative ratio of the center of mass in contrast response for each neuron in LAMP5^L23^ (orange), LAMP5^L1^ (purple), VIP^L23^ (green), SST^L23^ (blue) and PYN^L23^ (gray). (d) Visual responses at various contrast levels in LAMP5^L23^ population. (e) Averaged response trace in (left) low (5-10%) or (right) high (60-80%) contrasts for LAMP5^L23^. (f) Visual responses at various contrast levels in LAMP5^L1^ population. (g) Averaged response trace in (left) low (5-10%) or (right) high (60-80%) contrasts for LAMP5^L1^. (h) Visual responses at various contrast levels in LAMP5^L23^ population during running (orange) or stationary (black) trials. (i) Averaged response trace in (left) low (5-10%) or (right) high (60-80%) contrasts for LAMP5^L23^ during running (orange) or stationary (black) trials. (j) Visual responses at various contrast levels in LAMP5^L1^ population during running (purple) or stationary (black) trials. (k) Averaged response trace in (left) low (5-10%) or (right) high (60-80%) contrasts for LAMP5^L1^ during running (purple) or stationary (black) trials. In (d,f,h,j), each gray line and dot represent averaged data from an animal, while color line represents mean and SEM from all neurons. Responses at 0% contrast (‘blank trial’) were indicated by dot. In (e,g,i,k), moving gratings were presented between 0-2 s indicated by dashed gray vertical lines. The color lines represent mean and SEM from all animals. Mann Whitney test (b) and mixed linear model regression (d,f,h,j) were u sed for testing statistical significance. See supplementary data 1 - Table 5 for statistics.

Results from these experiments showed that LAMP5^L23^ responded similarly regardless of contrast levels **(Figure 5b-e)**, while LAMP5^L1^ showed a preference for lower contrast levels **(Figure 5b,c,f,g)**. However, further analysis revealed that locomotion significantly increased LAMP5^L23^ visual responses across all contrast levels, with their visual responses showing a preference for higher contrasts during running trials **(Figure 5h,i, Extended Data 9e)**. Conversely, LAMP5^L1^ continued to favor low-contrast gratings, with enhanced visual responses during locomotion **(Figure 5j,k, Extended Data 9e)**. Intriguingly, LAMP5^L1^ showed positive responses during “blank” trials **(Figure 5f,j, Extended Data 9d)**, a finding warranting further investigation. To further support these findings, we confirmed these results using *JAM2-Cre*, *SST-Cre* or *VIP-Cre* with AAV9.Dlx.DIO.jGCaMP8m in V1 **(Extended Data 9f-q)**.

Overall, our findings suggest that LAMP5^L23^ exhibit significant visual responses and a preference for higher contrast visual stimulus, but only during active behavioral states. This suggests that LAMP5^L23^ may require a combination of visual input and state-modulatory input to exhibit responsiveness to visual stimuli. Additionally, we found that LAMP5^L1^ displayed greater responsiveness to lower contrast levels in both stationary and running states, with enhanced responses during locomotion. This property is similar to VIP^L23^, suggesting that LAMP5^L1^ and VIP^L23^ may share similar input connectivities involved in contrast modulation^21,28,30,42–45^.

### Layer-dependent Circuit Connectivity of LAMP5+ cINs

The variations in contrast modulation of visual responses between LAMP5^L1^ and LAMP5^L23^ raises the possibility that these neurons are activated by different combinations of circuit inputs within a specific context. We hypothesized that the location of their cell bodies and the morphology of their short dendrites might restrict them to being driven by different inputs.

To map the local inhibitory inputs from other cINs to LAMP5+ cINs, we conducted optogenetics-assisted circuit mapping with slice electrophysiology. Channelrhodopsin was specifically expressed in PV+, SST+ or VIP+ cINs using an intersectional *Ai80(CatCh-EYFP)* reporter mouse line crossed to *JAM2-Cre; SST-FlpO/VIP-FlpO/PV-FlpO* alleles (see Method - Mouse). LAMP5+ cINs were labeled via stereotaxic injection of AAV9.Dlx.DIO.dTomato into V1. Inhibitory postsynaptic currents (IPSCs) were recorded in LAMP5+ cINs by voltage clamp in the presence of tetrodotoxin (TTX) and 4-aminopyridine (4-AP) in postnatal (P) 38-42 mice **(Extended Data 10a)**. Our findings revealed that SST+ cINs inhibit LAMP5+ cINs across all six cortical layers **(Extended Data 10b)**. Additionally, we observed that PV+ cINs inhibit all LAMP5^L2-6^ but not LAMP5^L1^ **(Extended Data 10c)**, likely because PV+ cINs lack axonal extensions into L1^12^. Furthermore, VIP+ cINs to LAMP5+ cINs connection was generally weak across all cortical layers **(Extended Data 10d)**.

We next examined whether LAMP5^L23^ and LAMP5^L1^ are driven by different combinations of excitatory inputs based on their layer location. To do so, we explored the afferent connectivity of LAMP5+ cINs in V1 using retrograde monosynaptic rabies tracing. AAV helpers and the delta-G pseudorabies virus RVdG-mCherry were administered into V1 of adult *JAM2-Cre* mice, and brain tissue was subsequently processed after 13 days **(Figure 6a)**. The starter cells were sampled in all layers of V1 **(Figure 6b)**, although biased to different layers in each experiment **(Figure 6e)**. Monosynaptic inputs to these neurons were analyzed by aligning the sectioned brain tissues with the Allen CCFv3 atlas and calculating the ratio of rabies traced (mCherry+) cells found in a defined anatomical region to the total count of traced cells.

**Figure 6.**
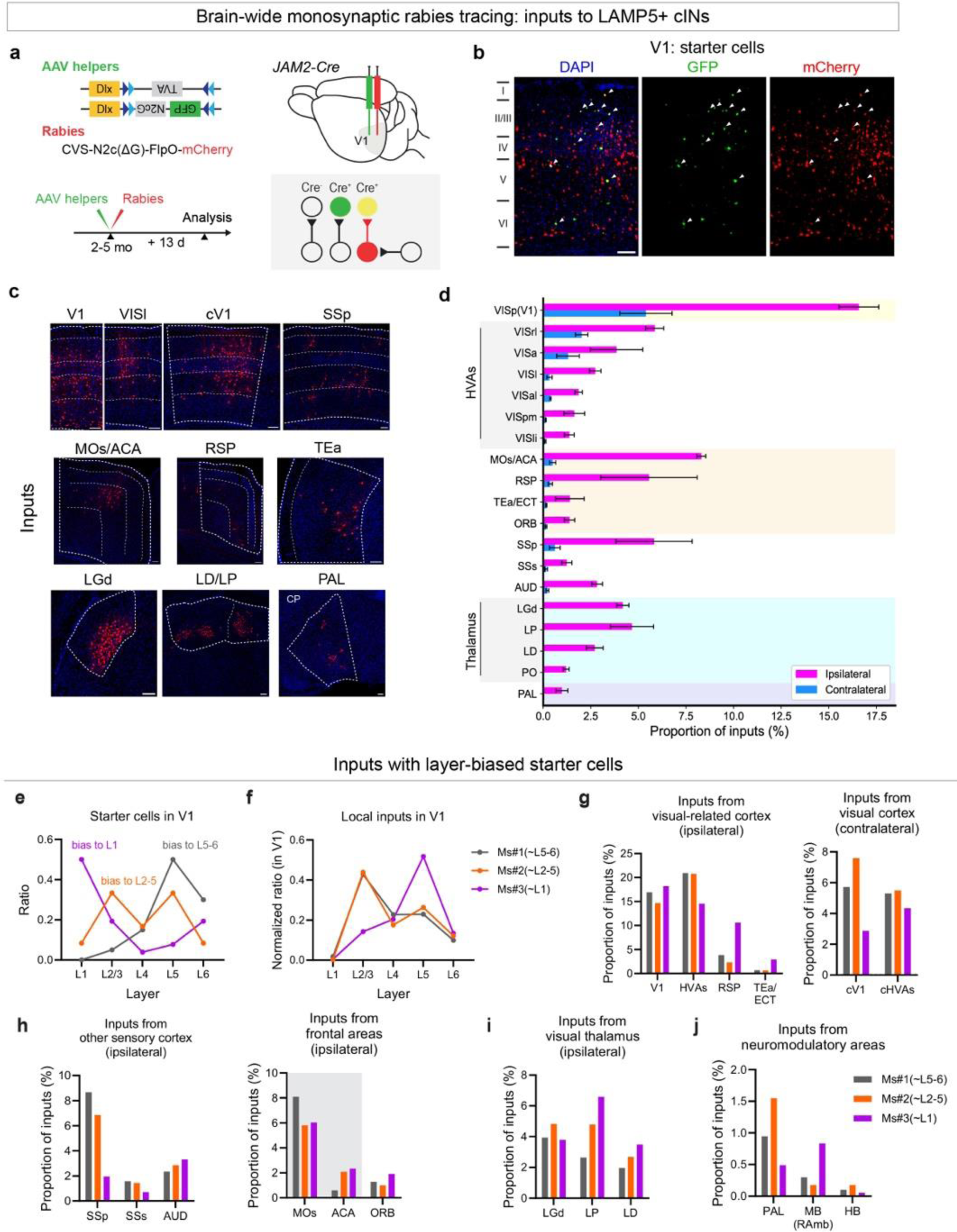
Brain-wide monosynaptic inputs to LAMP5+ cINs in V1 reveal layer-dependent circuit connectivity of LAMP5+ cINs. (a) Experimental design of rabies retrograde tracing from LAMP5+ cINs in V1. Helper AAVs, AAV1.Dlx.DIO.TVA and AAV1.DLX.DIO.GFP.N2cG (green), were co-injected with EnVA-pseudotyped CVS-N2c(ΔG)-FlpO-mCherry rabies virus (red) into V1 of *JAM2-Cre* mice. Rabies tracing patterns were analyzed 13 days post-infection. GFP+ cells represent N2cG protein expression, while mCherry+ cells indicate presynaptically traced neurons. Cells positive for both GFP and mCherry were identified as starter cells. (b) Example images showing starter cells in V1 (write arrow). Scale bar = 100 μm. (c) Example images showing brain regions with significant inputs. Scale bar = 100 μm. (d) Presynaptic inputs to LAMP5+ cINs in V1 were quantified as the percentage of rabies traced cells in each region out of the total number of cells labeled in the brain. Regions with >1% of inputs are included in the plot. Inputs from the ipsilateral side are colored in magenta, while those from the contralateral side are in blue. (e) Starter cell layer distribution for 3 experimental repeats, Ms#1 (gray) had starter cells biased to deep layers L5-6, Ms#2 (orange) had starter cells biased to middle layers L2-5, while Ms#3 (purple) had starter cells biased to L1. (f) Local inputs across cortical layers normalized to total local inputs (mCherry+, GFP-cells in V1) in V1 for 3 experimental repeats. (g-j) Inputs from ipsilateral visual areas, contralateral visual areas, ipsilateral non-visual sensory areas, ipsilateral motor and frontal areas, ipsilateral visual thalamus, and putative neuromodulatory areas, such as PAL (putative cholinergic projection neurons in pallidum), MB (RAmb) (putative serotonergic neurons) and HB (putative noradrenergic neurons) for 3 experimental repeats. The error bars represent SEM. Data was collected from N=3 animals. Abbreviations: ACA: anterior cingulate area, AUD: auditory areas, cV1: contralateral primary visual area, cHVAs: contralateral higher-order visual areas, ECT: ectorhinal area, HB: hindbrain, HVAs: higher-order visual areas, LD: lateral dorsal nucleus of thalamus, LGd: dorsal part of the lateral geniculate complex, LP: lateral posterior nucleus of the thalamus, MB: midbrain, MOs: secondary motor area, ORB: orbital area, PAL: pallidum, PO: posterior complex of the thalamus, RAmb: midbrain raphe nuclei, RSP: retrosplenial area, SSs: supplemental somatosensory area, SSp: primary somatosensory area, TEa: temporal association area, VISa: anterior area, VISal: anterolateral visual area, VISl: lateral visual area, VISli: laterointermediate area, VISpm: posteromedial visual area, VISp(V1): primary visual area, VISrl: rostrolateral visual area.

Our findings indicate that LAMP5+ cINs receive a wide range of both local V1 and long-range excitatory inputs. Major inputs to these cells in V1 originate from higher-order visual areas (HVAs), the contralateral V1 (cV1), the retrosplenial area (RSP), the secondary motor area (MOs)/anterior cingulate area (ACA), the primary somatosensory area (SSp), and auditory areas (AUD). Inputs from the visual thalamus, originate from the dorsal part of the lateral geniculate complex (LGd), the lateral posterior nucleus of the thalamus (LP), and the lateral dorsal nucleus of the thalamus (LD), as well as inputs from neuromodulatory areas like the basal forebrain (BF) were also identified **(Figure 6c-d)**.

Furthermore, the starter cells within each of the three experiments exhibited biases towards L1, L2-5, or L5-6 populations **(Figure 6e)**. This allowed us to compare differences in the afferents targeting LAMP5+ cINs in distinct laminae across experiments. We first examined local V1 inputs. L1-biased starter cells receive significant inputs from L5 (>50% of all local inputs, **Figure 6f)**, consistent with the results from the prior study on NDNF+ L1 cINs in V1^42^. In contrast, when the starter cells were biased towards L2-5 or L5-6, >40% of local inputs originated in L2/3, followed by ∼20% of inputs originating from either L4 or L5 **(Figure 6f)**. LAMP5^L1^ also showed differences in their long-range inputs compared to those in other cortical layers. L1-biased starter cells received a higher ratio of inputs from areas such as RSP, the temporal association area (TEa) **(Figure 6g)**, the orbital area (ORB) **(Figure 6h)**, and the higher-order visual thalamus - LP and LD **(Figure 6i)**. These results aligned with the preferential innervation of L1 in V1 from these regions^43,46,47^. Notably, with L1-biased starter cells, we found a higher ratio of inputs from the midbrain (MB) dorsal raphe nucleus, which may houses 5-hydroxytryptamine (5-HT) serotonergic neurons, but a lower ratio of inputs from the basal forebrain pallidum (PAL), where cholinergic projection neurons may reside **(Figure 6j)**.

In summary, monosynaptic tracing from LAMP5+ cINs showed that LAMP5^L1^ and LAMP5^L23^ receive distinct local and long-range inputs, likely contributing to their distinct visual response properties, such as the contrast preference. Specifically, we propose that LAMP5^L1^ primarily receive top-down inputs targeting L1, along with local excitatory inputs from L5. In contrast, LAMP5^L23^ rely less on top-down inputs but are driven more by inputs that convey bottom-up sensory signals **(Extended Data 10e)**. Additionally, we identified local inhibitory inputs to LAMP5+ cINs in each layer. These findings underscore the variability in inputs to LAMP5+ cINs based on their laminar location, where their short dendrites restrict inputs to those in close proximity. Consequently, the source of inputs to LAMP5+ cINs is primarily determined by the axonal innervation that reaches their home layer. Overall, our results suggest that LAMP5+ cINs in different layers may be recruited differently in behavioral contexts, highlighting the importance of morphology and circuit connectivity in determining neuronal activity beyond transcriptomic types.

## DISCUSSION

Neurogliaform cells, which comprise the majority of LAMP5+ cINs and are characterized in cortical circuits by their distinctive “volume transmission” output properties, remain the least studied type of cINs. This is largely due to the lack of genetic tools for targeting them. In this study, we introduced two genetic targeting strategies that selectively target LAMP5+ cINs **(Figure 1b, Extended Data 2a,b)**, enabling us to explore their circuit and function *in vivo*. We found that silencing the output activity of LAMP5+ cINs leads to increased spontaneous activity in PYN^L23^, especially during active behavioral states **(Figure 2)**. In addition, the spontaneous activity of LAMP5+ cINs was found to be highly correlated with the active behavioral states **(Figure 4)**. These findings suggest that LAMP5+ cINs are likely involved in regulating the behavioral state-dependent baseline activity of PYN^L23^. Notably, LAMP5+ cINs in L1 versus L2/3 are differentially recruited during visual stimulation. We demonstrated that the input connectivity of LAMP5+ cINs is constrained by their specific laminar locations and short dendritic morphology **(Figure 6, Extended Data 10e)**, likely contributing to the observed layer-specific preferences in contrast modulation of visual responses **(Figure 5)**. Given their non-selective output properties and layer-restricted axonal arborizations, these differences in recruitment may play a crucial role in modulating distinct compartments of PYNs under different contextual conditions **(Extended Data 10f)**.

Using our genetic labeling strategy, we observed that LAMP5+ cINs are primarily distributed in L1-3, with fewer cells in L4 and L6, with the low densities being observed in L5. This observation is interesting, considering that spontaneous and evoked cortical activity tends to be sparse in L2/3 but dense in L5 within sensory cortical areas, which matches the layer distribution of LAMP5+ cINs in L2-5. It is believed that the sparsity of PYN activity in superficial layers enhances the robustness and reliability of sensory encoding, thereby improving perception^48–52^. Our findings, that silencing LAMP5+ cINs results in increased spontaneous activity and synchrony in PYN^L23^, indicate that LAMP5+ cINs can be one of the key players that contribute to the observed sparsity of cortical activity in supragranular layers.

Although LAMP5+ cINs exhibit a continuum in their transcriptomic profiles and electrophysiological properties within the cardinal type population^12,53^, we believe that their layer-dependent input connectivity is crucial for shaping discrete activity patterns for LAMP5+ cINs in different cortical layers. For example, we showed that LAMP5^L1^ predominantly responds to low-contrast visual stimuli, similar to VIP+ cINs^41^, while LAMP5^L23^ preferentially responds to higher-contrast stimuli during locomotion states. This functional distinction likely results from the different combinations of inputs activating LAMP5^L1^ or LAMP5^L23^ during visual stimuli in different behavioral states. Brain-wide monosynaptic input analysis suggests that LAMP5+ cINs receive different inputs depending on their layer locations, primarily due to the limited reach of their short dendrites, confining them to receive inputs from nearby axons around their cell body locations. These input differences likely explain the distinct engagement of LAMP5^L23^ and LAMP5^L1^ in response to various visual stimuli. For example, fewer local excitatory inputs, more long-range cortical feedback inputs, and local inhibitory inputs from SST+ cINs may collectively contribute to the similar preference for low-contrast stimuli observed in LAMP5^L1^ and VIP+ cINs. In contrast, inputs that convey the bottom-up sensory signals, together with neuromodulatory signals in active behavioral states may drive the high contrast-preferring visual activity of LAMP5^L23^ during locomotion **(Extended Data 10e)**.

With regards to their outputs, LAMP5+ cINs also exhibit highly ramified thin axons primarily confined to their cell body layer and adjacent layers^12^. Although both use slow and prolonged “volume transmission” of GABA to mediate inhibition, LAMP5^L1^ and LAMP5^L2-6^ may regulate cortical activity differently due to their distinct organization within the cortex **(Extended Data 10f)**^22^. LAMP5^L1^, by modulating the distal dendrites of all PYNs in L1, can modulate the activity of an entire cortical column^30,54^. In contrast, LAMP5^L23^ are positioned adjacent to the somatic regions of PYN^L23^, and can mediate laminar-specific regulation of activity within L2/3 more directly. Furthermore, the elongated morphology of LAMP5^L1^ and their denser distribution in L1 may contribute to lateral inhibition among LAMP5^L1^, resulting in the inhibition of a subset of LAMP5^L1^ during locomotion in our spontaneous activity recordings. In contrast, LAMP5^L23^ are likely organized in a manner that minimizes interference among themselves, illustrating the morphological adaptation of cINs in different layers to regulate layer-dependent circuit activities. Despite these differences, most LAMP5+ cINs can inhibit all other cINs within their axonal target range or couple the membrane potentials of those they are electrically gap-junctioned. This includes not only other LAMP5+ cINs but also various other cIN types^22–24^. These two competing aspects of LAMP5+ cINs likely contribute to the complexity of inhibitory network dynamics.

Taking into account their layer-dependent recruitment and the morphological constraints on their outputs, we believe that LAMP5^L23^ and LAMP5^L1^ can be engaged in different behavioral contexts, each shaping cortical activity in L2/3 in distinct ways. For example, LAMP5^L23^ integrate a variety of inputs to monitor the local excitation levels in L2/3, while LAMP5^L1^ may play a more active role in regulating dendritic excitation in L1 in response to higher-order feedback inputs. Consequently, the recruitment of LAMP5^L23^ versus LAMP5^L1^ may not simply reflect a layer-based segregation of function but also represent an adaptive strategy to optimize cortical processing and output in response to dynamic environmental cues.

Our findings, along with those of others, have indicated that both LAMP5+ cINs and VIP+ cINs exhibit spontaneous activity highly correlated with locomotion speed in awake, behaving mice^38,39^. This observation is particularly interesting given the distinct inhibitory mechanisms of these two cIN populations. While LAMP5+ cINs are capable of broadly inhibiting both excitatory and inhibitory neurons, VIP+ cINs primarily function through disinhibition of PYNs on their distal dendrites by inhibiting SST+ cINs^55,56^. These results indicate that the state-dependent modulation of cortical activity is a consequence of intricate cortical network dynamics. Notably, LAMP5+ cINs exhibit high expression levels of excitatory receptors such as adrenergic receptors (ADRA1A, ADRA1B), muscarinic cholinergic receptor (CHRM1), and nicotinic cholinergic receptors (CHRNA4, CHRNA7, CHRNB2). These receptor expressions may contribute to the observed state-dependent activity in LAMP5+ cINs, and may mediate their responsiveness to neuromodulatory signals^10,27,38,57^.

In this study, we did not distinguish the ∼20% LAMP5^L1^ that are CHRNA7+. These neurons are transcriptomically distinguishable from the rest of the LAMP5+ cINs belonging to neurogliaform cells, and they do not share similar properties as neurogliaform cells discussed above. Their axons can extend into deep cortical layers^16^. The circuit connectivity and function of these neurons remain unclear. While we found that the activity pattern in LAMP5^L1^ is more heterogeneous than that seen in LAMP5^L23^, this is unlikely to be due to CHRNA7+ LAMP5^L1^, as these neurons have been reported to be more active during running state. Instead this heterogeneity is more likely to be a result of the lateral inhibition of neurogliaform cells in L1^16,31,53^.

The unique inhibition from LAMP5+ cINs may have important clinical implications. Interestingly, we found that LAMP5+ cINs, particularly neurogliaform cells, exhibit the highest level of GAD2 (glutamic acid decarboxylase 65-kd isoform) expression not only in mice but also in humans^27^. GAD2 is primarily present in presynaptic terminals and plays a crucial role in GABA synthesis for vesicle release^58^. Remarkably, dysregulation of GAD2 has been implicated in neurological disorders such as epilepsy. Studies have shown that both GAD1 (glutamic acid decarboxylase 67-kd isoform) and GAD2 expression levels in LAMP5+ cINs are reduced in patients with temporal lobe epilepsy^26^, suggesting a potential role of LAMP5+ cIN dysregulation in hypoinhibition underlying epileptogenesis. Furthermore, LAMP5+ cINs in these patients also exhibit decreased levels of SV2C (synaptic vesicle glycoprotein 2C)^59^, a conserved marker gene for neurogliaform cells within LAMP5+ cINs. This observation suggests that SV2C may play crucial roles in regulating vesicles in neurogliaform cells^60–62^.

In conclusion, our results highlight the functional uniqueness of LAMP5+ cINs and underscore their importance as a key population within the cortical inhibitory network. Furthermore, we advocate for the consideration of both transcriptomic cell types and circuit connectivity in parallel when categorizing cell type diversity in cortical circuits. Integrating these factors can provide a more comprehensive understanding of the functional roles of different cell populations and their contributions to cortical information processing.

## METHODS

### Mouse

All experimental procedures were approved by and in accordance with Harvard Medical School Institutional Animal Care and Use Committee (IACUC) protocol number IS00001269. Animals were group housed and maintained under standard, temperature-controlled laboratory conditions. Mice were kept on a 12:12 light/dark cycle and received water and food ad libitum. Mice used in the *in vivo* imaging experiments were kept on a reversed light cycle. Both female and male animals were used indiscriminately for all experiments. Though a systematic analysis was not performed to assess whether there are sex-related differences, no obvious pattern was observed. Data collection and analysis were not performed blind to the conditions of the experiments. Mouse lines used in this study: *DLX-Cre* (RRID:IMSR_JAX:008199), *LAMP5-FlpO* (RRID:IMSR_JAX:037340, a gift from Dr. John Ngai and Dr. David A. Stafford, University of California, Berkeley), *Ai65(RCFL-tdT)* (RRID:IMSR_JAX:021875, the Jackson Laboratory), *JAM2-Cre* (RRID:IMSR_JAX:031612, the Jackson Laboratory), *Ai65F(RCF-tdT)* (RRID:IMSR_JAX:032864, the Jackson Laboratory), *RC:PFTox^37^* (a gift from Dr. Susan Dymecki and Dr. David Ginty, Harvard Medical School), *Ai195* (RRID:IMSR_JAX:034112, a gift from Dr. Tanya L. Daigle, Dr. Bosiljka Tasic and Dr. Hongkui Zeng, Allen Brain Institute), *VIP-FlpO* (RRID:IMSR_JAX:028578, the Jackson Laboratory), *SST-FlpO* (RRID:IMSR_JAX:031629, the Jackson Laboratory), *PV-FlpO* (RRID:IMSR_JAX:022730, the Jackson Laboratory). All mice were maintained in house on a C57BL/6J (RRID:IMSR_JAX:000664, the Jackson Laboratory) background.

### Genetic strategy with *JAM2-Cre*

We chose *JAM2-Cre* because JAM2 is a conserved marker gene for neurogliaform cells **(Extended Data 1a1)**. However, JAM2 is also expressed in germ cells, with its expression becoming restricted to LAMP5+ cINs only in adults. This limits the use of this strategy to adult ages. In our experiments, *JAM2-Cre* served not only as a targeting tool for LAMP5+ cINs in adults using AAV, but also as a *Germline-Cre* to convert intersectional mouse lines (*RC:PFtox, Ai195,* or *Ai80*) into Flp-dependent mouse lines. Through DNA genotyping, we observed that offspring bred from *JAM2-Cre* mice and intersectional reporters became a Flp-dependent reporter. We found that JAM2 RNA has been detected in germ cells, including sperm^63,64^, suggesting that Cre activity in *JAM2-Cre* mice is activated in germ cells. Therefore, *JAM2-Cre* was also used as a *Germline-Cre* in our experiments.

### Plasmid construction

pAAV.Dlx.DIO.jGcamp8m plasmid was constructed by first flipping the MCS with BcuI with the backbone pAAV-VTKD2 (Addgene #170847)^65^, then inserting the jGCaMP8m^66^ fragment from pGP-AAV-syn-jGCaMP8m-WPRE (Addgene #162375) digested by EcoRI and HindIII. pAAV.Dlx.DIO.dTom has been donated to Addgene (Plasmid #83894) previously^34^. pAAV.Dlx.DIO.TVA was constructed by inserting the hDlx promoter fragment from pAAV-VTKD2 (Addgene #170847) into the backbone pAAV-EF1a-flex-TVA (Addgene #69618), digested by EcoRI and HindIII. pAAV.Dlx.DIO.GFP.N2cG was constructed by Gibson assembly the PCR fragment GFP-P2A-N2cG from pAAV-VTKS2-TVA-eGFP-N2cG (Addgene #175439) into the backbone pAAV-VTKD2 (Addgene #170847)^65^ digested by EcoRI and HindIII.

### Cell culture, transfection and AAV production

HEK293FT cells (Thermo Fisher Scientific, #R70007) were cultured in Dulbecco’s Modified Eagle’s medium with high glucose and pyruvate, GlutaMAX Supplement, 10% fetal bovine serum, penicillin (100 units/ml) and streptomycin (100 μg/ml). The cultures were incubated at 37 °C in a humidified atmosphere containing 5% CO2. For AAV production, HEK293FT cells were seeded on 15-cm dishes without antibiotics for close to 24 hours and co-transfected with the following plasmids using Polyethylenimine (100 μg/dish, Polysciences, #23966-1): pHGTI-Adeno1 helper (22 μg/dish), AAV9 helper (Addgene plasmid #112865, 9 μg/dish), and the AAV expression vector (12 μg/dish). 72 hours after transfection, transfected cells were harvested and lysed (150 mM NaCl, 20 mM Tris pH 8.0) by three freeze-thaw cycles and Benzonase treatment (375 U/dish; Sigma, #E1014) for 30 min at 37 °C. The supernatants were cleared by centrifugation at 4000 RPM for 20 min at 25 °C, then transferred to Iodixanol gradients (OptiPrep Density Gradient Medium, Sigma, #D1556) for ultracentrifugation (VTi50 rotor, Beckman Coulter) at 50,000 RPM for 1.5 hr at 16 °C. The 40% iodixanol fraction containing the AAVs was collected, underwent ultrafiltration with PBS in Amicon Ultra-15 Centrifugal Filter (15 ml, 100kDa MWCO, Millipore, #UFC910024) at 4000 RPM for 1 hr for 4 times, aliquoted and stored at −80 °C. The number of genomic viral copies was determined by qPCR using the following primers against the WPRE sequence: Fw: AGCTCCTTTCCGGGACTTTC and Rv: CACCACGGAATTGTCAGTGC.

### *In vivo* imaging: cranial window and virus injection surgery

For in vivo imaging experiments, surgeries were carried out in mice after they were 2 months old. 1 Rimadyl (2 mg/tablet, Bio-serv, #MD150-2) tablet per animal was placed in the cage 1 day prior to the surgery. Mice were anesthetized with isoflurane (5% for induction, 1-2% during surgery, in O^2^), mounted in a stereotaxic frame, and kept on a warm blanket (34 °C). The eyes were moistened with lubricant eye ointment (Systane). The scalp was disinfected with 10% Povidone-Iodine (PDI, #S41125) and a section of scalp was removed using microscissors. 0.3% hydrogen peroxide was applied on the skull to oxidize and facilitate removal of periosteal tissue with cotton tip swabs, and washed with sterile saline. The 3 mm cranial window was centered around 2.5 mm lateral to the midline (left hemisphere) and 1.5 mm anterior to the transverse sinus (or Lambda suture), skull was removed with 3 mm biopsy punch (Integra™ 3332) and micro knives (Fine Science Tools, 10315-12). Virus injections in V1 were performed using beveled glass micropipettes (Drummond, 3-000-203-G/X) with Nanoject III (Drummond). 300 nl of virus was injected into each of the 2-3 locations, at 3 depths (0.43, 0.38 and 0.33 mm) below the dura. AAV2/9.CaMKII.GCaMP6f.WPRE.SV40 was diluted to 6.7E+12 vector genomes per ml (vg/ml) with 1X PBS prior to injection for Tox experiments. AAV2/9.VTKD2.jGcamp8m was diluted to 1.0E+13 vg/ml with 1X PBS prior to injection to *JAM2^Cre^* mice. No virus injections were performed in the GCaMP reporter (Ai195) mice. The cranial window was kept moist with sterile saline during virus injection and sealed with two circular, pre-sanitized glass coverslips, 3 mm and 5 mm in diameter (Warner Instruments, #64-0700 and #64-0720), individually conjoined with optical adhesive (Norland, NOA 71). The 3 mm coverslip was laid over the pia surface within the cranial window. Tissue adhesive (3M Vetbond) was applied around the 3 mm coverslip. A custom-designed head plate was adhered over the glass window with Super Glue gel (Loctite) and C&B Metabond (Parkell, #171032) mixed with black powder paint. Mice were given Buprenorphine SR 0.5-1.0 mg/kg SC after the surgery and monitored for 5 days post-surgery.

### *In vivo* imaging: intrinsic imaging

Intrinsic optical signal imaging was conducted via the cranial window of head-fixed mice. Prior to imaging, mice received an intramuscular injection of 0.2mg/ml chlorprothixene (Sigma-Aldrich, Y0002088) at a dosage of 80µl per 20g of body weight, administered at least 15 minutes before the experiment. During imaging, additional 0.5-0.75% isoflurane anesthesia (in O^2^) was applied to mice, and the heat pad (34°C) was used to maintain body temperature. Illumination of the cortex was achieved using a white LED cold light source equipped with light guides and a filter slider (Leica, KL1600). For vessel imaging, a green filter (Schott, 258.304) was inserted, while intrinsic signals were recorded under a red filter (Schott, 258.303). Tandem lens systems were used, consisting of a Nikkor 85mm F2.0 AI-S (Nikon) as the top lens and an AF Nikkor 50mm f/1.8D (Nikon) as the bottom lens. Additionally, green (Edmund, #87-801) and red (Edmund, #88-018) emission filters were optionally used during vessel imaging and intrinsic signal recording, respectively, although they were not essential. Data acquisition was performed using a USB camera (FLIR, BFS-U3-19S4M-C)^67^ and SpinView 2.5.0.80 software. The image format for acquisition was configured to 12 bits, with a binning setting of 2 x 2 pixels, resulting in recordings at a resolution of 808 x 620 pixels with a pixel size 5.3 x 5.3µm. Vessel images were captured at the beginning of the experiment, then the lenses were focused around 0.5mm below the surface. Visual stimuli were generated using custom MATLAB (Mathworks) scripts with Psychtoolbox-3^68,69^ and presented at a distance of 16 cm from the right eye on a gamma-corrected LED-backlit LCD monitor (Dell P2317H, 509mm X 286mm) with a mean luminance of 20 cd/m². Each trial consisted of a 2s baseline period (full screen RGB black), followed by 1s of visual stimulation and 20s post-stimulus interval for recovery (full screen RGB black). The visual stimulation comprised four-direction full-field drifting gratings (at 0°, 45°, 90°, and 135°, each lasting 250ms) with 100% contrast, a temporal frequency of 2Hz, and a spatial frequency of 0.04 cycles per degree (cpd)^70^. A total of 20-25 trials were repeated, with a frame rate set at 58.89Hz, and 1000 frames were recorded for each repetition. A small block of bright pixels was positioned at the bottom left corner of the screen at the beginning of each trial. This signal was captured by a photodiode (Thorlabs, SM05PD1A) affixed to a duplicate secondary screen, processed by a custom current-to-voltage converter to generate a trigger for initiating the recording. V1 activation was identified by averaging frames across repetitions, obtaining the mean during the 2s baseline and during the 0.5s post-stimulus period, and then calculating the ratio of the post-stimulus mean to the baseline mean, where significant intrinsic signals were detected. These signals were then superimposed onto the vessel images to locate V1 for subsequent two-photon imaging.

### *In vivo* imaging: data acquisition and preprocessing

#### Locomotion speed

Following a two-week post-surgical recovery period, mice were habituated to head fixation for a minimum of 5 days until they demonstrated free running on the custom-built cylindrical treadmill (made in-house using an 8(W) x 4(H) inch round foam cake dummy and a rod through the center, surface covered with black Gaffers tape) before imaging experiments started. A E2 encoder (US Digital) was used to read out the shaft speed and the obtained digital signal was then converted into an analog voltage using a microcontroller (Arduino Micro) and was sent to NI DAQ for acquiring locomotion speed data at 30kHz using Thorsync 4.0 (Thorlabs). Speed data was smoothed with 1D convolution, binned to 30 Hz, and converted to cm/s unit using custom MATLAB and Python scripts.

#### Pupillometry

An in-house-built pupillometry system was used to monitor eyelid blinks and pupil changes. An infrared (850nm) LED (CM-IR30, CMVision) was used to illuminate the left eye (ipsilateral to the recording site). To evenly diffuse the light, an opaque piece of plastic was placed in front of the LED. Video recording was conducted with 1280 x 1024 pixels, at a frame rate of 19.06 Hz using a Chameleon3 monochrome camera (PointGrey FLIR, CM3-U3-13Y3M-CS) and the PointGrey FlyCap2 2.13.3.61 software. The Chameleon3 camera was equipped with a lens (Thorlabs, MVL16M23) attached via an extension adapter (Thorlabs, CML05). To minimize interference from ambient light and the two-photon laser, a 850/40 nm bandpass filter (Thorlabs, FB850-40) was positioned in front of the camera lens. Video preprocessing was performed offline using Fiji^71^ and Facemap^72^, followed by further processing with custom Python scripts. Frames containing artifacts from the trigger signal were removed at the start of the recording. Both blink and pupil data were z-scored for normalization. The blink data was smoothed with a Hanning window, and velocity calculation was performed to detect blink onset and offset. Blinks were adjusted by adding a buffer period before onset and after offset, with nearby blinks merged if a predefined threshold was met. Frames containing blinks were excluded from pupil data, and the resulting gaps were filled using cubic spline interpolation^73^.

#### Visual stimuli and trial design

Visual stimuli were generated using custom MATLAB (Mathworks) scripts with Psychtoolbox-3^68,69^ and presented at a distance of 16 cm from the right eye on a gamma-corrected LED-backlit LCD monitor (Dell P2317H, 509mm X 286 mm) with a mean luminance of 20 cd/m^2^.

In the spontaneous activity experiments, the screen was set to either a powered off state (dark screen experiment) or displayed a uniform mean luminance (gray screen experiment), the recordings usually last 730s.

In the visual orientation tuning experiments, a 10s gray screen preceded the trials as the baseline. Each 6s trial, randomized in order, began with a 1s gray screen, followed by 2s of full-screen moving gratings at 80% contrast, with a 1 Hz temporal frequency and a 0.04 cpd spatial frequency, and ended with a 3s gray screen. The moving gratings were presented in 12 distinct orientations, each separated by 30 degrees and were repeated 10 times for interneuron imaging or 30 times for excitatory neurons in the Tox experiments.

In the visual contrast tuning experiments, a 10s gray screen preceded the trials as the baseline. Each 3s trial began with a 0.8s gray screen, followed by 2s of full-screen moving gratings at one of six contrasts (80%, 60%, 40%, 20%, 10%, 5%), one of eight orientations (45 degrees apart), with a 1Hz temporal frequency and a 0.04cpd spatial frequency, and ended with a 0.2s gray screen. Each condition was repeated 15 times, with one blank trial randomly placed in (a continued gray screen instead of moving gratings) every 20 trials. All trials were randomized in order.

#### Two-photon calcium imaging

Imaging was performed with a custom-built two-photon microscope (Thorlabs, Bergamo®) equipped with a 8 kHz galvo-resonant scanner, Pockels cells and photomultiplier modules (PMTs). Tunable ultrafast lasers (Spectra-Physics, InSight®X3) were set at 920nm (tunable) for GCaMPs and 1045nm (fixed) for tdTomato. The objective was a 16x water immersion lens with a 0.8 numerical aperture (Nikon). Images were acquired with a 512 × 512 pixels field of view (412 x 412µm), targeting cells in L1 (<100µm under the pia mater) or L2/3 (120–250µm under the pia mater), using ThorImage 4.3 (Thorlabs) at a frame rate of 30 Hz. The laser power was adjusted up to 50 mW at the objective’s front aperture. During the imaging, the ultrasound water gel was used under the objective and black masking tape (Thorlabs, T743-1.0) was used to shield light from the screen.

To synchronize calcium imaging, locomotion speed, and pupillometry recordings, a small block of bright pixels was positioned at the bottom left corner of the screen. This signal was captured by a photodiode (Thorlabs, SM05PD1A) affixed to a duplicate secondary screen, processed by a custom current-to-voltage converter to generate a trigger, ensuring precise synchronization across the different data streams.

Calcium imaging data were preprocessed with Suite2p^74^ for motion correction and region of interest (ROI, or neuron) extraction. Sessions with significant z-drift movement were excluded. In some experiments, tdTomato signals were recorded as control for movement, and no significant signals were observed during locomotion of the animals. For every recorded field of view, detected ROIs were semi-manually adjusted based on identifiable cell bodies. ROIs with lower somatic than neuropil signals were excluded. Raw traces extracted by Suite2p were further processed in Python with custom scripts. Neuropil contamination was corrected using *F* = *F*_*somatic*_ − *F*_*neuropil*_ ∗ *neuropil factor* (*r*) with *r* = 0.7 for GCaMP6f ^75^ and GCaMP7s^76^, and *r* = 0.8 for jGCaMP8m^66,77^. Baseline fluorescence (F0) was estimated by identifying the 30th percentile over a moving window of 150s^41^. dF/F0 traces were computed by subtracting and dividing the raw trace by F0. To get standardized dF/F0, dF/F0 traces were then normalized by subtracting the median and dividing by the standard deviation^78^.

#### Data storage

All preprocessed data were stored in HDF5 format using Python and experimental metadata were stored into a database with MySQL Workbench 8.0.34.

### *In vivo* imaging: data analysis

#### Synchronization of activity

To assess the synchronization of neuronal activity, the standardized dF/F0 for each identified ROIs (neurons) was interpolated to 10 Hz, smoothed with a 5-point moving average, and then decimated to 5 Hz. ROIs with standardized dF/F0 ≥ 3 (at least 3-fold standard deviations from the mean) were included for this analysis. Pairwise zero-time cross-correlation, and Pearson’s correlation coefficients were computed based on the 5 Hz activity data from identified ROIs within the same field of view (FOV). The significance of correlation (p<0.05) was computed by shuffling one of the paired activities 1000 times.

#### Spontaneous activity

To evaluate the spontaneous activity of neurons, the standardized dF/F0 of each ROI (neuron) were processed, along with locomotion speed and z-scored pupil area. ROIs with standardized dF/F0 ≥ 3 (at least 3-fold standard deviations from the median) were included for further analysis. Experimental recordings were excluded if the mouse did not run during the trial. To assess the spontaneous activity of neurons, frames were classified based on the subject’s movement status: stationary (speed ≤ 1 cm/s) and running (speed > 1 cm/s). The average neuronal activity during each state was computed for individual neurons. The 2D density plot was plotted with the ‘scipy.stats.gaussian_kde’ function in Python. For correlation between neuronal activity and locomotion speed, data were first interpolated to a 10 Hz sampling rate, smoothed using a 5-point moving average, and then decimated to 5 Hz. The zero-time cross-correlation between neuronal activity and locomotion speed was computed, and the Pearson’s correlation coefficients were calculated at zero-time. The significance of these correlations was assessed through a shuffle test involving 1000 permutations of the locomotion speed data, with a significance threshold set at p<0.05. Neurons displaying significant correlations were further analyzed to identify the maximum correlation value and the corresponding time lag.

#### Orientation visual responses

To investigate the orientation tuning properties of neurons in V1, their visual responses to drifting gratings were analyzed at the population level. Visual responses were calculated on the average standardized dF/F0 during the period of moving grating presentation, adjusted by subtracting a 0.5s baseline period prior to the onset of the grating for each trial, then the responses were averaged across multiple repeats for each orientation tested for individual neurons. The neuron was categorized as a responsive neuron if in at least two trials it had a response that was 3 times larger than standard deviation of the baseline.

The global orientation selectivity index (gOSI) was calculated with

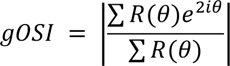

where *θ* was the orientation angle of moving gratings, and *R*(*θ*) denoted the mean response to moving gratings at that orientation, *i* was the imaginary unit.

The direction selectivity index (DSI) was calculated with

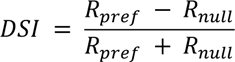

where *R*_*pref*_ was the neuron’s average response to the preferred orientation (the orientation that elicits the strongest response) and *R*_*null*_ was its averaged response to the antipodal orientation (180° opposite to the preferred orientation). To avoid the impact of negative values in the standardized dF/F0 data, we normalize the response following (*R* − *R*_*min*_)/(*R*_*max*_ − *R*_*min*_) before calculation of gOSI and DSI.

The *SNR* was calculated by averaging across *SNR*_*θ*_ for each neuron. For each orientation, it was computed by dividing the square of the mean response by the standard deviation of responses to repetitions for each orientation.

Signal correlations were calculated for each neuronal pair from the same recording. To avoid spontaneous correlations, half of the repetitions were randomly selected from neuron *j* to get the mean response for each orientation. The mean response from neuron *k* was averaged from the other half repetitions for each orientation. The neuron’s mean response to each orientation was calculated as <u>*R*</u>.

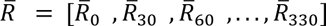

Pairwise signal correlation 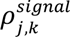 was calculated as the Pearson’s correlation coefficient between *R*_*j*_ and *R*_*k*_:

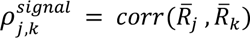

where *j* and *k* indicate neuron *j* and neuron *k*, respectively.

Noise correlations were calculated for each neuronal pair from the same recording^79^. The neuron’s response to each repetition of that orientation *θ* was denoted as:

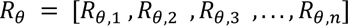

where n is the number of repetitions per orientation. Pairwise noise correlation 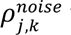 was calculated as the averaged Pearson’s correlation coefficient between *R*_*j*,*θ*_ and *R*_*k*,*θ*_ across all 12 orientations:

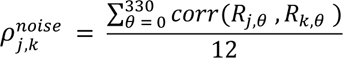

#### Contrast responses of cINs

To access the visual responses of cINs to moving gratings of varying contrasts, visual response for each trial was calculated as the mean standardized dF/F0 activity during the presentation of moving grating stimuli subtract the mean activity from baseline (the 0.5s period before the stimuli). The neuron was considered responsive and included in the analysis if it exhibited responses in at least two trials that were more than three times the standard deviation of the baseline activity. In this experiment, almost all neurons recorded were identified as ‘responsive’. Visual responses were averaged across multiple repetitions for different orientations at the same contrast level. Contrasts of 5% and 10% were classified as low contrasts, while 60% and 80% contrasts were considered high contrasts. Trials were classified into stationary (mean speed ≤ 1 cm/s) or running (mean speed > 1 cm/s) trials based on the averaged speed during visual stimulus presentation for each trial. Contrast preference computed by the log-scale center-of-mass *c*_*COM*_^41^, which was calculated from mean visual responses to various contrast stimuli averaged across all orientations:

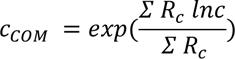

where *c* is the contrast of moving gratings, *R*_*c*_ is the averaged visual responses across all orientations at contrast *c*.

### Retrograde monosynaptic rabies tracing

For tracing inputs from LAMP5+ cINs in V1, *JAM2-Cre* mice at 2-5 month-old were stereotaxically injected with AAV1.DLX.DIO.TVA (titer: 3.5 x 10^12^ vg/mL, diluted to 9 x 10^11^ vg/mL), AAV1.DLX.DIO.GFP.N2cG (titer: 2.9 x 10^12^ vg/mL, diluted to 7 x 10^11^ vg/mL) and rabies virus EnvA-pseudotyped CVS-N2c(DG)-FlpO-mCherry (titer: 3.7 x 10^9^ U/ml, diluted to 1 x 10^8^ U/ml) at the same time in V1 (from Bregma: AP -3, ML ±2.5, DV 0.25-0.5 mm) with a total volume of 60 nl using Nanoject III (Drummond) at 1nl/s. The rabies virus construct was a gift from Thomas Jessell (Addgene #73471^80^) and EnvA-pseudotyped CVS-N2c(ΔG)-FlpO-mCherry was generously shared by K. Ritola at Janelia Farms Research Center as described in ^81^.

Animals were sacrificed and perfused 13 days later and brain tissue was collected. Fixed brain samples were sectioned to 50 µm slices with vibratome (Leica). Sections were analyzed every 150 μm along the rostral-caudal axis with immunohistochemistry to examine the rabies tracing patterns. Images were collected using a whole slide scanning microscope with a 10X objective (Olympus VS120 slide scanners).

NeuroInfo software (MBF Bioscience) was used for image registration and cell detection. All brain sections were manually reordered from rostral to caudal of the brain and the border of each brain section was identified. Initial alignment of sections utilized the software’s Most Accurate alignment option, followed by manual adjustments as necessary. A fixed distance of 150 µm between each section was specified. The Section Registration function of the software was then used to estimate the rostral-caudal location of each section by comparing it to an Allen Mouse Brain Common Coordinate Framework. Non-linear registration was applied to account for any minor distortions introduced during sectioning, mounting, or imperfections in sectioning angle. Cell detection parameters, including cell size and distance from background, were adjusted to optimize detection accuracy. Detection of rabies-infected cells in the red channel was performed using a Neural Network with the preset pyramidal-p64-c1-v15.pbx. Detected cells were manually reviewed to correct any potential detection errors. Additionally, starter cells were manually identified and marked as GFP co-localized rabies-infected cells.

### Perfusion

For all histological experiments, mice were deeply anesthetized with sodium pentobarbital (Euthasol) by intraperitoneal injection and transcardially perfused with 1X PBS followed by 4% paraformaldehyde (PFA) in 1X PBS. Brains were dissected out and post-fixed overnight at 4°C or 2 hours at room temperature. Fixed brain samples were then cryopreserved in 30% sucrose in 1X PBS.

### Immunohistochemistry

40 μm brain sections (if not specifically stated) were obtained through a Leica sliding microtome and preserved in the antifreeze buffer (30% glycerol, 30% ethylene glycol in 1X PBS) and stored at -20°C before experiments. Free-floating brain sections were incubated in blocking solution (10% normal donkey serum, 0.3% Triton X-100 in 1X PBS) at room temperature for 1 hour, followed by incubation in primary antibody diluted in blocking solution at 4°C overnight. The following day, sections were rinsed in 1X PBS for 10 minutes 3 times, followed by secondary antibody incubation in the same blocking solution for 1 hour at room temperature. Sections were then rinsed in 1X PBS for 10 minutes 3 times. Sections were counterstained with DAPI (5 μM in 1x PBS, Sigma #D9542) for 5 minutes and mounted using Fluoromount-G (Invitrogen). Images were collected using a whole slide scanning microscope with a 10X objective (Olympus, VS120 slide scanner) or using a motorized tiling scope (Zeiss, Axio Imager A1) with a 10X objective. Primary antibodies used in this study include Rabbit-anti-DsRed (Clontech #632496, 1:1000), Goat anti-GFP (Sicgen, #AB0020-200), Chicken-anti-GFP (Aves Labs, #1020), Rabbit-anti-SST (Peninsula Laboratories, #T4103, 1:3000). Secondaries (dilute at 1:500) used in this study include Alexa Fluor 488 Donkey anti-Goat (Thermo Fisher Scientific, #A-11055), Alexa Fluor 488 Donkey anti-Chicken (Jackson ImmunoResearch Labs, #703-545-155), Alexa Fluor 594 Donkey anti-Rabbit (Thermo Fisher Scientific, #A-21207), Alexa Fluor 647, Donkey anti-Rabbit (Thermo Fisher Scientific, #A-31573).

### RNAscope^®^ with Immunohistochemistry

20 μm brain sections were obtained using a Leica sliding microtome and preserved in section storage buffer (28% (w/v) sucrose, 30% (v/v) ethylene glycol in 0.1M sodium phosphate buffer, pH 7.4) and stored at -80 °C before the RNAscope^®^ experiments. Samples were processed according to the ACDBio Multiplex Fluorescent v2 Kit protocol (ACDBio #323100) for fixed frozen tissue. Briefly, tissue was pre-treated with a series of dehydration, H2O2, antigen retrieval, and protease III steps before incubation with the probe for 2 hours at 40 °C. Note here protease III incubation was performed at room temperature to better preserve protein. Three amplification steps were carried out prior to developing the signal with Opal™ or TSA^®^ Dyes (Akoya Biosciences). Immunostaining following the RNAscope^®^ experiment was performed in some experiments according to Technical Note 323100-TNS from ACDBio. Primary antibody Rabbit-anti-DsRed (Clontech #632496, 1:1000) and secondary antibody HRP-goat-anti-rabbit (1:500) was used, followed by Opal™ or TSA® Dyes for tdTomato or dTomato protein immunostaining. Sections were counterstained with DAPI (5 μM, Sigma #D9542) and mounted using Fluoromount-G (Invitrogen) or Prolong Gold antifade mounting medium. Images of RNAscope^®^ experiments were acquired with an upright confocal microscope (Zeiss LSM 800) with a 10X or 20X objective (Plan-Apochromat 10x/0.45 420640-9900, 20x/0.8 420650-9901). To analyze the marker RNA expression in genetically labeled neurons, labeled neurons were segmented using threshold method after background subtraction and applying gaussian blur filter in Fiji. Probe signals were detected by thresholding methods and the presence of probe signals within the boundary of segmented neurons was quantified as probe-positive labeled neurons. Probes used in this study include: DLX5-C1(#478151), GAD2-C2(#400951-C2), LHX6-C1(#422791), LSP1-C3(#511811-C3), NPY-C2(#313321-C2), NDNF-C1(#447471), NDNF-C3(#447471-C3), PVALB-C1(#421931), SNCG-C1(#482741), SST-C1(#404631), SV2C-C1(#545001), VIP-C3(#415961-C3).

### *In vitro* electrophysiology: whole-cell patch clamp

Mice were perfused with NMDG-HEPES aCSF containing 93 mM NMDG (Sigma, #M2004), 2.5 mM KCl (Sigma, #P9541), 1.2 mM NaH_2_PO_4_ (Himedia, #GRM3964), 30 mM NaHCO_3_ (Sigma, #S6014), 20 mM HEPES (Sigma, #H3375), 25 mM glucose (Sigma, #G8270), 2 mM thiourea (Sigma, #T8656), 5 mM Na-ascorbate (Sigma, #A4034), 3 mM Na-pyruvate (Sigma, #P2256), 0.5 mM CaCl_2_.2H_2_O (Quality Biological, #351-130-721) and 10 mM MgSO_4_.7H_2_O (Quality Biological, #351-033-721), equilibrated with hydrochloric acid (Sigma, #H1758) to pH 7.3–7.4. Mice were then decapitated, and the brain was quickly removed and immersed in NMDG-HEPES aCSF. 300 μm thick coronal slices were cut using a vibratome (Leica VT 1200S) through V1. Slices were recovered in a holding chamber with HEPES holding aCSF containing 92 mM NaCl (Sigma, #S3014), 2.5 mM KCl (Sigma, #P9541), 1.2 mM NaH_2_PO_4_ (Himedia, #GRM3964), 30 mM NaHCO_3_ (Sigma, #S6014), 20 mM HEPES (Sigma, #H3375), 25 mM glucose (Sigma, #G8270), 2 mM thiourea (Sigma, #T8656), 5 mM Na-ascorbate (Sigma, #A4034), 3 mM Na-pyruvate (Sigma, #P2256), 2 mM CaCl_2_.2H_2_O (Quality Biological, #351-130-721) and 2 mM MgSO_4_.7H_2_O (Quality Biological, #351-033-721), equilibrated with NaOH (Macron, #7708-10) or hydrochloric acid (Sigma, #H1758) to pH 7.3–7.4. During recovery, the NaCl was gradually added as described in ^82^. Slices were recovered at 34 °C for 25 minutes and at room temperature for at least 45 minutes prior to recording. All slice preparation and recording solutions were oxygenated with carbogen gas (95% O2, 5% CO2, pH 7.4). For recordings, slices were transferred to an upright microscope (Zeiss) with IR-DIC optics. Cells were visualized using a 40x water immersion objective. Slices were perfused with HEPES recording aCSF in a recording chamber at 2 mL/min at 30°C. HEPES recording aCSF contains 124 mM NaCl (Sigma, #S3014), 2.5 mM KCl (Sigma, #P9541), 1.2 mM NaH_2_PO_4_ (Himedia, #GRM3964), 24 mM NaHCO_3_ (Sigma, #S6014), 5 mM HEPES (Sigma, #H3375), 12.5 mM glucose (Sigma, #G8270), 2 mM CaCl_2_.2H_2_O (Quality Biological, #351-130-721) and 2 mM MgSO_4_.7H_2_O (Quality Biological, #351-033-721), equilibrated with NaOH (Macron, #7708-10) or hydrochloric acid (Sigma, #H1758) to pH 7.3–7.4. Patch electrodes (4–6 MΩ) were pulled from borosilicate glass (1.5 mm OD, Harvard Apparatus). For all recordings patch pipettes were filled with an internal voltage-clamp solution containing: 130 mM Cs-methanesulfonate (Sigma, #C1426), 5 mM CsCl (Sigma, #C3032), 10 mM HEPES (Sigma, #H3375), 0.2 mM EGTA.CsOH (Sigma, #E3889), 4 mM MgATP (Sigma, #A9187), 0.3 mM Na_2_GTP (Sigma, #G8877), 8 mM Phosphocreatine-Tris_2_ (Sigma, #P1937), 5 mM QX314-Cl (Tocris, #2313), 0.4% biocytin (Sigma, #B4261), equilibrated with 0.5 M CsOH at pH 7.3, or an internal current-clamp solution containing: 130 mM K D-gluconate (Sigma, #G4500), 5 mM KCl (Sigma, #P9541), 10 mM HEPES (Sigma, #H3375), 0.2 mM EGTA.KOH (Sigma, #E3889), 4 mM MgATP (Sigma, #A9187), 0.3 mM Na_2_GTP (Sigma, #G8877), 8 mM Phosphocreatine-Tris_2_ (Sigma, #P1937) and 0.4% biocytin (Sigma, #B4261), equilibrated with 1 M KOH at pH 7.3. Recordings were performed using a Multiclamp 700B amplifier (Molecular Devices) and digitized using a Digidata 1440A and the Clampex 10 program suite (Molecular Devices). Cells were only accepted for analysis if the initial series resistance was less than 40 MΩ and did not change by more than 20% during the recording period. No corrections were made for the liquid junction potential. Intrinsic properties were obtained from JAM2+ cells labeled by AAV virus at a holding potential of -60 to -65 mV with the current-clamp internal. For optogenetic-assisted circuit mapping, voltage-clamp signals were filtered at 3 kHz and recorded with a sampling rate of 20 kHz. IPSCs were recorded at a holding potential of 0 mV. Whole-cell patch-clamp recordings were obtained from JAM2+ cells labeled by AAV virus and unlabeled putative PYN or L4-5 SST+ cINs labeled by AAVPHP.s9e10.dTom (unpublished SST enhancer) in the same column with the voltage-clamp internal. To activate afferents expressing hChR2, blue light was transmitted from a collimated LED (Mightex) attached to the epifluorescence port of the upright microscope. 5 ms pulses of a fixed light intensity were directed to the slice in the recording chamber via a mirror coupled to the 40x objective. Flashes were delivered every 15s for a total of 10 trials. The LED output was driven by a transistor-transistor logic output from the Clampex software. Recordings were performed in the presence of 1 µM TTX and 1 mM 4-AP (Tocris). In some experiments, the IPSCs were confirmed with 10 µM SR95531 (Tocris, #1262). Data analysis was performed offline using the Clampfit module of pClamp (Molecular Devices). Individual waveforms from all trials per cell were averaged, and the averaged peak amplitude was analyzed.

### Statistical analysis

Statistical details of experiments can be found in the figure legends and supplementary tables. All statistical analyses were performed with hierarchical bootstrap^83^, linear mixed effect models (considering animal as random effect)^84^ or one-way ANOVA. Mann Whitney test (such as ratios) or Wilcoxon signed rank test (such as two groups from the same animals) were used for animal level data. GraphPad Prism and Python were used to conduct statistical tests. *p < 0.05, ** < 0.01, *** < 0.001, **** < 0.0001.

## ACKNOWLEDGEMENTS

This work was supported by grants from the National Institutes of Health R01 NS081297, R37 MH071679, P01 NS074972, UG3 MH120096, and the Simons Foundation Autism Research Initiative to G.F., NINDS P30 NS072030 to Harvard Medical School Neurobiology Imaging Facility, Quan Predoctoral Fellowship (FY22), Edward R. and Anne G. Lefler Center Predoctoral Fellowship (FY23, FY24) and Ryan Fellowship (FY23) to S.H.. We thank Dr. Emilia Favuzzi, Dr. Mark Andermann and Dionnet Bhatti for comments on the manuscript. We thank Dr. John Ngai, Dr. David A. Stafford, Dr. Susan M. Dymecki, Dr. David Ginty, Dr. Tanya L. Daigle, Dr. Bosiljka Tasic and Dr. Hongkui Zeng for generously sharing the genetic mouse strains. We thank Dr. Kimberly Ritola for sharing the rabies virus. We thank Dr. Gabrielle Pouchelon for sharing the VTK plasmid toolkit. We thank Dr. Jordane Dimidschstein and Sofi Vergara for sharing AAVPHP.s9e10.dTom enhancer AAV for SST targeting. We extend our sincere gratitude to Dr. Xindong Song for his invaluable guidance for the intrinsic imaging experiments. We thank John LeBlanc (HMS Machine Shop), Pavel Gorelik, and Dr. Ofer Mazor (HMS Research Instrumentation Core) for their assistance in engineering the *in vivo* imaging platform. We are grateful to Dr. Sergey Matveev and Dr. Utsab Khadka (Thorlabs) for their expertise and assistance with the two-photon microscopy setup. We thank Dr. Min Dai for discussion on transcriptomic data analysis. We thank all colleagues in the Fishell laboratory, Dr. Christopher D. Harvey, Dr. Anne Takesian, Dr. Daniel Polley, Dr. Bernardo Rudy, and Dr. Robert Machold for helpful discussion on this project over the years.

## AUTHOR CONTRIBUTION

Conceptualization and methodology, S.H.; investigation, S.H., D.R., S.J.W., L.Z.; analysis, S.H., D.R.; resources - plasmids and AAV, Q.X., S.H.; resources - mouselines, D.A.S., T.L.D., B.T., H.Z.; writing - original draft, S.H.; writing - review and editing, G.F., L.A.I., S.J.W; supervision, G.F., L.A.I.; funding and resources, G.F..

