## Supplemental figures and tables for "Neurogliaform Cells Exhibit Laminar-specific Responses in the Visual Cortex and Modulate Behavioral State-dependent Cortical Activity"

### Extended Data 1 - related to Figure 1

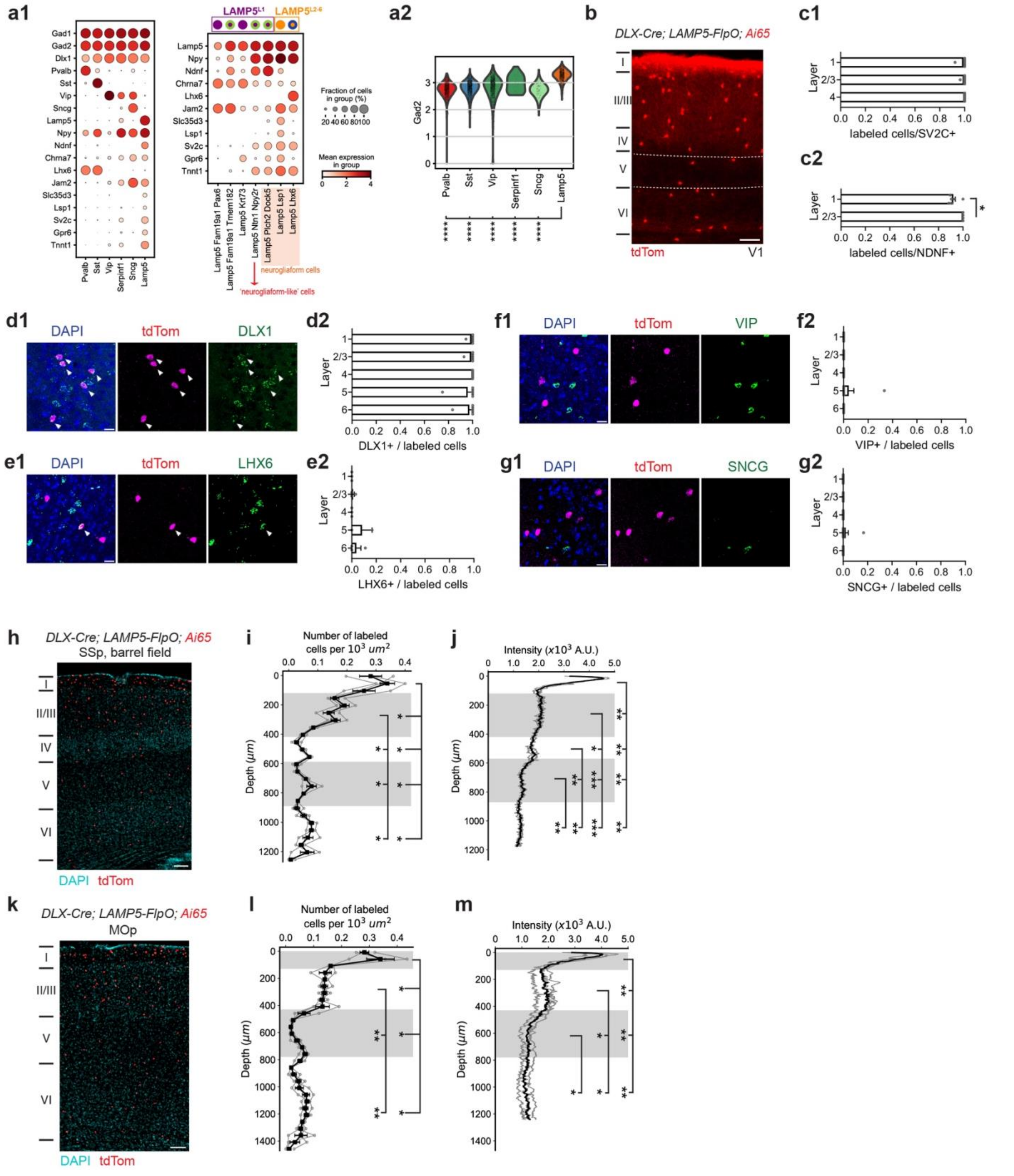

**Extended Data 1 (Related to Figure 1) - Further characterization of LAMP5+ cINs with *DLX-Cre; LAMP5-FlpO; Ai65(RCFL-tdT)*.**

(a1) Enriched gene expression log(CPM+1) (log counts per million) from scRNAseq data [10] for both (left) cardinal cINs and (right) molecular subclusters within LAMP5+ cINs. log(CPM+1)<1 was set to 0. Nearly all $LAMP5^{L2\text{-}6}$ belong to the Lamp5/Lsp1 molecular cluster, with a minority subset falling into the Lamp5/Plch2/Dock5 molecular cluster. Both of these clusters closely align with the neurogliaform cells described in the literature[6,12,13,18]. The other molecular cluster in $LAMP5^{L2\text{-}6}$ expresses LHX6 (Lamp5/Lhx6, labeled by the blue boundary) and represents a minority neurogliaform cell population (<0.5% of cINs) in the deep layers of the cortex. This subset exhibits similarities to hippocampal Ivy cells and has been found to be more prominent in the primate cortex[12,33,85,86]. Within L1 there are no excitatory neuron cell bodies; residing there are only cINs commonly referred to as L1 interneurons. Approximately 90% of these L1 interneurons fall under the classification of LAMP5+ cINs[5,10]. Among LAMP5+ cINs in L1, about 80% express NDNF (labeled by the green boundary), with around half of this subset (the Lamp5/Plch2/Dock5 molecular cluster) demonstrating the characteristic late-spiking electrophysiological profile associated with classical neurogliaform cells[12,16]. The extent of resemblance between the remaining NDNF+ cINs (mostly in the Lamp5/Ntn1/Npy2r molecular cluster) that lack late-spiking attributes, and conventionally defined neurogliaform cells remains a subject of ongoing discussion. Recent studies, however, suggest more continuity than distinction within the NDNF+ population, with some extremes resembling the previously classified 'canopy' cells[12,16,53]. In addition, the remaining 20% of $LAMP5^{L1}$ express CHRNA7, and are distinct from neurogliaform cells[10,16].

(a2) Violin plot with log(CPM+1) expression of GAD2 in the cardinal cIN types.

(b) Example image of *DLX-Cre; LAMP5-FlpO; Ai65(RCFL-tdT)* genetic labeling in V1 with saturated intensity for neurites visualization. The lowest neurites signal intensity was found in L5 (indicated between dashed lines). Scale bar = 100 μm.

(c) Bar plot showing ratio of *DLX-Cre; LAMP5-FlpO; Ai65(RCFL-tdT)* labeled cells out of (c1) all SV2C+ cells in L1-4 or (c2) all NDNF+ cells in L1-3 of V1. SV2C is specific to LAMP5+ cINs in L1-4, while it is also expressed in a subpopulation of L5-6 excitatory neurons.

(d-g) RNAscope images and bar plots similar to Figure 1e-h. (d1-d2) DLX1, indicative of all cINs. (e1-e2) LHX6; (f1-f2) VIP and (g1-g2) SNCG (a subset of which also expresses VIP).

(h) Example image of *DLX-Cre; LAMP5-FlpO; Ai65(RCFL-tdT)* genetic labeling in SSp barrel field. Scale bar = 100 μm.

(i) Quantification for the number of labeled cells per $10^3$ $\mu m^2$ and (j) the intensity of labeled neurites in $10^3$ arbitrary units (A.U.) in SSp barrel field. Gray indicates L2/3 and L5. Similar to Figure 1c-d.

(k) Example image of *DLX-Cre; LAMP5-FlpO; Ai65(RCFL-tdT)* genetic labeling in MOp. Scale bar = 100 μm.

(l) Quantification for the number of labeled cells per $10^3$ $\mu m^2$ and (m) the intensity of labeled neurites in $10^3$ arbitrary units (A.U.) in MOp. Gray indicates L1 and L5. Similar to Figure 1c-d.

Data from N = 3 animals. Error bar represents SEM. Wilcoxon signed-rank test (a2, g2), repeated measures ANOVA (c1-g1), repeated measures ANOVA followed by uncorrected Fisher's LSD (i-j, l-m) were used for testing statistical significance. See supplementary data 1 - Table 1.1 for statistics.

Abbreviations: SSp: primary somatosensory area, MOp: primary motor area.

Extended Data 2 - related to Figure 1

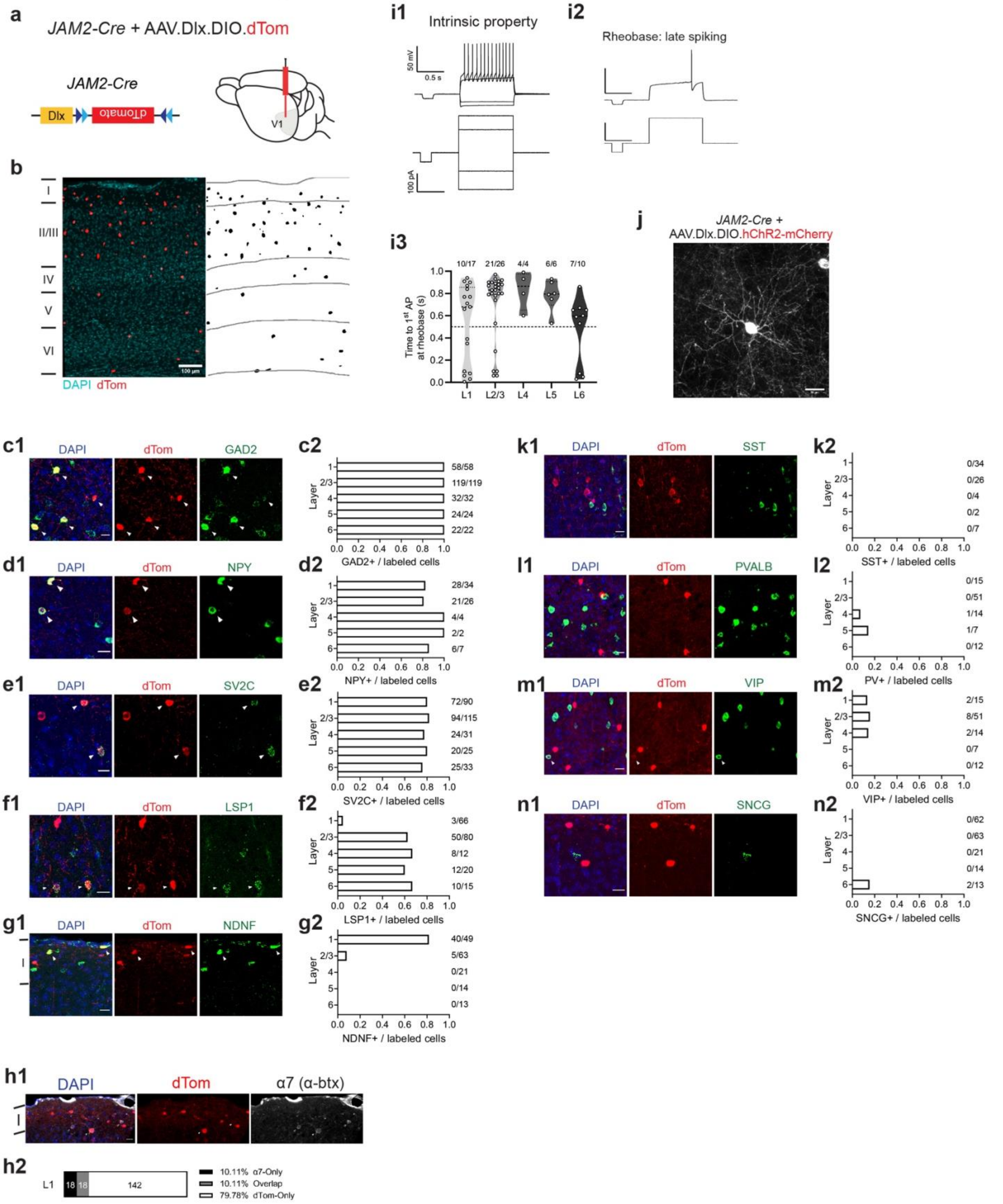

**Extended Data 2 (Related to Figure 1) - An alternative strategy for targeting LAMP5+ cINs using *JAM2-Cre* with AAV.Dlx.DIO intracranial injection.**

(a) Illustration of genetics: *JAM2-Cre* is a knock-in mouse model where Cre is expressed following JAM2 expression. Specifically, the expression of JAM2 becomes restricted to LAMP5+ cINs postnatally (see Method - Mouse). AAV9.Dlx.DIO.dTom was injected into the V1 to restrict expression to cINs and enable visualization of LAMP5+ cINs.
(b) An example of genetic labeling in V1 (left) and segmented cell bodies (right). Scale bar = 100 µm.
(c-g) Representative RNAscope assays images taken by confocal microscopy from the V1 with corresponding bar plots showing the ratios of marker gene-positive cells to dTom labeled cells in each cortical layer. Each panel displays an overlay figure (left) with nuclei dye DAPI (in blue), labeled cells from *JAM2-Cre* with AAV9.Dlx.DIO.dTom (middle, in red), and the marker gene expression detected by RNAscope assay (right, in green). Arrow indicates dTom and marker gene co-localized cells. Number of (marker+ and dTom+)/dTom+ cells were labeled for each layer. (c1-c2) GAD2, indicative of all cINs; (d1-d2) NPY; (e1-e2) SV2C; (f1-f2) LSP1; (g1-g2) NDNF. Scale bar = 20 µm. Data was collected from N = 3 animals.
(h1) Representative images taken by confocal microscopy from L1 of V1. The panel displays an overlay figure (left) with nuclei dye DAPI (in blue), labeled cells from *JAM2-Cre* with AAV9.Dlx.DIO.dTom (middle, in red), and α7 labeled by fluorophore-conjugated α-btx[16] (right, in white). Scale bar = 20 µm. (h2) Quantification showing the ratios of α-btx+ cells among all dTom+ cells in L1 of V1. Data was collected from N = 3 animals.
(i1) An example of intrinsic property from one dTom+ cell. (i2) An example of rheobase from one dTom+ cell showing the classical late-spiking feature of neurogliaform cells. Scale bar x = 0.5 s, y = 50 mV or 100 pA. (i3) Quantification of the time to 1$^{st}$ action potential (AP) at rheobase, and ratio of cells showing the late spiking feature in each layer among all examined dTom+ cells in V1 (threshold was set to 0.5s). Data was collected from N = 12 animals.
(j) An example image taken by confocal microscopy showing a labeled dTom+ cell with short, dense, spherical dendrites and highly ramified thin axons around the cell body. Scale bar = 20 µm.
(k-n) Similar to (c-g), but with negative markers: (h1-h2) SST, marker for SST+ cINs; (i1-i2) PVALB, marker for PV+ cINs; (j1-j2) VIP; (k1-k2) SNCG.

### Extended Data 3 - related to Figure 2

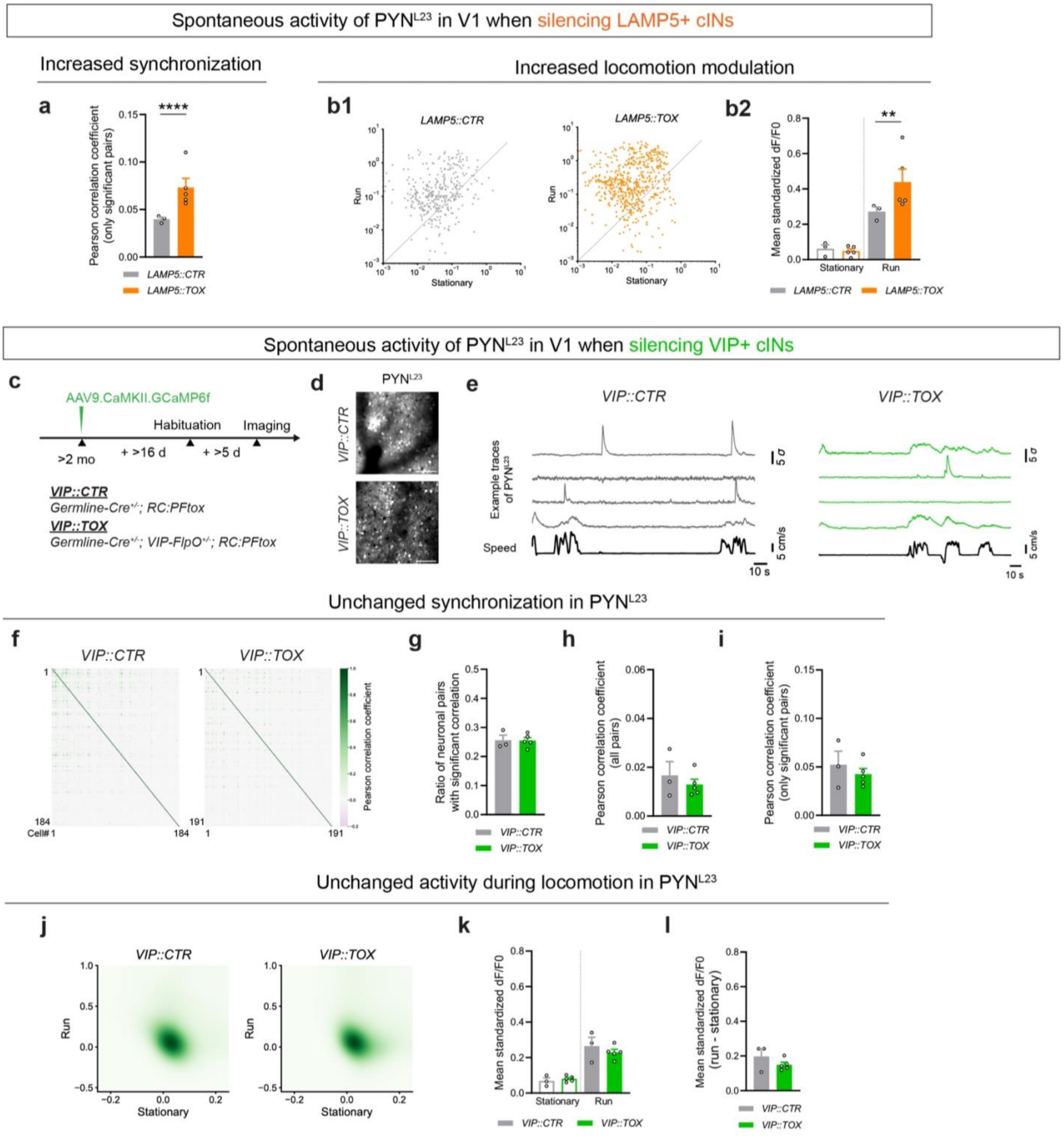

**Extended Data 3 (Related to Figure 2) - LAMP5+ but not VIP+ cIN silencing affects the spontaneous activity of $PYN^{L23}$ in V1 and their synchrony.**

(a) Bar plot showing Pearson's correlation coefficient of all significantly correlated ($p < 0.05$ in shuffle test) neuron pairs in *LAMP5::CTR* (gray) and *LAMP5::TOX* (orange).

(b1) Scatter plot with averaged standardized dF/F0 of spontaneous activity during either stationary or running period for each neuron from *LAMP5::CTR* (gray) or *LAMP5::TOX* (orange). (b2) Bar plot showing mean standardized dF/F0 activity during either stationary or running period for both LAMP5::CTR (gray) and LAMP5::TOX (orange).

(c) Similar to Figure 2a, but the experimental animals were generated by crossing the *Germline-Cre; VIP-FlpO* mouse model with *RC:PFtox*. Mutants (*VIP::TOX*) carried the genotype *Germline-Cre*$^{(+/-)}$*; VIP-FlpO*$^{(+/-)}$*; RC:PFtox*$^{(Flox/+,\ Frt/+)}$. Controls (*VIP::CTR*) were *Germline-Cre*$^{(+/-)}$*; RC:PFtox*$^{(Flox/+,\ Frt/+)}$ genotype. AAV9.CaMKII.GCaMP6f.WPRE.SV40 was injected into V1 to express GCaMP6f preferentially in $PYN^{L23}$.

(d) Example images from the maximum intensity projection of the two-photon imaging experiments, showing GCaMP6f expression within L2/3 of V1 in *VIP::CTR* (upper) and *VIP::TOX* (bottom). Scale bar = 100 μm.

(e) Example traces from spontaneous activity recordings of $PYN^{L23}$ in both *VIP::CTR* (left) and *VIP::TOX* (right). The top four traces in each set (gray for *VIP::CTR* and green for *VIP::TOX*) represent the smoothed standardized dF/F0 activity of four randomly chosen neurons during a randomly selected time interval for representation. The bottom trace (in black) in each set showed the corresponding locomotion speed of the animal (measured in cm/s).

(f) Example heatmap showing Pearson's correlation coefficient of neuron pairs in one imaging session from *VIP::CTR* (left) or *VIP::TOX* (right).

(g) Bar plot showing ratio of significantly ($p < 0.05$) correlated pairs out of all neuron pairs with shuffling (refer to Methods) in *VIP::CTR* (gray) and *VIP::TOX* (green).

(h) Bar plot showing Pearson's correlation coefficient of all neuron pairs in *VIP::CTR* (gray) and *VIP::TOX* (green).

(i) Bar plot showing Pearson's correlation coefficient of all significantly correlated ($p < 0.05$ in shuffle test) neuron pairs in *VIP::CTR* (gray) and *VIP::TOX* (green).

(j) Averaged standardized dF/F0 spontaneous activity for each neuron during either stationary (speed ≤ 1 cm/s) or running (speed > 1 cm/s) period for *VIP::CTR* (left) and *VIP::TOX* (right) groups. The data is fit with a Gaussian kernel for visualization.

(k) Bar plot showing mean standardized dF/F0 activity during either stationary or running period for *VIP::CTR* (gray) and *VIP::TOX* (green).

(l) Bar plot showing the differences in mean standardized dF/F0 activity between running and stationary period, obtained by subtracting the latter from the former, for *VIP::CTR* (gray) and *VIP::TOX* (green).

Error bar represents SEM. Each dot represents the (averaged) result from an individual animal. Mann-Whitney test (g) and hierarchical bootstrap (a,h-i,k-l) were used for testing statistical significance. See supplementary data 1 - Table 2.1 for statistics.

### Extended Data 4 - related to Figure 3

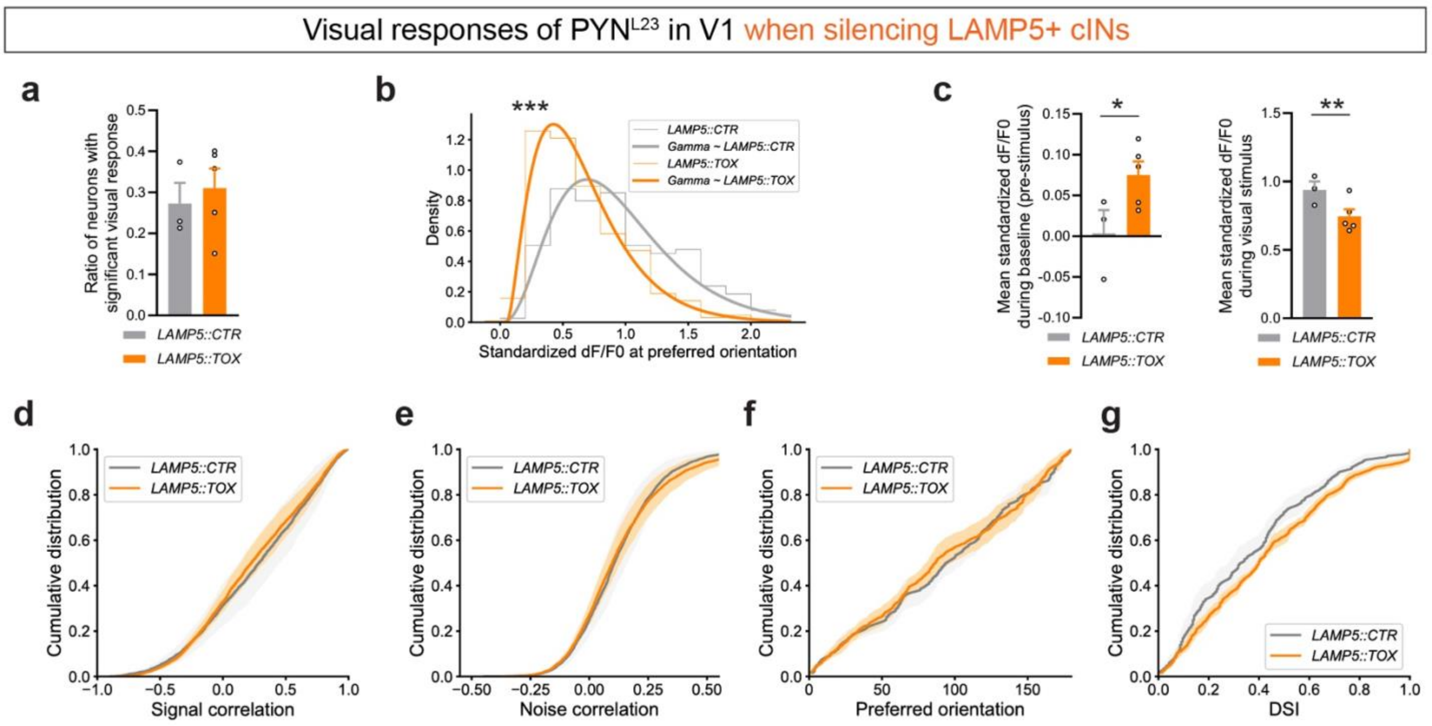


**Extended Data 4 (Related to Figure 3) - LAMP5+ cIN silencing results in impaired visual response properties of PYN$^{L23}$ in V1.**

(a) Bar plot showing the proportion of visually responsive neurons in this experiment in *LAMP5::CTR* (gray) and *LAMP5::TOX* (orange). Each dot represents an individual animal.

(b) Histogram showing probability density distribution and gamma fit of visual responses at preferred orientation for all visually responsive PYN$^{L23}$ in *LAMP5::CTR* (gray) and *LAMP5::TOX* (orange).

(c) Bar plot showing mean standardized dF/F0 of PYN$^{L23}$ (left) during baseline (pre-stimulus period) and (right) during the presence of moving gratings in *LAMP5::CTR* (gray) and *LAMP5::TOX* (orange).

(d) Cumulative ratio of noise correlation of visual responses for *LAMP5::CTR* (gray) and *LAMP5::TOX* (orange).

(e) Cumulative ratio of preferred orientation for *LAMP5::CTR* (gray) and *LAMP5::TOX* (orange).

(f) Cumulative ratio of direction selective index (DSI) for *LAMP5::CTR* (gray) and *LAMP5::TOX* (orange).

Error bar represents SEM. Each dot in the bar plots represents data from an individual mouse. Mann-Whitney test (a) and hierarchical bootstrap (b-f) were used for testing statistical significance. See supplementary data 1 - Table 3.1 for statistics.

### Extended Data 5 - related to Figure 4

#### Spontaneous activity of LAMP5+ cINs in V1

##### Activity Synchronization

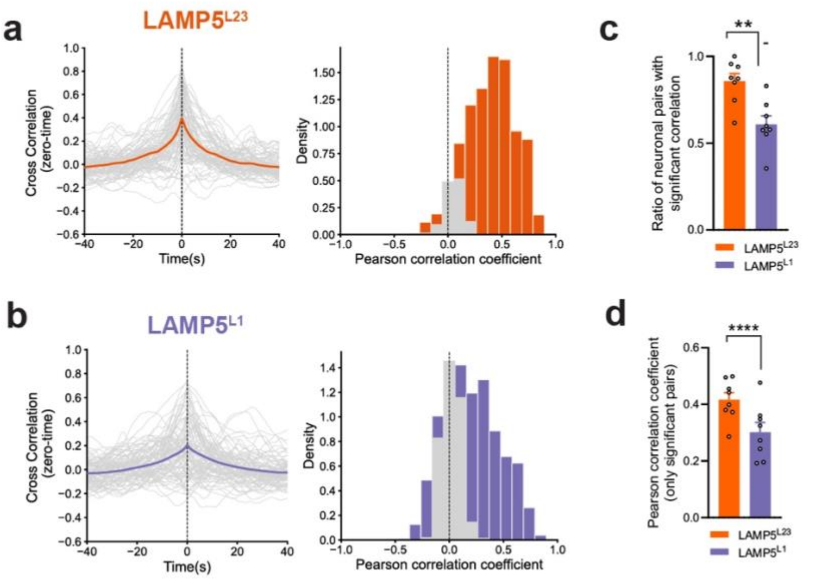


#### Spontaneous activity of LAMP5+ cINs in V1 with dark screen is similar to that with gray screen

##### Neuronal activity and locomotion state

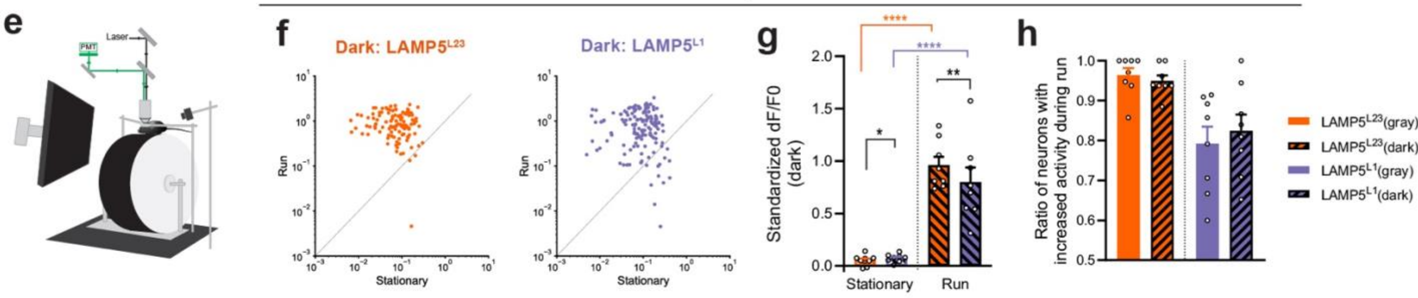


##### Neuronal activity and locomotion speed

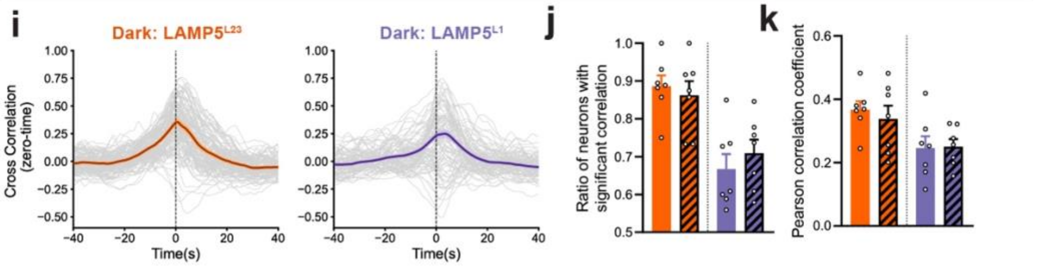

**Extended Data 5 (Related to Figure 4) - Further analysis on LAMP5+ cIN spontaneous activity and behavioral states.**

(a-b) Pairwise zero-time cross-correlation analysis of neuronal activity among $LAMP5^{L23}$ (orange) or $LAMP5^{L1}$ (purple). Histogram of Pearson's correlation coefficient of $LAMP5^{L23}$ (orange) or $LAMP5^{L1}$ (purple) activity pairs. Gray bars represent pairs with no significant correlation, determined by comparing against shuffled time series for each pair for 1000 times (refer to Methods).
(c) Bar plot showing the ratio of significantly correlated pairs in $LAMP5^{L23}$ (orange) or $LAMP5^{L1}$ (purple).
(d) Bar plot showing Pearson's correlation coefficient within significantly correlated $LAMP5^{L23}$ (orange) or $LAMP5^{L1}$ (purple) pairs.
(e) Illustration of the experimental setup during the spontaneous activity recordings. The animal was presented with a dark screen (zero luminance) while the GCaMP signal was imaged with two-photon microscopy.
(f) Scatter plot showing the mean standardized dF/F0 during stationary and running periods with dark screen for $LAMP5^{L23}$ (orange) or $LAMP5^{L1}$ (purple). Each dot represents a neuron.
(g) Bar plot showing the mean standardized dF/F0 during stationary and run periods for $LAMP5^{L23}$ (orange) and $LAMP5^{L1}$ (purple) with dark screen.
(h) Bar plot showing ratio of neurons with increased activity during running periods for $LAMP5^{L23}$ (orange) and $LAMP5^{L1}$ (purple) with gray (no black line filled) or dark (black line filled) screen.
(i) Zero-time cross-correlation analysis between neuronal activity of $LAMP5^{L23}$ (orange) or $LAMP5^{L1}$ (purple), and locomotion speed during the dark screen experiment.
(j) Bar plot showing the ratio of significant correlation between neuronal activity of $LAMP5^{L23}$ (orange) or $LAMP5^{L1}$ (purple), and locomotion speed in the gray (no black line filled) or dark (black line filled) environment.
(k) Bar plot showing Pearson's correlation coefficient between neuronal activity of $LAMP5^{L23}$ (orange) or $LAMP5^{L1}$ (purple), and locomotion speed in the gray (no black line filled) or dark (black line filled) environment.
Each dot in bar plots represents data from an individual animal. Mann-Whitney test (c,h,j,k), mixed linear model regression (g) and hierarchical bootstrap (d) were used for testing statistical significance. See supplementary data 1 - Table 4.1 for statistics.

Extended Data 6 - related to Figure 4

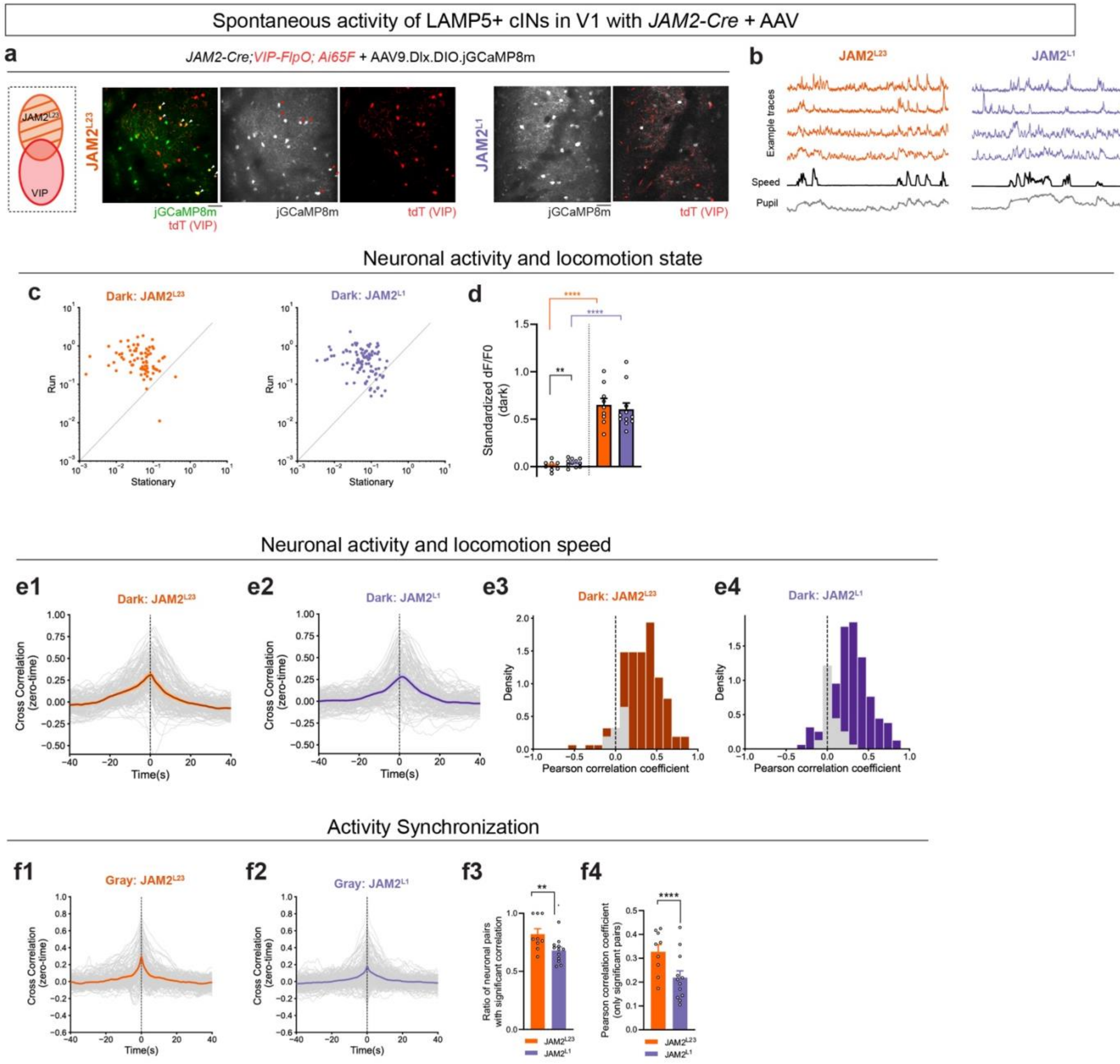

**Extended Data 6 (Related to Figure 4) - Comparable data from *JAM2-Cre* with AAV injection method on spontaneous activity of LAMP5+ cINs.**

(a) Representative mean projection image from two-photon microscopy in L2/3 (left) and L1 (right) of V1 of *JAM2-Cre; VIP-FlpO; Ai65F(RCF-tdT)* with AAV9.Dlx.DIO.jGCaMP8m. *JAM2-Cre* was used as the Cre line for targeting LAMP5+ cINs in adults (Extended Data 2). VIP+ cINs that may get off-targeted by this genetic strategy were labeled by tdTom from *Ai65F* and excluded from analysis.
(b) Example traces from spontaneous activity recordings in $JAM2^{L23}$ (left, in orange) or $JAM2^{L1}$ (right, in purple). The top four traces represent the standardized dF/F0 activity of four randomly chosen neurons during a randomly selected time interval. The black trace indicates the animal's locomotion speed (in cm/s), and the gray trace shows the z-scored pupil size, both measured concurrently.
(c) Scatter plot showing the mean standardized dF/F0 during stationary and running periods with dark screen for $JAM2^{L23}$ (left, in orange) or $JAM2^{L1}$ (right, in purple). Each dot represents a neuron.
(d) Bar plot showing the mean standardized dF/F0 during stationary (white bar) and run (color filled bar) periods with dark screen for $JAM2^{L23}$ (orange) or $JAM2^{L1}$ (purple).
(e) Zero-time cross-correlation analysis between neuronal activity (e1) $JAM2^{L23}$ (orange) or (e2) $JAM2^{L1}$ (purple), and locomotion speed with dark screen. Histogram of Pearson's correlation coefficient for neuronal activity (e3) $JAM2^{L23}$ (orange) or (e4) $JAM2^{L1}$ (purple), and locomotion speed. Gray bars represent pairs with no significant correlation, determined by comparing against shuffling (refer to Methods).
(f) Pairwise zero-time cross-correlation analysis of neuronal activity among (f1) $JAM2^{L23}$ (orange) or (f2) $JAM2^{L1}$ (purple) with gray screen. (f3) Bar plot showing the ratio of significantly correlated pairs in $JAM2^{L23}$ (orange) or $JAM2^{L1}$ (purple). (f4) Bar plot showing Pearson's correlation coefficient within significantly correlated $JAM2^{L23}$ (orange) or $JAM2^{L1}$ (purple) pairs.
Each dot in bar plots represents data from an individual animal. Mann-Whitney test (f3), mixed linear model regression (d) and hierarchical bootstrap (f4) were used for testing statistical significance. See supplementary data 1 - Table 4.2 for statistics.

Extended Data 7 - related to Figure 4

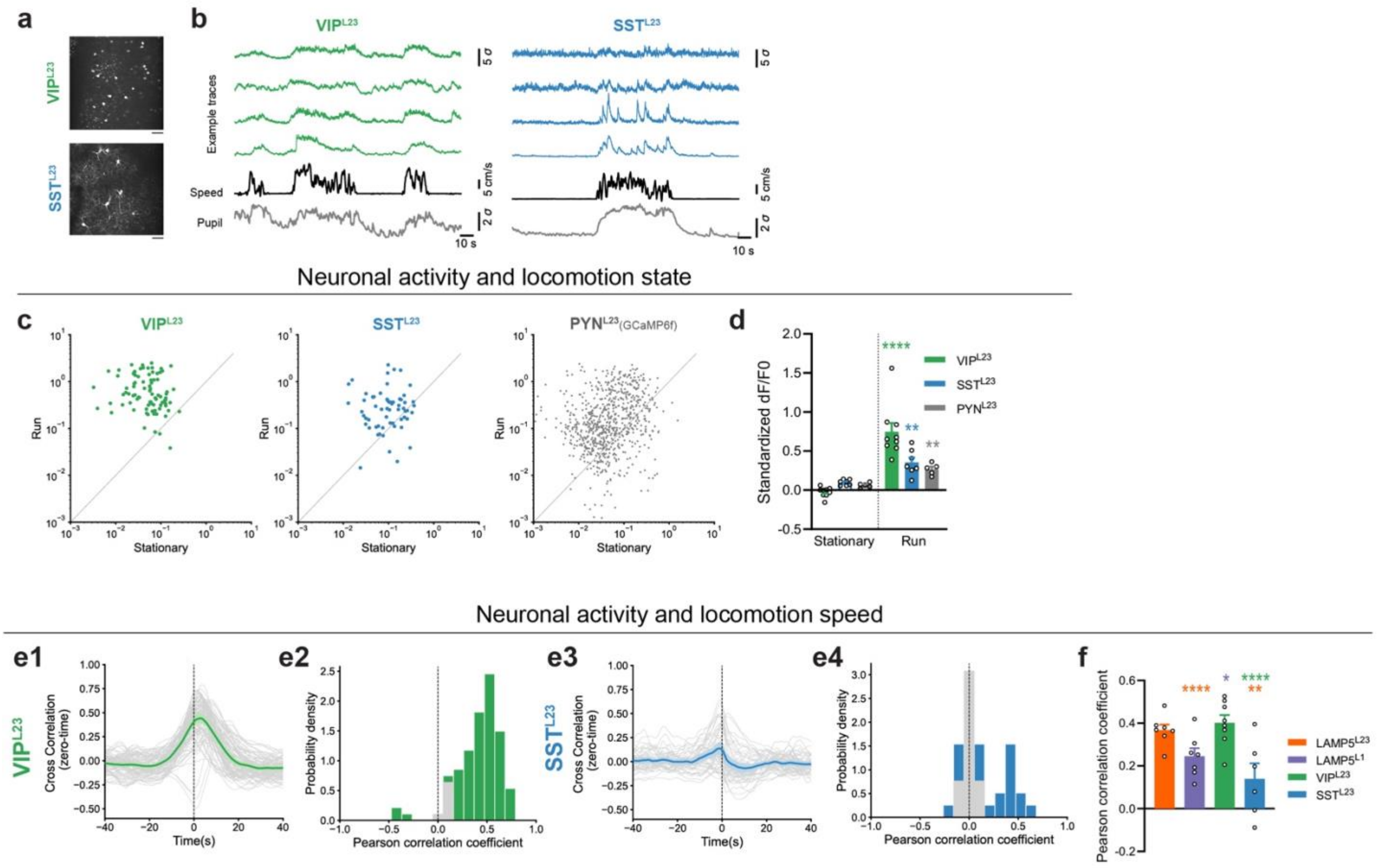


**Extended Data 7 (Related to Figure 4) - Analysis on SST+ and VIP+ cIN spontaneous activity and locomotion speed.**

(a) Representative mean projection image from two-photon microscopy in L2/3 of V1 of *Germline-Cre; VIP-FlpO; Ai195* (upper) or *Germline-Cre; SST-FlpO; Ai195* (bottom). *Ai195* was originally an intersectional reporter, *Germline-Cre* converted *Ai195* to a FlpO-reporter (see Method - Mouse).
(b) Example traces from spontaneous activity recordings in VIP$^{L23}$ (green) or SST$^{L23}$ (blue). The top four traces represent the standardized dF/F0 activity of four randomly chosen neurons during a randomly selected time interval. The black trace indicates the animal's locomotion speed (in cm/s), and the gray trace shows the z-scored pupil size, both measured concurrently.
(c) Scatter plot showing the mean standardized dF/F0 during stationary and running periods for VIP$^{L23}$ (green), SST$^{L23}$ (blue) and PYN$^{L23}$ (gray, data from TOX-controls with AAV.GCaMP6f). Each dot represents a neuron.
(d) Bar plot showing the mean standardized dF/F0 during stationary and run periods for VIP$^{L23}$ (green), SST$^{L23}$ (blue) and PYN$^{L23}$ (gray).
(e) Zero-time cross-correlation analysis between neuronal activity of (e1) VIP$^{L23}$ (green) or (e3) SST$^{L23}$ (blue), and locomotion speed. Histogram of Pearson's correlation coefficient for neuronal activity (e2) VIP$^{L23}$ (green) or (e4) SST$^{L23}$ (blue), and locomotion speed. Gray bars represent pairs with no significant correlation, determined by comparing against shuffling (refer to Methods).
(f) Bar plot showing Pearson's correlation coefficient between neuronal activity and locomotion speed for LAMP5$^{L23}$ (orange), LAMP5$^{L1}$ (purple), VIP$^{L23}$ (green) or SST$^{L23}$ (blue).
Each dot in bar plots represents data from an individual animal. Mann-Whitney test (d) and mixed linear model regression (f) were used for testing statistical significance. See supplementary data 1 - Table 4.3 for statistics.

Extended Data 8 - related to Figure 5

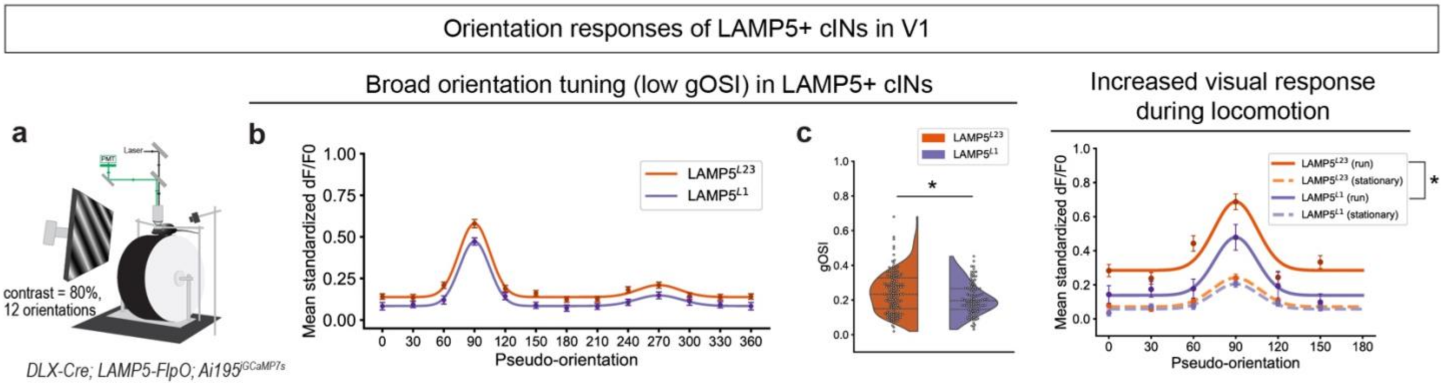


**Extended Data 8 (Related to Figure 5) - LAMP5+ cINs show low orientation selectivity.**

(a) Illustration of the experimental setup during the visual response recordings for orientation tuning. The *DLX-Cre; LAMP5-FlpO; Ai195* animal was presented with moving gratings on the screen, while the GCaMP signal was imaged with two-photon microscopy. Each trial was 6s, consisting of 2s full-field moving gratings with a random orientation and fixed contrast (80%), spatial frequency (0.04 cpd) and temporal frequency (1 Hz), with 4s inter-trial-intervals of gray screen (mean luminance). 12 orientations were examined: 0°, 30°, 60°, 90°, 120°, 150°, 180°, 210°, 240°, 270°, 300°, 330°.
(b) Mean standardized dF/F0 of Pseudo-orientation (set preferred orientation to Pseudo-90° for each neuron) tuning curve for $LAMP5^{L23}$ (orange) and $LAMP5^{L1}$ (purple). Data represents mean and SEM from all neurons.
(c) Violin plot of gOSI values of $LAMP5^{L23}$ (orange) and $LAMP5^{L1}$ (purple). Dot indicates individual neurons.
(d) Mean standardized dF/F0 of Pseudo-orientation (set preferred orientation to Pseudo-90° for each neuron) tuning curve for $LAMP5^{L23}$ (orange) and $LAMP5^{L1}$ (purple) during running and stationary trials. Data represents mean and SEM from all neurons. Hierarchical bootstrap (c) and mixed linear model regression (d) were used for testing statistical significance. See supplementary data 1 - Table 5.1 for statistics.

Extended Data 9- related to Figure 5

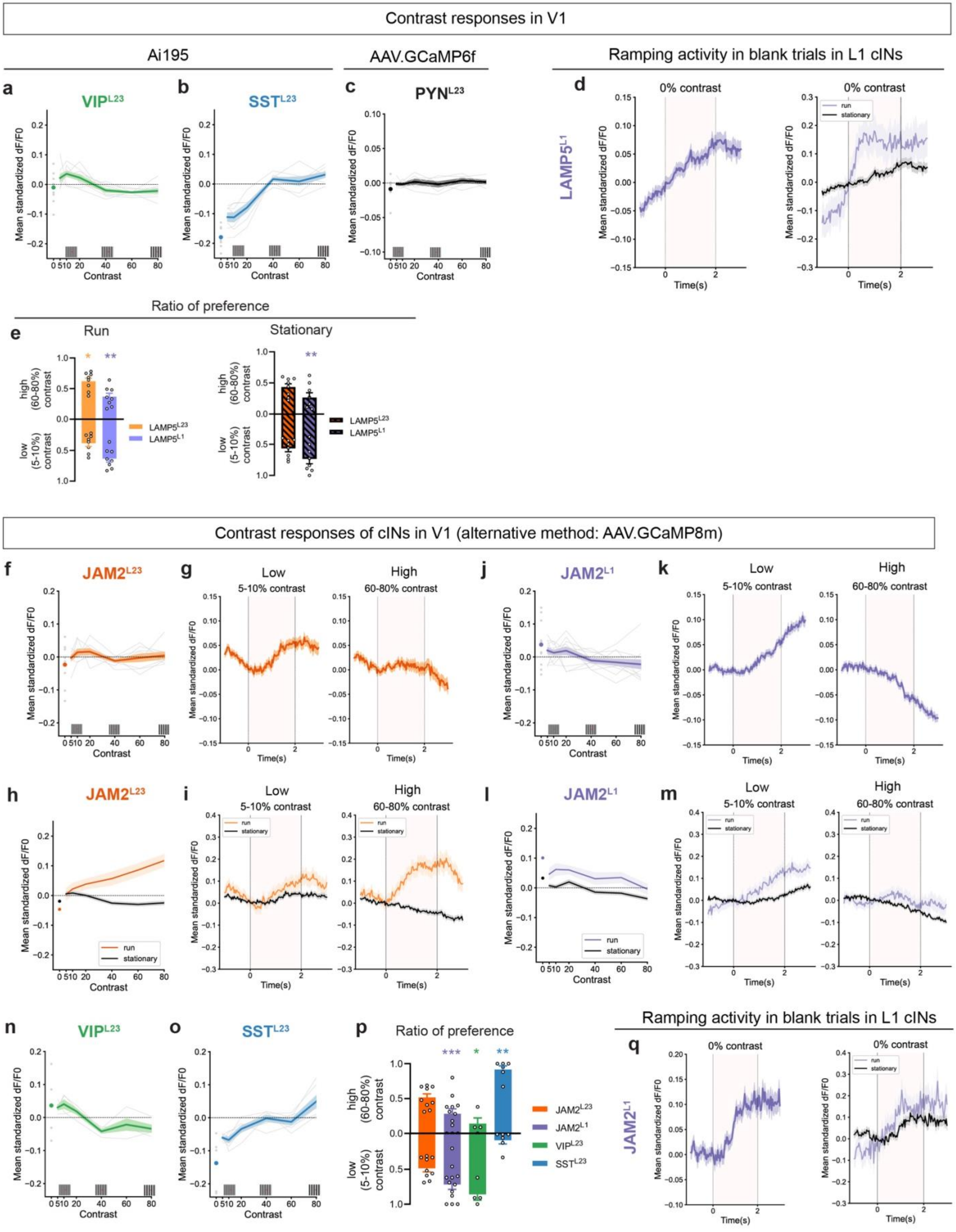

**Extended Data 9 (Related to Figure 5) - Visual responses and locomotion modulation of LAMP5+ cINs.**

(a-c) Visual responses at various contrast levels in (a) $VIP^{L23}$ (green) population labeled by *Germline-Cre; VIP-FlpO; Ai195,* (b) $SST^{L23}$ (blue) population labeled by *Germline-Cre; SST-FlpO; Ai195, and* (c) $PYN^{L23}$ (black) population (data from TOX-control animals with AAV.GCaMP6f).
(d) (left) Averaged response trace in 0% contrast ('blank trials') for $LAMP5^{L1}$. (right) Averaged response trace in 0% contrast ('blank trials') for $LAMP5^{L1}$ during running (purple) or stationary (black) trials.
(e) Ratio of high (60-80%) or low (5-10%) contrast preferring neurons in $LAMP5^{L23}$ (orange), $LAMP5^{L1}$ (purple) during (left) running and (right) stationary trials. Each dot represents data from an individual animal.
(f) Visual responses at various contrast levels in $JAM2^{L23}$ population labeled by *JAM-Cre* with AAV.Dlx.DIO.jGCaMP8m in L2/3 of V1.
(g) Averaged response trace in (left) low (5-10%) or (right) high (60-80%) contrasts for $JAM2^{L23}$.
(h) Visual responses at various contrast levels in $JAM2^{L23}$ population during running (orange) or stationary (black) trials.
(i) Averaged response trace in (left) low (5-10%) or (right) high (60-80%) contrasts for $JAM2^{L23}$ during running (orange) or stationary (black) trials.
(j) Visual responses at various contrast levels in $JAM2^{L1}$ population labeled by *JAM-Cre* with AAV.Dlx.DIO.jGCaMP8m in L1 of V1.
(k) Averaged response trace in (left) low (5-10%) or (right) high (60-80%) contrasts for $JAM2^{L1}$.
(l) Visual responses at various contrast levels in $JAM2^{L1}$ population during running (purple) or stationary (black) trials.
(m) Averaged response trace in (left) low (5-10%) or (right) high (60-80%) contrasts for $JAM2^{L1}$ during running (purple) or stationary (black) trials.
(n-o) Visual responses at various contrast levels in (n) $VIP^{L23}$ (green) population labeled by *VIP-Cre* with AAV.Dlx.DIO.jGCaMP8m, (o) $SST^{L23}$ (blue) population labeled by *SST-Cre* with AAV.Dlx.DIO.jGCaMP8m.
(p) Ratio of high (60-80%) or low (5-10%) contrast preferring neurons in $JAM2^{L23}$ (orange), $JAM2^{L1}$ (purple), $VIP^{L23}$ (green), $SST^{L23}$ (blue). Each dot represents data from an individual animal.
(q) (left) Averaged response trace in 0% contrast ('blank trials') for $JAM2^{L1}$. (right)) Averaged response trace in 0% contrast ('blank trials') for $JAM2^{L1}$ during running (purple) or stationary (black) trials.
In (a-c,f,h,j,l,n,o), each gray line and dot represent averaged data from an animal, while color line represents mean and SEM from all neurons. Responses at 0% contrast ('blank trial') were indicated by dot. In (g,k,i,m), moving gratings were presented between 0-2 s indicated by dashed gray vertical lines. In (d,q), 0% contrast (the gray screen) was continually presented between 0-2 s indicated by dashed gray vertical lines. Data were averaged among neurons within each animal before plotting the line. The color line represents mean and SEM from all animals. Mann Whitney test (e,p) and mixed linear model regression (a-c,f,j,h,l,n,o) were used for testing statistical significance. See supplementary data 1 - Table 5.2 for statistics.

Extended Data 10 - related to Figure 6

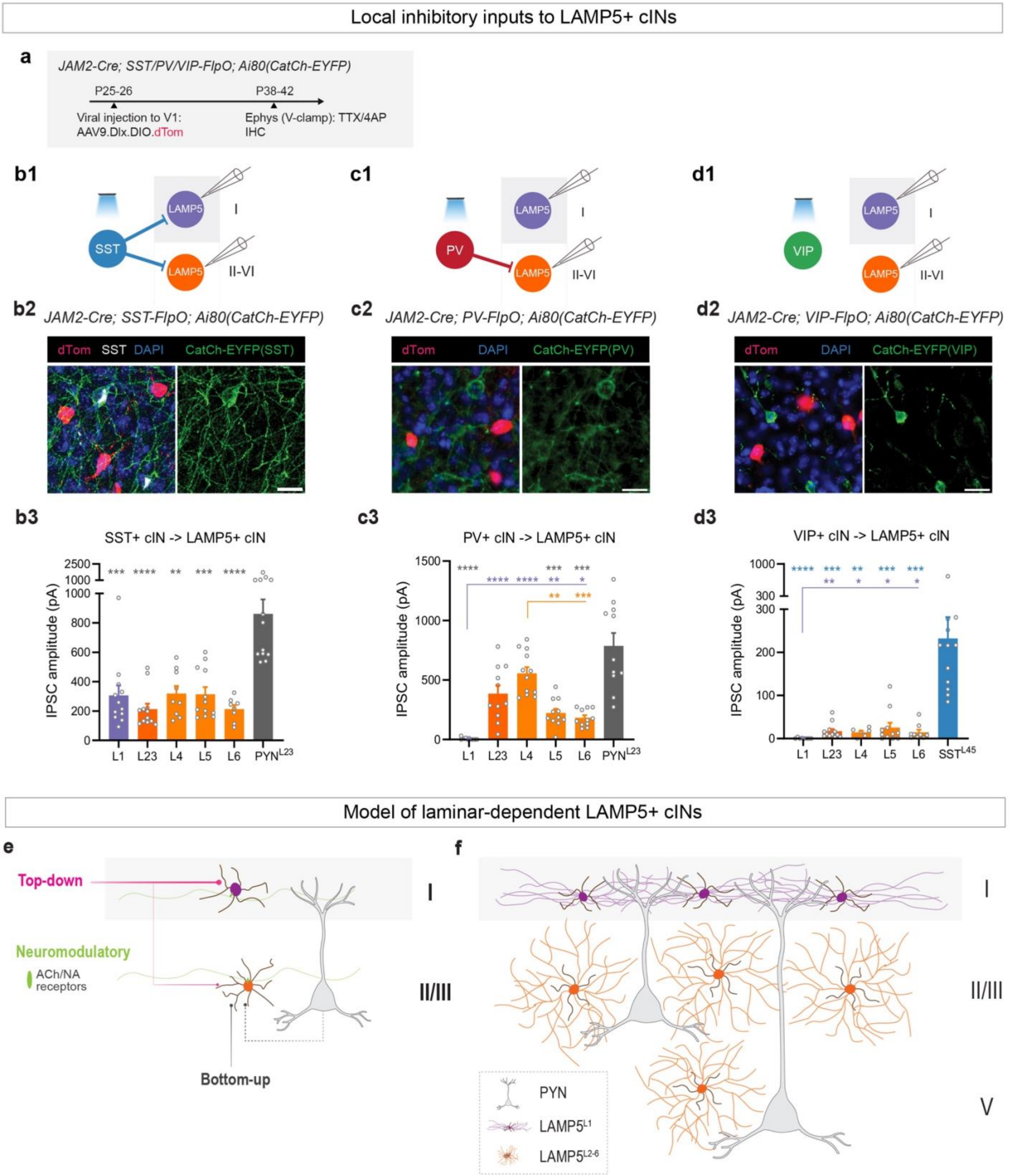

**Extended Data 10 (Related to Figure 6) - Local inhibitory inputs to LAMP5+ cINs in V1.**

(a) Experimental design for local inhibitory input mapping for LAMP5+ cINs in V1. *JAM2-Cre; SST/PV/VIP-FlpO; Ai80(CatCh-EYFP*) mice were injected with AAV9.Dlx.DIO.dTom to label LAMP5+ cINs in V1 at P25-26. Here, we used *JAM2-Cre* not only as a tool for targeting LAMP5+ cINs in adults using AAV **(Extended Data 2)**, but also as a *Germline-Cre* to convert Ai80 into a Flp-dependent reporter (see Method - Mouse). Mice were used for optogenetics-assisted circuit mapping with slice electrophysiology using voltage clamp with TTX and 4AP, and for immunohistochemistry at P38-42.
(b) Local inhibitory input from SST+ cINs to LAMP5+ cINs in V1. (b1) Experimental diagram for optogenetics-assisted circuit mapping with slice electrophysiology. (b2) Representative images for *JAM2-Cre; SST-FlpO; Ai80(CatCh-EYFP)* with AAV9.DLX.DIO.dTom in V1. (left) Overlay image with DAPI, dTom and CatCh-EYFP. LAMP5+ cINs expressed dTom and SST+ cINs expressed (right) CatCh-EYFP. Images were taken with confocal microscopy. (b3) Quantification of the IPSC amplitude for SST+ cIN input to LAMP5+ cINs across cortical layers in V1. SST+ cIN input to $PYN^{L23}$ was also recorded for comparison.
(c) Local inhibitory input from PV+ cINs to LAMP5+ cINs in V1. (c1) Experimental diagram for optogenetics-assisted circuit mapping with slice electrophysiology. (c2) Representative images for *JAM2-Cre; PV-FlpO; Ai80(CatCh-EYFP)*. Images were taken with wide-field fluorescence microscopy. (c3) Quantification of the IPSC amplitude for PV+ cIN input to LAMP5+ cINs across cortical layers in V1. PV+ cIN input to $PYN^{L23}$ was also recorded for comparison.
(d) Local inhibitory input from VIP+ cINs to LAMP5+ cINs in V1. (d1) Experimental diagram for optogenetics-assisted circuit mapping with slice electrophysiology. (d2) Representative images for *JAM2-Cre; VIP-FlpO; Ai80(CatCh-EYFP)*. Images were taken with wide-field fluorescence microscopy. (d3) Quantification of the IPSC amplitude for VIP+ cIN input to LAMP5+ cINs across cortical layers in V1. VIP+ cIN input to SST+ cINs was also recorded as the positive control. To label SST+ cINs, AAVPHP.s9e10.dTom (unpublished AAV enhancer to target SST+ cINs) was locally delivered to label SST+ cINs in L4/5 of V1.
(e) Abstract model of the laminar-dependent input connectivity of LAMP5+ cINs. Their dendrites were shown, while the axons were not shown.
(f) Model of the laminar-dependent output regulation from LAMP5+ cINs onto PYNs.
Scale bar = 20 μm. Error bar represents SEM. Kruskal-Wallis test followed by uncorrected Dunn's test were used for testing statistical significance. See supplementary data 1 - Table 6.1 for statistics.

**Supplementary data 1 - Statistics**

**Table 1 - related to Figure 1**

| | Mean ± SEM | P value | |
|---|---|---|---|
| Figure 1b:<br>V1 cell number | L1: 0.364 ± 0.015<br>L2/3: 0.130 ± 0.010<br>L4: 0.032 ± 0.006<br>L5: 0.033 ± 0.003<br>L6: 0.035 ± 0.002<br><br>*N = 3, n = 822 cells* | Repeated measures ANOVA:<br>p < 0.0001 (****) | Uncorrected Fisher's LSD:<br>L1 vs. L2/3: p = 0.0025 (**)<br>L1 vs. L4: p = 0.0008 (***)<br>L1 vs. L5: p = 0.0013 (**)<br>L1 vs. L6: p = 0.0017 (**)<br>L2/3 vs. L4: p = 0.0038 (**)<br>L2/3 vs. L5: p = 0.0079 (**)<br>L2/3 vs. L6: p = 0.0118 (*)<br>L4 vs. L5: p = 0.7266 (ns)<br>L4 vs. L6: p = 0.6449 (ns)<br>L5 vs. L6: p = 0.5200 (ns) |
| Figure 1c:<br>V1 neurite intensity | L1: 3130 ± 34.97<br>L2/3: 1746 ± 42.31<br>L4: 1455 ± 59.67<br>L5: 1124 ± 83.32<br>L6: 1270 ± 113.8<br><br>*N = 3, n = 6 slices* | Repeated measures ANOVA:<br>p = 0.0042 (**) | Uncorrected Fisher's LSD:<br>L1 vs. L2/3: p = 0.0028 (**)<br>L1 vs. L4: p = 0.0006 (***)<br>L1 vs. L5: p = 0.0032 (**)<br>L1 vs. L6: p = 0.0062 (**)<br>L2/3 vs. L4: p = 0.0681 (ns)<br>L2/3 vs. L5: p = 0.0044 (**)<br>L2/3 vs. L6: p = 0.0236 (*)<br>L4 vs. L5: p = 0.0937 (ns)<br>L4 vs. L6: p = 0.3476 (ns)<br>L5 vs. L6: p = 0.0867 (ns) |
| Figure 1e:<br>NPY<br>(N = 3) | L1: 0.796 ± 0.046<br>L2/3: 0.994 ± 0.006<br>L4: 1.000 ± 0.000<br>L5: 0.958 ± 0.042<br>L6: 0.928 ± 0.049<br><br>*N=3, n = 103 (L1), 157 (L2/3), 35 (L4), 27 (L5), 24 (L6) tdTom+ cells* | Mixed-effects model:<br>p = 0.0004 (***) | Uncorrected Fisher's LSD:<br>L1 vs. L2/3: p < 0.0001 (****)<br>L1 vs. L4: p < 0.0001 (****)<br>L1 vs. L5: p = 0.0025 (**)<br>L1 vs. L6: p = 0.0029 (**)<br>L2/3 vs. L4: p = 0.8683 (ns)<br>L2/3 vs. L5: p = 0.1988 (ns)<br>L2/3 vs. L6: p = 0.0953 (ns)<br>L4 vs. L5: p = 0.1539 (ns)<br>L4 vs. L6: p = 0.0703 (ns)<br>L5 vs. L6: p = 0.7561 (ns) |
| Figure 1f:<br>SV2C<br>(N = 3) | L1: 0.905 ± 0.021<br>L2/3: 0.978 ± 0.013<br>L4: 1.000 ± 0.000<br>L5: 1.000 ± 0.000<br>L6: 1.000 ± 0.000<br><br>*N=3, n = 164 (L1), 169 (L2/3), 26 (L4), 30 (L5), 54 (L6) tdTom+ cells* | Repeated measures ANOVA:<br>p < 0.0001 (****) | Uncorrected Fisher's LSD:<br>L1 vs. L2/3: p = 0.0002 (***)<br>L1 vs. L4: p < 0.0001 (****)<br>L1 vs. L5: p < 0.0001 (****)<br>L1 vs. L6: p < 0.0001 (****)<br>2/3 vs. L4: p = 0.2051 (ns)<br>2/3 vs. L5: p = 0.2051 (ns)<br>2/3 vs. L6: p = 0.2051 (ns)<br>4 vs. L5: p > 0.9999 (ns)<br>4 vs. L6: p > 0.9999 (ns)<br>5 vs. L6: p > 0.9999 (ns) |
| Figure 1g:<br>LSP1<br>(N = 3) | L1: 0.069 ± 0.020<br>L2/3: 0.491 ± 0.023<br>L4: 0.835 ± 0.060<br>L5: 0.781 ± 0.074<br>L6: 0.738 ± 0.068<br><br>*N=3, n = 162 (L1), 133 (L2/3), 39 (L4), 23 (L5), 52 (L6) tdTom+ cells* | Repeated measures ANOVA:<br>p < 0.0001 (****) | Uncorrected Fisher's LSD:<br>L1 vs. L2/3: p < 0.0001 (****)<br>L1 vs. L4: p < 0.0001 (****)<br>L1 vs. L5: p < 0.0001 (****)<br>L1 vs. L6: p < 0.0001 (****)<br>L2/3 vs. L4: p = 0.0016 (**)<br>L2/3 vs. L5: p = 0.0111 (*)<br>L2/3 vs. L6: p = 0.0241 (*)<br>L4 vs. L5: p = 0.6431 (ns)<br>L4 vs. L6: p = 0.2183 (ns)<br>L5 vs. L6: p = 0.6467 (ns) |
| Figure 1h:<br>NDNF<br>(N = 3) | L1: 0.854 ± 0.023<br>L2/3: 0.097 ± 0.035<br>L4: 0.000 ± 0.000<br>L5: 0.042 ± 0.042<br>L6: 0.061 ± 0.039<br><br>*N=3, n = 154 (L1), 115 (L2/3), 30 (L4), 30 (L5), 39 (L6) tdTom+ cells* | Repeated measures ANOVA:<br>p < 0.0001 (****) | Uncorrected Fisher's LSD:<br>L1 vs. L2/3: p < 0.0001 (****)<br>L1 vs. L4: p < 0.0001 (****)<br>L1 vs. L5: p < 0.0001 (****)<br>L1 vs. L6: p < 0.0001 (****)<br>L2/3 vs. L4: p = 0.0476 (*)<br>L2/3 vs. L5: p = 0.2424 (ns)<br>L2/3 vs. L6: p = 0.4433 (ns)<br>L4 vs. L5: p = 0.3759 (ns)<br>L4 vs. L6: p = 0.1990 (ns)<br>L5 vs. L6: p = 0.6771 (ns) |

**Table 1.1 - related to Extended Data 1 - Figure 1**

| | Mean ± SEM | P value | |
|---|---|---|---|
| Ex 1b3:<br>Gad2<br>expression | Lamp5: 3465.267 ± 54.847<br>Pvalb: 1091.106 ± 19.346<br>Sst: 1153.380 ± 18.853<br>Vip: 1382.798 ± 25.582<br>Serpinf1: 1988.185 ± 305.195<br>Sncg: 985.960 ± 57.597 | | Wilcoxon signed-rank test:<br>Lamp5 vs. Pvalb: p < 0.0001 (****)<br>Lamp5 vs. Sst: p < 0.0001 (****)<br>Lamp5 vs. Vip: p < 0.0001 (****)<br>Lamp5 vs. Serpinf1: p < 0.0001 (****)<br>Lamp5 vs. Sncg: p < 0.0001 (****) |
| Ex 1c1:<br>SV2C for<br>robustness | L1: 0.990 ± 0.010<br>L2/3: 0.996 ± 0.004<br>L4: 1.000 ± 0.000<br><br>*N=3, n = 149 (L1), 166 (L2/3), 26 (L4) SV2C+ cells* | Repeated measures ANOVA:<br>p = 0.5696 (ns) | |
| Ex 1c2:<br>NDNF for<br>robustness<br>(N = 3) | L1: 0.920 ± 0.017<br>L2/3: 1.000 ± 0.000<br><br>*N=3, n = 141 (L1), 11 (L2/3) NDNF+ cells* | | Wilcoxon signed rank test:<br>p = 0.1250 (ns) |
| Ex 1d:<br>DLX5<br>(N = 3) | L1: 0.991 ± 0.009<br>L2/3: 0.988 ± 0.012<br>L4: 1.000 ± 0.000<br>L5: 0.958 ± 0.042<br>L6: 0.972 ± 0.028<br><br>*N=3, n = 127 (L1), 100 (L2/3), 31 (L4), 19 (L5), 32 (L6) tdTom+ cells* | Repeated measures ANOVA:<br>p = 0.6901 (ns) | |
| Ex 1e:<br>LHX6 | L1: 0.000 ± 0.000<br>L2/3: 0.010 ± 0.010<br>L4: 0.000 ± 0.000<br>L5: 0.083 ± 0.083<br>L6: 0.037 ± 0.037<br><br>*N=3, n = 89 (L1), 71 (L2/3), 11 (L4), 11 (L5), 17 (L6) tdTom+ cells* | Mixed-effects model:<br>p = 0.3529 (ns) | |
| Ex 1f:<br>VIP<br>(N = 3) | L1: 0.000 ± 0.000<br>L2/3: 0.000 ± 0.000<br>L4: 0.000 ± 0.000<br>L5: 0.042 ± 0.042<br>L6: 0.000 ± 0.000<br><br>*N=3, n = 146 (L1), 164 (L2/3), 32 (L4), 29 (L5), 50 (L6) tdTom+ cells* | Repeated measures ANOVA:<br>p = 0.4241 (ns) | |
| Ex 1g:<br>SNCG<br>(N = 3) | L1: 0.000 ± 0.000<br>L2/3: 0.000 ± 0.000<br>L4: 0.000 ± 0.000<br>L5: 0.021 ± 0.021<br>L6: 0.000 ± 0.000<br><br>*N=3, n = 181 (L1), 158 (L2/3), 34 (L4), 29 (L5), 39 (L6) tdTom+ cells* | Repeated measures ANOVA:<br>p = 0.4241 (ns) | |
| Ex 1i:<br>S1 cell<br>number<br>(N = 3) | L1: 0.293 ± 0.034<br>L2/3: 0.130 ± 0.011<br>L4: 0.048 ± 0.002<br>L5: 0.046 ± 0.005<br>L6: 0.056 ± 0.007<br><br>*N = 3, n = 1183 cells* | Repeated measures ANOVA:<br>F = 46.74, p = 0.0109 (*) | Uncorrected Fisher's LSD:<br>L1 vs. L2/3: p = 0.0486 (*)<br>L1 vs. L4: p = 0.0174 (*)<br>L1 vs. L5: p = 0.0154 (*)<br>L1 vs. L6: p = 0.0130 (*)<br>L2/3 vs. L4: p = 0.0199 (*)<br>L2/3 vs. L5: p = 0.0313 (*)<br>L2/3 vs. L6: p = 0.0258 (*)<br>L4 vs. L5: p = 0.6141 (ns)<br>L4 vs. L6: p = 0.3297 (ns)<br>L5 vs. L6: p = 0.2770 (ns) |
| Ex 1j:<br>S1 neurite<br>intensity<br>(N = 3) | L1: 2935 ± 71.73<br>L2/3: 2030 ± 8.518<br>L4: 1752 ± 35.29<br>L5: 1387 ± 15.56<br>L6: 1246 ± 23.88<br><br>*N = 3, n = 6 slices* | Repeated measures ANOVA:<br>F = 267.0, p = 0.0025 (**) | Uncorrected Fisher's LSD:<br>L1 vs. L2/3: p = 0.0065 (**)<br>L1 vs. L4: p = 0.0067 (**)<br>L1 vs. L5: p = 0.0029 (**)<br>L1 vs. L6: p = 0.0032 (**)<br>L2/3 vs. L4: p = 0.0101 (*)<br>L2/3 vs. L5: p = 0.0003 (***)<br>L2/3 vs. L6: p = 0.0008 (***)<br>L4 vs. L5: p = 0.0031 (**)<br>L4 vs. L6: p = 0.0023 (**)<br>L5 vs. L6: p = 0.0090 (**) |
| Ex 1l:<br>M1 cell | L1: 0.261 ± 0.026<br>L2/3: 0.137 ± 0.008 | Repeated measures ANOVA:<br>F = 51.47, p = 0.0151 (*) | Uncorrected Fisher's LSD:<br>L1 vs. L2/3: p = 0.0413 (*) |

| | | | |
|---|---|---|---|
| number<br>(N = 3) | L5: 0.038 ± 0.006<br>L6: 0.052 ± 0.002<br><br>*N = 3, n = 997 cells* | | L1 vs. L5: p = 0.0178 (*)<br>L1 vs. L6: p = 0.0168 (*)<br>L2/3 vs. L5: p = 0.0034 (**)<br>L2/3 vs. L6: p = 0.0070 (**)<br>L5 vs. L6: p = 0.0830 (ns) |
| Ex 1m:<br>M1 cell<br>number<br>(N = 3) | L1: 2786 ± 185.1<br>L2/3: 1874 ± 199.1<br>L5: 1340 ± 135.0<br>L6: 1169 ± 129.4<br><br>*N = 3, n = 6 slices* | Repeated measures ANOVA:<br>F = 227.4, p = 0.0016 (**) | Uncorrected Fisher's LSD:<br>L1 vs. L2/3: p = 0.0019 (**)<br>L1 vs. L5: p = 0.0027 (**)<br>L1 vs. L6: p = 0.0033 (**)<br>L2/3 vs. L5: p = 0.0165 (*)<br>L2/3 vs. L6: p = 0.0137 (*)<br>L5 vs. L6: p = 0.0103 (*) |

**Table 2 - related to Figure 2**

| | Mean ± SEM | P value |
|---|---|---|
| Figure 2e: Ratio of correlated pairs | LAMP5::CTR: 0.321 ± 0.037<br>LAMP5::TOX: 0.417 ± 0.018<br><br>*LAMP5::CTR: N = 3, n = 39443 pairs*<br>*LAMP5::TOX: N = 5, n = 95227 pairs* | Mann-Whitney Test: p = 0.0357 (*) |
| Figure 2f: Pearson's r (All Pairs) | LAMP5::CTR: 0.0159 ± 0.0028<br>LAMP5::TOX: 0.0348 ± 0.0038<br><br>*LAMP5::CTR: N = 3, n = 39443 pairs*<br>*LAMP5::TOX: N = 5, n = 95227 pairs* | Hierarchical Bootstrap: p < 0.0001 (****)<br><br>Mixed Linear Model Regression: p = 0.001 (**)<br><br>Mann-Whitney Test: p = 0.0357 (*) |
| Figure 2h: Diff (run - stationary) | LAMP5::CTR: 0.211 ± 0.010<br>LAMP5::TOX: 0.391 ± 0.063<br><br>*LAMP5::CTR: N = 3, n = 622 cells*<br>*LAMP5::TOX: N = 5, n = 1151 cells* | Hierarchical Bootstrap: p = 0.0016 (**)<br><br>Mixed Linear Model Regression: p = 0.035 (*)<br><br>Mann-Whitney Test: p = 0.0357 (*) |

**Table 2.1 - related to Extended Data 3 - Figure 2**

| | Mean ± SEM | P value |
|---|---|---|
| Ex 3a:<br>Pearson's r<br>(p<0.05): | LAMP5::CTR: 0.0398 ± 0.0022<br>LAMP5::TOX: 0.0732 ± 0.0096<br><br>*LAMP5::CTR: N = 3, n = 13387 pairs*<br>*LAMP5::TOX: N = 5, n = 40182 pairs* | Hierarchical Bootstrap: p < 0.0001 (****)<br><br>Mixed Linear Model Regression: p = 0.010 (*)<br><br>Mann-Whitney Test: p = 0.0357 (*) |
| Ex 3b2:<br>Stationary and Running | LAMP5::CTR (stationary): 0.061 ± 0.023<br>LAMP5::TOX (stationary): 0.049 ± 0.012<br>LAMP5::CTR (run): 0.272 ± 0.026<br>LAMP5::TOX (run): 0.439 ± 0.073<br><br>*LAMP5::CTR: N = 3, n = 622 cells*<br>*LAMP5::TOX: N = 5, n = 1151 cells* | Hierarchical Bootstrap:<br>LAMP5::CTR (stationary) vs. LAMP5::TOX (stationary): p = 0.258 (ns)<br>LAMP5::CTR (run) vs. LAMP5::TOX (run): p = 0.009 (**)<br><br>Mann-Whitney Test:<br>LAMP5::CTR (stationary) vs. LAMP5::TOX (stationary): p = 0.5714 (ns)<br>LAMP5::CTR (run) vs. LAMP5::TOX (run): p = 0.0357 (*) |
| Ex 3g:<br>VIP Ratio of Correlated Pairs | VIP::CTR: 0.256 ± 0.018<br>VIP::TOX: 0.255 ± 0.010<br><br>*VIP::CTR: N = 3, n = 57452 pairs*<br>*VIP::TOX: N = 5, n = 174848 pairs* | Mann-Whitney Test: p > 0.9999 (ns) |
| Ex 3h:<br>VIP Pearson's r (All Pairs) | VIP::CTR: 0.017 ± 0.006<br>VIP::TOX: 0.013 ± 0.002<br><br>*VIP::CTR: N = 3, n = 57452 pairs*<br>*VIP::TOX: N = 5, n = 174848 pairs* | Hierarchical Bootstrap: p = 0.28 (ns)<br><br>Mixed Linear Model Regression: p = 0.451 (ns)<br><br>Mann-Whitney Test: p = 0.7857 (ns) |
| Ex 3i:<br>VIP Pearson's r (p < 0.05) | VIP::CTR: 0.052 ± 0.014<br>VIP::TOX: 0.043 ± 0.006<br><br>*VIP::CTR: N = 3, n = 15420 pairs*<br>*VIP::TOX: N = 5, n = 45212 pairs* | Hierarchical Bootstrap: p = 0.23 (ns)<br><br>Mixed Linear Model Regression: p = 0.476 (ns)<br><br>Mann-Whitney Test: p = 0.7857 (ns) |
| Ex 3k:<br>VIP Stationary and Running | VIP::CTR (stationary): 0.068 ± 0.017<br>VIP::TOX (stationary): 0.081 ± 0.007<br>VIP::CTR (run): 0.265 ± 0.049<br>VIP::TOX (run): 0.230 ± 0.017<br><br>*VIP::CTR: N = 3, n = 675 cells*<br>*VIP::TOX: N = 5, n = 1623 cells* | Hierarchical Bootstrap<br>VIP::CTR (stationary) vs. VIP::TOX (stationary): p = 0.078 (ns)<br>VIP::CTR (run) vs. VIP::TOX (run): p = 0.492 (ns)<br><br>Mann-Whitney Test<br>VIP::CTR (stationary) vs. VIP::TOX (stationary): p = 0.5714 (ns)<br>VIP::CTR (run) vs. VIP::TOX (run): p = 0.7857 (ns) |
| Ex 3l:<br>VIP Diff (run - stationary) | VIP::CTR: 0.197 ± 0.045<br>VIP::TOX: 0.149 ± 0.014<br><br>*VIP::CTR: N = 3, n = 675 cells*<br>*VIP::TOX: N = 5, n = 1623 cells* | Hierarchical Bootstrap: p = 0.294 (ns)<br><br>Mixed Linear Model Regression: p = 0.263 (ns)<br><br>Mann-Whitney Test: p = 0.5714 (ns) |

**Table 3 - related to Figure 3**

| | Mean ± SEM | P value |
|---|---|---|
| Figure 3d: Pseudo-orientation | Orientation 0°<br>LAMP5::CTR: 0.057 ± 0.018<br>LAMP5::TOX: 0.040 ± 0.023<br>Orientation 30°<br>LAMP5::CTR: 0.052 ± 0.031<br>LAMP5::TOX: 0.063 ± 0.039<br>Orientation 60°<br>LAMP5::CTR: 0.187 ± 0.039<br>LAMP5::TOX: 0.139 ± 0.050<br>Orientation 90°<br>LAMP5::CTR: 0.936 ± 0.077<br>LAMP5::TOX: 0.667 ± 0.097<br>Orientation 120°<br>LAMP5::CTR: 0.198 ± 0.041<br>LAMP5::TOX: 0.138 ± 0.052<br>Orientation 150°<br>LAMP5::CTR: 0.091 ± 0.020<br>LAMP5::TOX: 0.056 ± 0.026<br>Orientation 180°<br>LAMP5::CTR: 0.051 ± 0.022<br>LAMP5::TOX: 0.039 ± 0.028<br>Orientation 210°<br>LAMP5::CTR: 0.066 ± 0.030<br>LAMP5::TOX: 0.038 ± 0.038<br>Orientation 240°<br>LAMP5::CTR: 0.180 ± 0.042<br>LAMP5::TOX: 0.126 ± 0.053<br>Orientation 270°<br>LAMP5::CTR: 0.424 ± 0.054<br>LAMP5::TOX: 0.228 ± 0.069<br>Orientation 300°<br>LAMP5::CTR: 0.173 ± 0.023<br>LAMP5::TOX: 0.110 ± 0.030<br>Orientation 330°<br>LAMP5::CTR: 0.069 ± 0.016<br>LAMP5::TOX: 0.037 ± 0.020<br><br>*LAMP5::CTR: N = 3, n = 188 neurons*<br>*LAMP5::TOX: N = 5, n = 318 neurons* | Hierarchical Bootstrap:<br>Orientation 0°: p = 0.275 (ns)<br>Orientation 30°: p = 0.292 (ns)<br>Orientation 60°: p = 0.181 (ns)<br>Orientation 90°: p = 0.0005 (***)<br>Orientation 120°: p = 0.150 (ns)<br>Orientation 150°: p = 0.115 (ns)<br>Orientation 180°: p = 0.400 (ns)<br>Orientation 210°: p = 0.273 (ns)<br>Orientation 240°: p = 0.162 (ns)<br>Orientation 270°: p = 0.0013 (**)<br>Orientation 300°: p = 0.040 (*)<br>Orientation 330°: p = 0.0899 (ns)<br><br>Mixed Linear Model Regression:<br>Orientation 0°: p = 0.462 (ns)<br>Orientation 30°: p = 0.772 (ns)<br>Orientation 60°: p = 0.331 (ns)<br>Orientation 90°: p = 0.006 (**)<br>Orientation 120°: p = 0.251 (ns)<br>Orientation 150°: p = 0.172 (ns)<br>Orientation 180°: p = 0.661 (ns)<br>Orientation 210°: p = 0.466 (ns)<br>Orientation 240°: p = 0.308 (ns)<br>Orientation 270°: p = 0.004 (**)<br>Orientation 300°: p = 0.033 (*)<br>Orientation 330°: p = 0.111 (ns) |
| Figure 3e: Signal-to-Noise Ratio (SNR) | LAMP5::CTR: 0.388 ± 0.038<br>LAMP5::TOX: 0.279 ± 0.048<br><br>*LAMP5::CTR: N = 3, n = 188 neurons*<br>*LAMP5::TOX: N = 5, n = 318 neurons* | Hierarchical Bootstrap: p = 0.008 (**)<br>Mixed Linear Model Regression: p = 0.022 (*) |
| Figure 3f: gOSI | LAMP5::CTR: 0.413 ± 0.034<br>LAMP5::TOX: 0.340 ± 0.043<br><br>*LAMP5::CTR: N = 3, n = 188 neurons*<br>*LAMP5::TOX: N = 5, n = 318 neurons* | Hierarchical Bootstrap: p = 0.0182 (*)<br>Mixed Linear Model Regression: p = 0.089 (ns) |

**Table 3.1 - related to Extended Data 4 - Figure 3**

| | Mean ± SEM | P value |
|---|---|---|
| Ex 4a:<br>Responsive Ratio | LAMP5::CTR: 0.272 ± 0.051<br>LAMP5::TOX: 0.310 ± 0.048<br><br>*LAMP5::CTR: N = 3, n = 651 neurons*<br>*LAMP5::TOX: N = 5, n = 1152 neurons* | Mann-Whitney Test: p = 0.5714 (ns) |
| Ex 4b:<br>Peak Response | LAMP5::CTR: 0.936 ± 0.077<br>LAMP5::TOX: 0.667 ± 0.097<br><br>*LAMP5::CTR: N = 3, n = 188 neurons*<br>*LAMP5::TOX: N = 5, n = 318 neurons* | Hierarchical Bootstrap: p = 0.0005 (***)<br>Mixed Linear Model Regression: p = 0.006 (**) |
| Ex 4c-left:<br>Mean Standardized dF/F0 During Trial Baseline (Pre-Stimulus) | LAMP5::CTR: 0.003 ± 0.025<br>LAMP5::TOX: 0.074 ± 0.031<br><br>*LAMP5::CTR: N = 3, n = 188 neurons*<br>*LAMP5::TOX: N = 5, n = 318 neurons* | Hierarchical Bootstrap: p = 0.024 (*)<br>Mixed Linear Model Regression: p = 0.023 (*) |
| Ex 4c-right:<br>Mean Standardized dF/F0 During Visual Stimulus | LAMP5::CTR: 0.939 ± 0.063<br>LAMP5::TOX: 0.74 ± 0.08<br><br>*LAMP5::CTR: N = 3, n = 188 neurons*<br>*LAMP5::TOX: N = 5, n = 318 neurons* | Hierarchical Bootstrap: p = 0.003 (**)<br>Mixed Linear Model Regression: p = 0.013 (*) |
| Ex 4d:<br>Signal Correlation | LAMP5::CTR: 0.252 ± 0.095<br>LAMP5::TOX: 0.230 ± 0.120<br><br>*LAMP5::CTR: N = 3, n = 188 neurons*<br>*LAMP5::TOX: N = 5, n = 318 neurons* | Hierarchical Bootstrap: p = 0.379 (ns)<br>Mixed Linear Model Regression: p = 0.852 (ns) |
| Ex 4e:<br>Noise Correlation | LAMP5::CTR: 0.123 ± 0.046<br>LAMP5::TOX: 0.129 ± 0.058<br><br>*LAMP5::CTR: N = 3, n = 188 neurons*<br>*LAMP5::TOX: N = 5, n = 318 neurons* | Hierarchical Bootstrap: p = 0.408 (ns)<br>Mixed Linear Model Regression: p = 0.920 (ns) |
| Ex 4f:<br>Preferred Orientation | LAMP5::CTR: 94.204 ± 7.767<br>LAMP5::TOX: 92.504 ± 9.844<br><br>*LAMP5::CTR: N = 3, n = 188 neurons*<br>*LAMP5::TOX: N = 5, n = 318 neurons* | Hierarchical Bootstrap: p = 0.494 (ns)<br>Mixed Linear Model Regression: p = 0.863 (ns) |
| Ex 4g:<br>Direction Selectivity Index (DSI) | LAMP5::CTR: 0.372 ± 0.023<br>LAMP5::TOX: 0.435 ± 0.029<br><br>*LAMP5::CTR: N = 3, n = 188 neurons*<br>*LAMP5::TOX: N = 5, n = 318 neurons* | Hierarchical Bootstrap: p = 0.053 (ns)<br>Mixed Linear Model Regression: p = 0.032 (*) |

**Table 4 - related to Figure 4**

| | Mean ± SEM | P value | |
|---|---|---|---|
| Figure 4e:<br>mean activity<br>(stationary) | LAMP5$^{L23}$: 0.062 ± 0.016<br>LAMP5$^{L1}$: 0.106 ± 0.011<br><br>*LAMP5$^{L23}$: N = 8, n = 160 neurons*<br>*LAMP5$^{L1}$: N = 8, n = 228 neurons* | Mixed Linear Model Regression: $p < 0.0001$ (****)<br>Hierarchical bootstrap: p = 0.008 (**) | Mann Whitney test:<br>LAMP5$^{L23}$: $p < 0.0001$ (****)<br>LAMP5$^{L1}$: $p < 0.0001$ (****) |
| Figure 4e:<br>mean activity (run) | LAMP5$^{L23}$: 0.971 ± 0.092<br>LAMP5$^{L1}$: 0.804 ± 0.057<br><br>*LAMP5$^{L23}$: N = 8, n = 160 neurons*<br>*LAMP5$^{L1}$: N = 8, n = 228 neurons* | Mixed Linear Model Regression: p = 0.004 (**)<br>Hierarchical bootstrap: p = 0.146 (ns) | |
| Figure 4f:<br>Ratio of Increased<br>Activity During Run | LAMP5$^{L23}$: 0.964 ± 0.018<br>LAMP5$^{L1}$: 0.792 ± 0.043<br><br>*LAMP5$^{L23}$: N = 8, n = 160 neurons*<br>*LAMP5$^{L1}$: N = 8, n = 228 neurons* | Mann Whitney test: p = 0.0012 (**) | |
| Figure 4k:<br>Ratio of Significant<br>Correlation between<br>activity and speed | LAMP5$^{L23}$: 0.886 ± 0.029<br>LAMP5$^{L1}$: 0.667 ± 0.040<br><br>*LAMP5$^{L23}$: N = 7, n = 125 neurons*<br>*LAMP5$^{L1}$: N = 7, n = 200 neurons* | Mann Whitney test: p = 0.0012 (**) | |
| Figure 4l:<br>Pearson's<br>Correlation between<br>activity and speed | LAMP5$^{L23}$: 0.370 ± 0.041<br>LAMP5$^{L1}$: 0.249 ± 0.027<br><br>*LAMP5$^{L23}$: N = 7, n = 125 neurons*<br>*LAMP5$^{L1}$: N = 7, n = 200 neurons* | Hierarchical bootstrap: p = 0.007 (**)<br>Mixed linear model regression: $p < 0.0001$ (****) | |

**Table 4.1 - related to Extended Data 5 - Figure 4**

| | Mean ± SEM | P value |
|---|---|---|
| Ex 5c:<br>Ratio of significant correlation of neuronal pairs | $LAMP5^{L23}$: 0.857 ± 0.043<br>$LAMP5^{L1}$: 0.623 ± 0.051<br><br>*$LAMP5^{L23}$: N = 8, n = 991 pairs*<br>*$LAMP5^{L1}$: N = 8, n = 1822 pairs* | Mann Whitney test: p = 0.003 (**) |
| Ex 5d:<br>Pearson's correlation of neuronal pairs | $LAMP5^{L23}$: 0.383 ± 0.034<br>$LAMP5^{L1}$: 0.213 ± 0.010<br><br>*$LAMP5^{L23}$: N = 8, n = 991 pairs*<br>*$LAMP5^{L1}$: N = 8, n = 1822 pairs* | Hierarchical Bootstrap: p < 0.0001 (****)<br>Mixed Linear Model Regression: p < 0.0001 (****) |
| Ex 5g:<br>Mean activity with dark screen (Stationary) | $LAMP5^{L23}$: 0.051 ± 0.014<br>$LAMP5^{L1}$: 0.076 ± 0.010<br><br>*$LAMP5^{L23}$: N = 8, n = 172 neurons*<br>*$LAMP5^{L1}$: N = 8, n = 252 neurons* | Mixed Linear Model Regression: p = 0.017 (*)<br>Hierarchical Bootstrap: p = 0.090 (ns) |
| Ex 5g:<br>Mean activity with dark screen (Run) | $LAMP5^{L23}$: 0.991 ± 0.089<br>$LAMP5^{L1}$: 0.840 ± 0.054<br><br>*$LAMP5^{L23}$: N = 8, n = 172 neurons*<br>*$LAMP5^{L1}$: N = 8, n = 252 neurons* | Mixed Linear Model Regression: p = 0.006 (**)<br>Hierarchical Bootstrap: p = 0.109 (ns) |
| Ex 5h:<br>Ratio of increased activity during run in dark vs gray screen | Gray ($LAMP5^{L23}$): 0.964 ± 0.018<br>Dark ($LAMP5^{L23}$): 0.950 ± 0.013<br><br>Gray ($LAMP5^{L1}$): 0.792 ± 0.043<br>Dark ($LAMP5^{L1}$): 0.824 ± 0.041<br><br>*$LAMP5^{L23}$: N = 8, n = 160 neurons (gray), n = 172 neurons (dark)*<br>*$LAMP5^{L1}$: N = 8, n = 228 neurons (gray), n = 252 neurons (dark)* | ($LAMP5^{L23}$) Mann-Whitney test: p = 0.3350<br><br>($LAMP5^{L1}$) Mann-Whitney test: p = 0.7209 |
| Ex 5j:<br>Ratio of significant correlation with dark screen | Gray ($LAMP5^{L23}$): 0.886 ± 0.029<br>Dark ($LAMP5^{L23}$): 0.862 ± 0.037<br><br>Gray ($LAMP5^{L1}$): 0.667 ± 0.040<br>Dark ($LAMP5^{L1}$): 0.709 ± 0.036<br><br>*$LAMP5^{L23}$: N = 7, n = 125 neurons (gray), n = 133 neurons (dark)*<br>*$LAMP5^{L1}$: N = 7, n = 200 neurons (gray), n = 207 neurons (dark)* | ($LAMP5^{L23}$) Mann-Whitney test: p = 0.5991 (ns)<br><br>($LAMP5^{L1}$) Mann-Whitney test: p = 0.3345 (ns) |
| Ex 5k:<br>Pearson’s correlation coefficient with dark screen | Gray ($LAMP5^{L23}$): 0.367 ± 0.026<br>Dark ($LAMP5^{L23}$): 0.338 ± 0.042<br><br>Gray ($LAMP5^{L1}$): 0.246 ± 0.038<br>Dark ($LAMP5^{L1}$): 0.250 ± 0.024<br><br>*$LAMP5^{L23}$: N = 7, n = 125 neurons (gray), n = 133 neurons (dark)*<br>*$LAMP5^{L1}$: N = 7, n = 200 neurons (gray), n = 207 neurons (dark)* | ($LAMP5^{L23}$) Mann-Whitney test: p = 0.7104 (ns)<br><br>($LAMP5^{L1}$) Mann-Whitney test: p = 0.7104 (ns) |

**Table 4.2 - related to Extended Data 6 - Figure 4**

| | Mean ± SEM | P value |
|---|---|---|
| Ex 6d:<br>mean activity (stationary) with dark screen | $JAM2^{L23}$: 0.001 ± 0.016<br>$JAM2^{L1}$: 0.032 ± 0.012<br><br>$JAM2^{L23}$: N = 9, n = 163 neurons<br>$JAM2^{L1}$: N = 11, n = 170 neurons | Mixed Linear Model Regression: p = 0.007 (**)<br>Hierarchical bootstrap: p = 0.051 (ns) |
| Ex 6d:<br>mean activity (run) with dark screen | $JAM2^{L23}$: 0.635 ± 0.087<br>$JAM2^{L1}$: 0.601 ± 0.051<br><br>*$JAM2^{L23}$: N = 9, n = 163 neurons*<br>*$JAM2^{L1}$: N = 11, n = 170 neurons* | Mixed Linear Model Regression: p = 0.502 (ns)<br>Hierarchical bootstrap: p = 0.336 (ns) |
| Ex 6f3:<br>Ratio of significant correlation of neuronal pairs | $LAMP5^{L23}$: 0.821 ± 0.048<br>$LAMP5^{L1}$: 0.679 ± 0.030<br><br>*$JAM2^{L23}$: N = 9, n = 302 pairs*<br>*$JAM2^{L1}$: N = 11, n = 739 pairs* | Mann Whitney test: p = 0.0092 (**) |
| Ex 6f4:<br>Pearson's correlation of neuronal pairs (only significant pairs) | $LAMP5^{L23}$: 0.316 ± 0.025<br>$LAMP5^{L1}$: 0.225 ± 0.018<br><br>*$JAM2^{L23}$: N = 9, n = 251 pairs*<br>*$JAM2^{L1}$: N = 11, n = 550 pairs* | Hierarchical Bootstrap: p < 0.0001 (****)<br>Mixed Linear Model Regression: p < 0.0001 (****) |

**Table 4.3 - related to Extended Data 7 - Figure 4**

| | Mean ± SEM | P value |
|---|---|---|
| Ex 7d:<br>Mean activity (stationary) | $VIP^{L23}$: -0.028 ± 0.021<br>$SST^{L23}$: 0.097 ± 0.014<br>$PYN^{L23}$: 0.062 ± 0.015<br><br>*$VIP^{L23}$: N = 9, n = 205 neurons*<br>*$SST^{L23}$: N = 7, n = 80 neurons*<br>*$PYN^{L23}$: N = 6, n = 1297 neurons* | Mann Whitney test:<br>$VIP^{L23}$: p < 0.0001 (****)<br>$SST^{L23}$: p = 0.0023 (**)<br>$PYN^{L23}$: p = 0.0079 (**) |
| Ex 7d:<br>Mean activity (run) | $VIP^{L23}$: 0.747 ± 0.113<br>$SST^{L23}$: 0.356 ± 0.062<br>$PYN^{L23}$: 0.268 ± 0.033<br><br>*$VIP^{L23}$: N = 9, n = 205 neurons*<br>*$SST^{L23}$: N = 7, n = 80 neurons*<br>*$PYN^{L23}$: N = 6, n = 1297 neurons* | |
| Ex 7f:<br>Pearson's correlation coefficient | $LAMP5^{L23}$: 0.370 ± 0.041<br>$LAMP5^{L1}$: 0.249 ± 0.027<br>$VIP^{L23}$: 0.401 ± 0.061<br>$SST^{L23}$: 0.135 ± 0.069<br><br>*$LAMP5^{L23}$: N = 7, n = 125 neurons*<br>*$LAMP5^{L1}$: N = 7, n = 200 neurons*<br>*$VIP^{L23}$: N = 8, n = 89 neurons*<br>*$SST^{L23}$: N = 6, n = 37 neurons* | Mixed Linear Model Regression:<br>$LAMP5^{L23}$ vs $LAMP5^{L1}$: p < 0.0001 (****)<br>$LAMP5^{L23}$ vs $VIP^{L23}$: p = 0.614 (ns)<br>$LAMP5^{L23}$ vs $SST^{L23}$: p = 0.0010 (**)<br>$LAMP5^{L1}$ vs $VIP^{L23}$: p = 0.0110 (*)<br>$LAMP5^{L1}$ vs $SST^{L23}$: p = 0.096 (ns)<br>$VIP^{L23}$ vs $SST^{L23}$: p < 0.0001 (****) |

**Table 5 - related to Figure 5**

| | Mean ± SEM | P value | |
|---|---|---|---|
| Figure 5b:<br>Ratio of contrast preference | Ratio preferring high contrasts vs low contrasts:<br>LAMP5$^{L23}$: 0.537 vs 0.462 ± 0.065<br>LAMP5$^{L1}$: 0.240 vs 0.244 ± 0.070<br>VIP$^{L23}$: 0.133 vs 0.867 ± 0.035<br>SST$^{L23}$: 0.876 vs 0.124 ± 0.067<br>PYN$^{L23}$: 0.510 vs 0.490 ± 0.022<br><br>*LAMP5$^{L23}$: N = 8, n = 180 neurons*<br>*LAMP5$^{L1}$: N = 8, n = 251 neurons*<br>*VIP$^{L23}$: N = 9, n = 228 neurons*<br>*SST$^{L23}$: N = 7, n = 55 neurons*<br>*PYN$^{L23}$: N = 6, n = 893 neurons* | Mann Whitney test:<br>LAMP5$^{L23}$: p = 0.245 (ns)<br>LAMP5$^{L1}$: p = 0.0003 (***)<br>VIP$^{L23}$: p < 0.0001 (****)<br>SST$^{L23}$: p = 0.0012 (**)<br>PYN$^{L23}$: p = 0.4719 (ns) | |
| Figure 5d:<br>LAMP5$^{L23}$ contrast | 0%: -0.024 ± 0.022<br>5%: -0.024 ± 0.008<br>10%: -0.007 ± 0.007<br>20%: 0.002 ± 0.007<br>40%: -0.007 ± 0.007<br>60%: -0.015 ± 0.008<br>80%: 0.011 ± 0.004<br><br>*LAMP5$^{L23}$: N = 8, n = 180 neurons* | Mixed Linear Model Regression:<br>0% vs 5%: p = 0.554 (ns), **0% vs 10%: p = 0.049 (*),**<br>**0% vs 20%: p = 0.004 (**), 0% vs 40%: p = 0.046 (*),**<br>0% vs 60%: p = 0.110 (ns), **0% vs 80%: p = 0.0003 (***),**<br>**5% vs 10%: p = 0.036 (*), 5% vs 20%: p = 0.0002 (***),**<br>**5% vs 40%: p = 0.021 (*),** 5% vs 60%: p = 0.116 (ns),<br>**5% vs 80%: p < 0.0001 (****),** 10% vs 20%: p = 0.229 (ns),<br>10% vs 40%: p = 0.882 (ns), 10% vs 60%: p = 0.476 (ns),<br>**10% vs 80%: p = 0.023 (*)**, 20% vs 40%: p = 0.089 (ns),<br>**20% vs 60%: p = 0.027 (*)**, 20% vs 80%: p = 0.194 (ns),<br>40% vs 60%: p = 0.473 (ns), **40% vs 80%: p = 0.003 (**)**,<br>**60% vs 80%: p = 0.001 (**)** | Hierarchical Bootstrap:<br>0% vs 5%: p = 0.604 (ns), 0% vs 10%: p = 0.165 (ns),<br>0% vs 20%: p = 0.107 (ns), 0% vs 40%: p = 0.207 (ns),<br>0% vs 60%: p = 0.292 (ns), **0% vs 80%: p = 0.043 (*),**<br>5% vs 10%: p = 0.069 (ns), **5% vs 20%: p = 0.031 (*),**<br>5% vs 40%: p = 0.146 (ns), 5% vs 60%: p = 0.306 (ns),<br>**5% vs 80%: p = 0.004 (**),** 10% vs 20%: p = 0.212 (ns),<br>10% vs 40%: p = 0.437 (ns), 10% vs 60%: p = 0.312 (ns),<br>10% vs 80%: p = 0.107 (ns), 20% vs 40%: p = 0.122 (ns),<br>20% vs 60%: p = 0.126 (ns), 20% vs 80%: p = 0.251 (ns),<br>40% vs 60%: p = 0.295 (ns), **40% vs 80%: p = 0.029 (*),**<br>**60% vs 80%: p = 0.013 (*)** |
| Figure 5f:<br>LAMP5$^{L1}$ contrast | 0%: 0.048 ± 0.008<br>5%: 0.024 ± 0.009<br>10%: 0.036 ± 0.012<br>20%: 0.033 ± 0.009<br>40%: 0.005 ± 0.005<br>60%: -0.003 ± 0.004<br>80%: -0.013 ± 0.007<br><br>*LAMP5$^{L1}$: N = 8, n = 251 neurons* | Mixed Linear Model Regression:<br>**0% vs 5%: p = 0.005 (**), 0% vs 10%: p = 0.040 (*),**<br>**0% vs 20%: p = 0.028 (*), 0% vs 40%: p < 0.0001 (****),**<br>**0% vs 60%: p < 0.0001 (****), 0% vs 80%: p < 0.0001 (****),**<br>5% vs 10%: p = 0.209 (ns), 5% vs 20%: p = 0.341 (ns),<br>**5% vs 40%: p < 0.0001 (****), 5% vs 60%: p < 0.0001 (****),**<br>**5% vs 80%: p < 0.0001 (****),** 10% vs 20%: p = 0.788 (ns),<br>**10% vs 40%: p < 0.0001 (****), 10% vs 60%: p < 0.0001 (****),**<br>**10% vs 80%: p < 0.0001 (****), 20% vs 40%: p < 0.0001 (****),**<br>**20% vs 60%: p < 0.0001 (****), 20% vs 80%: p < 0.0001 (****),**<br>**40% vs 60%: p = 0.027 (*), 40% vs 80%: p < 0.0001 (****),**<br>**60% vs 80%: p = 0.002 (**)** | Hierarchical Bootstrap:<br>**0% vs 5%: p = 0.027 (*),** 0% vs 10%: p = 0.114 (ns),<br>0% vs 20%: p = 0.115 (ns), **0% vs 40%: p = 0.001 (**),**<br>**0% vs 60%: p < 0.0001 (****), 0% vs 80%: p < 0.0001 (****),**<br>5% vs 10%: p = 0.186 (ns), 5% vs 20%: p = 0.233 (ns),<br>**5% vs 40%: p = 0.037 (*), 5% vs 60%: p = 0.005 (**),**<br>**5% vs 80%: p = 0.0005 (***),** 10% vs 20%: p = 0.487 (ns),<br>**10% vs 40%: p = 0.027 (*), 10% vs 60%: p = 0.002 (**),**<br>**10% vs 80%: p = 0.0007 (***), 20% vs 40%: p = 0.012 (*),**<br>**20% vs 60%: p = 0.0007 (***), 20% vs 80%: p = 0.0004 (***),**<br>40% vs 60%: p = 0.098 (ns), **40% vs 80%: p = 0.0009 (***),**<br>**60% vs 80%: p = 0.048 (*)** |
| Figure 5h:<br>LAMP5$^{L23}$ contrast (run) | 0%: -0.010 ± 0.054<br>5%: 0.013 ± 0.037<br>10%: 0.028 ± 0.043<br>20%: 0.054 ± 0.021<br>40%: 0.083 ± 0.025<br>60%: 0.065 ± 0.021<br>80%: 0.115 ± 0.027 | Mixed Linear Model Regression:<br>0% vs 5%: p = 0.312 (ns), 0% vs 10%: p = 0.360 (ns),<br>0% vs 20%: p = 0.073 (ns), **0% vs 40%: p = 0.022 (*),**<br>**0% vs 60%: p = 0.019 (*), 0% vs 80%: p = 0.0005 (***),**<br>5% vs 10%: p = 0.786 (ns), 5% vs 20%: p = 0.332 (ns),<br>5% vs 40%: p = 0.113 (ns), 5% vs 60%: p = 0.090 (ns),<br>**5% vs 80%: p = 0.001 (**),** 10% vs 20%: p = 0.128 (ns),<br>**10% vs 40%: p = 0.018 (*), 10% vs 60%: p = 0.015 (*),**<br>**10% vs 80%: p < 0.0001 (****),** 20% vs 40%: p = 0.409 (ns),<br>20% vs 60%: p = 0.358 (ns), **20% vs 80%: p = 0.002 (**),**<br>40% vs 60%: p = 0.920 (ns), **40% vs 80%: p = 0.042 (*),**<br>60% vs 80%: p = 0.055 (ns) | Hierarchical Bootstrap:<br>0% vs 5%: p = 0.314 (ns), 0% vs 10%: p = 0.306 (ns),<br>0% vs 20%: p = 0.158 (ns), 0% vs 40%: p = 0.112 (ns),<br>0% vs 60%: p = 0.119 (ns), 0% vs 80%: p = 0.058 (ns),<br>5% vs 10%: p = 0.485 (ns), 5% vs 20%: p = 0.763 (ns),<br>5% vs 40%: p = 0.779 (ns), 5% vs 60%: p = 0.814 (ns),<br>5% vs 80%: p = 0.050 (ns), 10% vs 20%: p = 0.262 (ns),<br>10% vs 40%: p = 0.097 (ns), 10% vs 60%: p = 0.152 (ns),<br>10% vs 80%: p = 0.059 (ns), 20% vs 40%: p = 0.347 (ns),<br>20% vs 60%: p = 0.393 (ns), 20% vs 80%: p = 0.060 (ns),<br>40% vs 60%: p = 0.554 (ns), 40% vs 80%: p = 0.174 (ns),<br>60% vs 80%: p = 0.141 (ns) |

| | | | |
|---|---|---|---|
| Figure 5h: LAMP5$^{L23}$ contrast (stationary) | 0%: -0.023 ± 0.022<br>5%: -0.028 ± 0.007<br>10%: -0.015 ± 0.004<br>20%: -0.008 ± 0.007<br>40%: -0.026 ± 0.007<br>60%: -0.024 ± 0.010<br>80%: -0.015 ± 0.007 | Mixed Linear Model Regression:<br>0% vs 5%: p = 0.812 (ns), 0% vs 10%: p = 0.224 (ns),<br>0% vs 20%: p = 0.075 (ns), 0% vs 40%: p = 0.840 (ns),<br>0% vs 60%: p = 0.736 (ns), 0% vs 80%: p = 0.488 (ns),<br>**5% vs 10%: p = 0.033 (*), 5% vs 20%: p = 0.001 (**),**<br>5% vs 40%: p = 0.961 (ns), 5% vs 60%: p = 0.381 (ns),<br>5% vs 80%: p = 0.161 (ns), 10% vs 20%: p = 0.525 (ns),<br>**10% vs 40%: p = 0.033 (*),** 10% vs 60%: p = 0.199 (ns),<br>10% vs 80%: p = 0.443 (ns), **20% vs 40%: p = 0.0003 (***),**<br>**20% vs 60%: p = 0.020 (*),** 20% vs 80%: p = 0.101 (ns),<br>40% vs 60%: p = 0.382 (ns), 40% vs 80%: p = 0.164 (ns),<br>60% vs 80%: p = 0.589 (ns) | Hierarchical Bootstrap:<br>0% vs 5%: p = 0.470 (ns), 0% vs 10%: p = 0.284 (ns),<br>0% vs 20%: p = 0.221 (ns), 0% vs 40%: p = 0.490 (ns),<br>0% vs 60%: p = 0.465 (ns), 0% vs 80%: p = 0.338 (ns),<br>5% vs 10%: p = 0.094 (ns), 5% vs 20%: p = 0.058 (ns),<br>5% vs 40%: p = 0.461 (ns), 5% vs 60%: p = 0.397 (ns),<br>5% vs 80%: p = 0.187 (ns), 10% vs 20%: p = 0.308 (ns),<br>10% vs 40%: p = 0.102 (ns), 10% vs 60%: p = 0.221 (ns),<br>10% vs 80%: p = 0.341 (ns), **20% vs 40%: p = 0.019 (*),**<br>20% vs 60%: p = 0.124 (ns), 20% vs 80%: p = 0.175 (ns),<br>40% vs 60%: p = 0.418 (ns), 40% vs 80%: p = 0.197 (ns),<br>60% vs 80%: p = 0.304 (ns) |
| Figure 5j: LAMP5$^{L1}$ contrast (run) | 0%: 0.154 ± 0.071<br>5%: 0.064 ± 0.019<br>10%: 0.116 ± 0.045<br>20%: 0.096 ± 0.024<br>40%: 0.088 ± 0.074<br>60%: 0.018 ± 0.016<br>80%: 0.043 ± 0.010 | Mixed Linear Model Regression:<br>**0% vs 5%: p < 0.0001 (****), 0% vs 10%: p = 0.005 (**),**<br>**0% vs 20%: p = 0.005 (**), 0% vs 40%: p < 0.0001 (****),**<br>**0% vs 60%: p < 0.0001 (****), 0% vs 80%: p < 0.0001 (****),**<br>**5% vs 10%: p = 0.001 (**), 5% vs 20%: p = 0.021 (*),**<br>5% vs 40%: p = 0.579 (ns), 5% vs 60%: p = 0.065 (ns),<br>5% vs 80%: p = 0.313 (ns), 10% vs 20%: p = 0.642 (ns),<br>**10% vs 40%: p < 0.0001 (****), 10% vs 60%: p < 0.0001 (****),**<br>**10% vs 80%: p < 0.0001 (****), 20% vs 40%: p = 0.007 (**),**<br>**20% vs 60%: p < 0.0001 (****), 20% vs 80%: p = 0.003 (**),**<br>40% vs 60%: p = 0.209 (ns), 40% vs 80%: p = 0.614 (ns),<br>60% vs 80%: p = 0.535 (ns) | Hierarchical Bootstrap:<br>**0% vs 5%: p = 0.042 (*),** 0% vs 10%: p = 0.106 (ns),<br>0% vs 20%: p = 0.087 (ns), 0% vs 40%: p = 0.064 (ns),<br>**0% vs 60%: p = 0.008 (**), 0% vs 80%: p = 0.018 (*),**<br>5% vs 10%: p = 0.105 (ns), 5% vs 20%: p = 0.088 (ns),<br>5% vs 40%: p = 0.378 (ns), 5% vs 60%: p = 0.069 (ns),<br>5% vs 80%: p = 0.253 (ns), 10% vs 20%: p = 0.423 (ns),<br>10% vs 40%: p = 0.158 (ns), **10% vs 60%: p = 0.018 (*),**<br>**10% vs 80%: p = 0.040 (*),** 20% vs 40%: p = 0.234 (ns),<br>**20% vs 60%: p = 0.004 (**), 20% vs 80%: p = 0.036 (*),**<br>40% vs 60%: p = 0.362 (ns), 40% vs 80%: p = 0.461 (ns),<br>60% vs 80%: p = 0.272 (ns) |
| Figure 5j: LAMP5$^{L1}$ contrast (stationary) | 0%: 0.036 ± 0.010<br>5%: 0.013 ± 0.010<br>10%: 0.021 ± 0.009<br>20%: 0.023 ± 0.010<br>40%: -0.010 ± 0.007<br>60%: -0.007 ± 0.005<br>80%: -0.026 ± 0.008 | Mixed Linear Model Regression:<br>0% vs 5%: p = 0.068 (ns), 0% vs 10%: p = 0.056 (ns),<br>0% vs 20%: p = 0.069 (ns), **0% vs 40%: p < 0.0001 (****),**<br>**0% vs 60%: p < 0.0001 (****), 0% vs 80%: p < 0.0001 (****),**<br>5% vs 10%: p = 0.942 (ns), 5% vs 20%: p = 0.945 (ns),<br>**5% vs 40%: p < 0.0001 (****), 5% vs 60%: p < 0.0001 (****),**<br>**5% vs 80%: p < 0.0001 (****),** 10% vs 20%: p = 0.883 (ns),<br>**10% vs 40%: p < 0.0001 (****), 10% vs 60%: p < 0.0001 (****),**<br>**10% vs 80%: p < 0.0001 (****), 20% vs 40%: p < 0.0001 (****),**<br>**20% vs 60%: p < 0.0001 (****), 20% vs 80%: p < 0.0001 (****),**<br>40% vs 60%: p = 0.905 (ns), **40% vs 80%: p < 0.0001 (****),**<br>**60% vs 80%: p < 0.0001 (****)** | Hierarchical Bootstrap:<br>0% vs 5%: p = 0.079(ns), 0% vs 10%: p = 0.111 (ns),<br>0% vs 20%: p = 0.142 (ns), **0% vs 40%: p = 0.005 (**),**<br>**0% vs 60%: p = 0.003 (**), 0% vs 80%: p < 0.0001 (****),**<br>5% vs 10%: p = 0.375 (ns), 5% vs 20%: p = 0.325 (ns),<br>**5% vs 40%: p = 0.043 (*), 5% vs 60%: p = 0.042 (*),**<br>**5% vs 80%: p = 0.0004 (***),** 10% vs 20%: p = 0.409 (ns),<br>**10% vs 40%: p = 0.031 (*), 10% vs 60%: p = 0.014 (*),**<br>**10% vs 80%: p = 0.0002 (***), 20% vs 40%: p = 0.015 (*),**<br>**20% vs 60%: p = 0.008 (**), 20% vs 80%: p < 0.0001 (****),**<br>40% vs 60%: p = 0.441 (ns), **40% vs 80%: p = 0.028 (*),**<br>**60% vs 80%: p = 0.016 (*)** |

**Table 5.1 - related to Extended Data 8 - Figure 5**

| | Mean ± SEM | P value |
|---|---|---|
| Ex 8c:<br>gOSI | $LAMP5^{L23}$: 0.244 ± 0.014<br>$LAMP5^{L1}$: 0.210 ± 0.012<br><br>*$LAMP5^{L23}$: N = 8, n = 132 neurons*<br>*$LAMP5^{L1}$: N = 8, n = 98 neurons* | Hierarchical bootstrap: p = 0.033 (*)<br>Mixed Linear Model Regression: p = 0.018 (*) |
| Ex 8d:<br>Visual response (run) | Orientation 0°<br>$LAMP5^{L23}$: 0.345 ± 0.066<br>$LAMP5^{L1}$: 0.093 ± 0.068<br>Orientation 30°<br>$LAMP5^{L23}$: 0.283 ± 0.054<br>$LAMP5^{L1}$: 0.166 ± 0.060<br>Orientation 60°<br>$LAMP5^{L23}$: 0.446 ± 0.074<br>$LAMP5^{L1}$: 0.322 ± 0.085<br>Orientation 90°<br>$LAMP5^{L23}$: 0.845 ± 0.104<br>$LAMP5^{L1}$: 0.609 ± 0.130<br>Orientation 120°<br>$LAMP5^{L23}$: 0.376 ± 0.073<br>$LAMP5^{L1}$: 0.310 ± 0.106<br>Orientation 150°<br>$LAMP5^{L23}$: 0.378 ± 0.072<br>$LAMP5^{L1}$: 0.248 ± 0.090 | Mixed Linear Model Regression:<br>0° $LAMP5^{L23}$ vs $LAMP5^{L1}$: p = 0.158 (ns)<br>30° $LAMP5^{L23}$ vs $LAMP5^{L1}$: p = 0.031 (*)<br>60° $LAMP5^{L23}$ vs $LAMP5^{L1}$: p = 0.093 (ns)<br>90° $LAMP5^{L23}$ vs $LAMP5^{L1}$: p = 0.023 (*)<br>120° $LAMP5^{L23}$ vs $LAMP5^{L1}$: p = 0.362 (ns)<br>150° $LAMP5^{L23}$ vs $LAMP5^{L1}$: p = 0.070 (ns) |
| Ex 8d:<br>Visual response (stationary) | Orientation 0°<br>$LAMP5^{L23}$: 0.078 ± 0.021<br>$LAMP5^{L1}$: 0.040 ± 0.031<br>Orientation 30°<br>$LAMP5^{L23}$: 0.055 ± 0.020<br>$LAMP5^{L1}$: 0.073 ± 0.029<br>Orientation 60°<br>$LAMP5^{L23}$: 0.106 ± 0.022<br>$LAMP5^{L1}$: 0.085 ± 0.035<br>Orientation 90°<br>$LAMP5^{L23}$: 0.252 ± 0.028<br>$LAMP5^{L1}$: 0.219 ± 0.049<br>Orientation 120°<br>$LAMP5^{L23}$: 0.100 ± 0.023<br>$LAMP5^{L1}$: 0.084 ± 0.031<br>Orientation 150°<br>$LAMP5^{L23}$: 0.077 ± 0.020<br>$LAMP5^{L1}$: 0.070 ± 0.028 | Mixed Linear Model Regression:<br>0° $LAMP5^{L23}$ vs $LAMP5^{L1}$: p = 0.078 (ns)<br>30° $LAMP5^{L23}$ vs $LAMP5^{L1}$: p = 0.348 (ns)<br>60° $LAMP5^{L23}$ vs $LAMP5^{L1}$: p = 0.335 (ns)<br>90° $LAMP5^{L23}$ vs $LAMP5^{L1}$: p = 0.242 (ns)<br>120° $LAMP5^{L23}$ vs $LAMP5^{L1}$: p = 0.497 (ns)<br>150° $LAMP5^{L23}$ vs $LAMP5^{L1}$: p = 0.321 (ns) |

**Table 5.2 - related to Extended Data 9 - Figure 5**

| | Mean ± SEM | P value | |
|---|---|---|---|
| Ex 9e:<br>Ratio of contrast preference (run) | Ratio preferring high contrasts vs low contrasts:<br>LAMP5$^{L23}$: 0.619 vs 0.381 ± 0.052<br>LAMP5$^{L1}$: 0.367 vs 0.632 ± 0.057<br><br>*LAMP5$^{L23}$: N = 8, n = 180 neurons*<br>*LAMP5$^{L1}$: N = 8, n = 251 neurons* | Mann Whitney test:<br>LAMP$^{L23}$: p = 0.0104 (*)<br>LAMP5$^{L}$: p = 0.0044 (**) | |
| Ex 9e:<br>Ratio of contrast preference (stationary ) | Ratio preferring high contrasts vs low contrasts:<br>LAMP5$^{L23}$: 0.434 vs 0.566 ± 0.053<br>LAMP5$^{L1}$: 0.265 vs 0.735 ± 0.077<br><br>*LAMP5$^{L23}$: N = 8, n = 180 neurons*<br>*LAMP5$^{L1}$: N = 8, n = 251 neurons* | Mann Whitney test:<br>LAMP$^{L23}$: p = 0.241 (ns)<br>LAMP5$^{L}$: p = 0.0025 (**) | |
| Ex 9f:<br>JAM2$^{L23}$ contrast | 0%: -0.022 ± 0.022<br>5%: -0.006 ± 0.010<br>10%: 0.007 ± 0.013<br>20%: 0.013 ± 0.010<br>40%: -0.014 ± 0.005<br>60%: -0.000 ± 0.013<br>80%: 0.001 ± 0.011<br><br>*JAM$^{L23}$: N = 9, n = 120 neurons* | Mixed Linear Model Regression:<br>0% vs 5%: p = 0.479 (ns), 0% vs 10%: p = 0.128 (ns),<br>0% vs 20%: p = 0.109 (ns), 0% vs 40%: p = 0.894 (ns),<br>0% vs 60%: p = 0.571 (ns), 0% vs 80%: p = 0.421 (ns),<br>5% vs 10%: p = 0.195 (ns), 5% vs 20%: p = 0.121 (ns),<br>5% vs 40%: p = 0.289 (ns), 5% vs 60%: p = 0.766 (ns),<br>5% vs 80%: p = 0.895 (ns), 10% vs 20%: p = 0.970 (ns),<br>**10% vs 40%: p = 0.014 (*)**, 10% vs 60%: p = 0.082 (ns),<br>10% vs 80%: p = 0.206 (ns), **20% vs 40%: p = 0.002 (**),**<br>**20% vs 60%: p = 0.031 (*),** 20% vs 80%: p = 0.113 (ns),<br>40% vs 60%: p = 0.370 (ns), 40% vs 80%: p = 0.178 (ns),<br>60% vs 80%: p = 0.618 (ns) | Hierarchical Bootstrap:<br>0% vs 5%: p = 0.304 (ns), 0% vs 10%: p = 0.130 (ns),<br>0% vs 20%: p = 0.147 (ns), 0% vs 40%: p = 0.418 (ns),<br>0% vs 60%: p = 0.318 (ns) 0% vs 80%: p = 0.274 (ns),<br>5% vs 10%: p = 0.176 (ns), 5% vs 20%: p = 0.159 (ns),<br>5% vs 40%: p = 0.244 (ns), 5% vs 60%: p = 0.525 (ns),<br>5% vs 80%: p = 0.451 (ns), 10% vs 20%: p = 0.495 (ns), 10% vs 40%: p = 0.064 (ns), 10% vs 60%: p = 0.183 (ns), 10% vs 80%: p = 0.234 (ns),<br>**20% vs 40%: p = 0.029 (*),** 20% vs 60%: p = 0.146 (ns), 20% vs 80%: p = 0.166 (ns),<br>40% vs 60%: p = 0.278 (ns), 40% vs 80%: p = 0.196 (ns),<br>60% vs 80%: p = 0.417 (ns) |
| Ex 9h:<br>JAM2$^{L23}$ contrast (run) | 0%: -0.084 ± 0.138<br>5%: 0.003 ± 0.022<br>10%: 0.005 ± 0.032<br>20%: 0.051 ± 0.039<br>40%: 0.039 ± 0.022<br>60%: 0.121 ± 0.053<br>80%: 0.110 ± 0.033 | Mixed Linear Model Regression:<br>0% vs 5%: p = 0.343 (ns), 0% vs 10%: p = 0.184 (ns),<br>0% vs 20%: p = 0.107 (ns), **0% vs 40%: p = 0.048 (*)**,<br>**0% vs 60%: p = 0.011 (*)**, **0% vs 80%: p = 0.002 (**),**<br>5% vs 10%: p = 0.525 (ns), 5% vs 20%: p = 0.192 (ns),<br>5% vs 40%: p = 0.078 (ns), **5% vs 60%: p = 0.005 (**),**<br>**5% vs 80%: p < 0.0001 (****),** 10% vs 20%: p = 0.522 (ns),<br>10% vs 40%: p = 0.232 (ns), **10% vs 60%: p = 0.022 (*),**<br>**10% vs 80%: p = 0.0006 (***),** 20% vs 40%: p = 0.498 (ns),<br>20% vs 60%: p = 0.063 (ns), **20% vs 80%: p = 0.003 (**),**<br>40% vs 60%: p = 0.317 (ns), **40% vs 80%: p = 0.038 (*),**<br>60% vs 80%: p = 0.224 (ns) | Hierarchical Bootstrap:<br>0% vs 5%: p = 0.380 (ns), 0% vs 10%: p = 0.301 (ns),<br>0% vs 20%: p = 0.287 (ns), 0% vs 40%: p = 0.18 (ns),<br>0% vs 60%: p = 0.109 (ns), 0% vs 80%: p = 0.054 (ns),<br>5% vs 10%: p = 0.377 (ns), 5% vs 20%: p = 0.267 (ns),<br>5% vs 40%: p = 0.170 (ns), 5% vs 60%: p = 0.104 (ns),<br>5% vs 80%: p = 0.016 (*), 10% vs 20%: p = 0.368 (ns),<br>10% vs 40%: p = 0.262 (ns), 10% vs 60%: p = 0.143 (ns), **10% vs 80%: p = 0.029 (*)**, 20% vs 40%: p = 0.409 (ns), 20% vs 60%: p = 0.217 (ns), 20% vs 80%: p = 0.091 (ns),<br>40% vs 60%: p = 0.264 (ns), 40% vs 80%: p = 0.089 (ns),<br>60% vs 80%: p = 0.247 (ns) |
| Ex 9h:<br>JAM2$^{L23}$ contrast (stationary ) | 0%: -0.031 ± 0.020<br>5%: 0.002 ± 0.010<br>10%: 0.005 ± 0.007<br>20%: -0.001 ± 0.006<br>40%: -0.024 ± 0.007<br>60%: -0.037 ± 0.011<br>80%: -0.024 ± 0.008 | Mixed Linear Model Regression:<br>0% vs 5%: p = 0.107 (ns), 0% vs 10%: p = 0.077 (ns),<br>0% vs 20%: p = 0.201 (ns), 0% vs 40%: p = 0.680 (ns),<br>0% vs 60%: p = 0.483 (ns), 0% vs 80%: p = 0.722 (ns),<br>5% vs 10%: p = 0.791 (ns), 5% vs 20%: p = 0.565 (ns),<br>**5% vs 40%: p = 0.0005 (***), 5% vs 60%: p < 0.0001 (****),**<br>**5% vs 80%: p = 0.001 (**),** 10% vs 20%: p = 0.378 (ns),<br>**10% vs 40%: p < 0.0001 (****), 10% vs 60%: p < 0.0001 (****),**<br>**10% vs 80%: p = 0.0003 (***), 20% vs 40%: p = 0.002 (**),**<br>**20% vs 60%: p = 0.0002 (***), 20% vs 80%: p = 0.004 (**),**<br>40% vs 60%: p = 0.590 (ns), 40% vs 80%: p = 0.926 (ns),<br>60% vs 80%: p = 0.553 (ns) | Hierarchical Bootstrap:<br>0% vs 5%: p = 0.122 (ns), 0% vs 10%: p = 0.108 (ns),<br>0% vs 20%: p = 0.179 (ns), 0% vs 40%: p = 0.462 (ns),<br>0% vs 60%: p = 0.397 (ns), 0% vs 80%: p = 0.474 (ns)<br>5% vs 10%: p = 0.377 (ns), 5% vs 20%: p = 0.608 (ns),<br>**5% vs 40%: p = 0.025 (*), 5% vs 60%: p = 0.025 (*),**<br>**5% vs 80%: p = 0.026 (*),** 10% vs 20%: p = 0.271 (ns), **10% vs 40%: p = 0.003 (**), 10% vs 60%: p = 0.006 (**), 10% vs 80%: p = 0.011 (*), 20% vs 40%: p = 0.020 (*), 20% vs 60%: p = 0.027 (*), 20% vs 80%: p = 0.024 (*),**<br>40% vs 60%: p = 0.360 (ns), 40% vs 80%: p = 0.506 (ns),<br>60% vs 80%: p = 0.627 (ns) |
| Ex 9j:<br>JAM2$^{L1}$ contrast | 0%: 0.042 ± 0.024<br>5%: 0.016 ± 0.009<br>10%: 0.011 ± 0.008<br>20%: 0.018 ± 0.012<br>40%: -0.006 ± 0.007<br>60%: -0.018 ± 0.010<br>80%: -0.023 ± 0.014<br><br>*JAM2$^{L1}$: N = 11, n = 144 neurons* | Mixed Linear Model Regression:<br>0% vs 5%: p = 0.142 (ns), **0% vs 10%: p = 0.024 (*),**<br>0% vs 20%: p = 0.067 (ns), **0% vs 40%: p = 0.0002 (***),**<br>**0% vs 60%: p < 0.0001 (****), 0% vs 80%: p < 0.0001 (****),**<br>5% vs 10%: p = 0.103 (ns), 5% vs 20%: p = 0.435 (ns),<br>**5% vs 40%: p < 0.0001 (****), 5% vs 60%: p < 0.0001** | Hierarchical Bootstrap:<br>0% vs 5%: p = 0.152 (ns), 0% vs 10%: p = 0.081 (ns),<br>0% vs 20%: p = 0.143 (ns), **0% vs 40%: p = 0.013 (*),**<br>**0% vs 60%: p = 0.007 (**), 0% vs 80%: p = 0.007 (**),**<br>5% vs 10%: p = 0.206 (ns), 5% vs 20%: p = 0.393 (ns), |

| | | | |
|---|---|---|---|
| | | **(****),**<br>**5% vs 80%: p < 0.0001 (****),** 10% vs 20%: p = 0.437 (ns), **10% vs 40%: p = 0.002 (**), 10% vs 60%: p = 0.0001 (***), 10% vs 80%: p < 0.0001 (****), 20% vs 40%: p = 0.0002 (***), 20% vs 60%: p < 0.0001 (****), 20% vs 80%: p < 0.0001 (****),**<br>40% vs 60%: p = 0.282 (ns), **40% vs 80%: p = 0.004 (**),**<br>60% vs 80%: p = 0.088 (ns) | **5% vs 40%: p = 0.009 (**), 5% vs 60%: p = 0.005 (**),**<br>**5% vs 80%: p = 0.002 (**),** 10% vs 20%: p = 0.366 (ns), **10% vs 40%: p = 0.036 (*), 10% vs 60%: p = 0.017 (*), 10% vs 80%: p = 0.008 (**), 20% vs 40%: p = 0.026 (*), 20% vs 60%: p = 0.011 (*), 20% vs 80%: p = 0.006 (**),**<br>40% vs 60%: p = 0.231 (ns), 40% vs 80%: p = 0.107 (ns),<br>60% vs 80%: p = 0.263 (ns) |
| Ex 9l: JAM2$^{L1}$ contrast (run) | 0%: 0.130 ± 0.075<br>5%: 0.036 ± 0.038<br>10%: 0.066 ± 0.021<br>20%: 0.050 ± 0.027<br>40%: 0.048 ± 0.041<br>60%: 0.048 ± 0.033<br>80%: 0.022 ± 0.031 | Mixed Linear Model Regression:<br>0% vs 5%: p = 0.160 (ns), 0% vs 10%: p = 0.319 (ns), 0% vs 20%: p = 0.262 (ns), 0% vs 40%: p = 0.060 (ns),<br>0% vs 60%: p = 0.073 (ns), **0% vs 80%: p = 0.006 (*),**<br>5% vs 10%: p = 0.566 (ns), 5% vs 20%: p = 0.518 (ns),<br>5% vs 40%: p = 0.562 (ns), 5% vs 60%: p = 0.638 (ns),<br>**5% vs 80%: p = 0.035 (*),** 10% vs 20%: p = 0.950 (ns),<br>10% vs 40%: p = 0.243 (ns), 10% vs 60%: p = 0.291 (ns),<br>**10% vs 80%: p = 0.011 (*),** 20% vs 40%: p = 0.208 (ns),<br>20% vs 60%: p = 0.239 (ns)**, 20% vs 80%: p = 0.003 (**),**<br>40% vs 60%: p = 0.867 (ns), 40% vs 80%: p = 0.165 (ns),<br>60% vs 80%: p = 0.092 (ns) | Hierarchical Bootstrap:<br>0% vs 5%: p = 0.260 (ns), 0% vs 10%: p = 0.335 (ns),<br>0% vs 20%: p = 0.249 (ns), 0% vs 40%: p = 0.151 (ns),<br>0% vs 60%: p = 0.190 (ns), 0% vs 80%: p = 0.108 (ns),<br>5% vs 10%: p = 0.331 (ns), 5% vs 20%: p = 0.410 (ns),<br>5% vs 40%: p = 0.458 (ns), 5% vs 60%: p = 0.480 (ns),<br>5% vs 80%: p = 0.180 (ns), 10% vs 20%: p = 0.368 (ns), 10% vs 40%: p = 0.216 (ns), 10% vs 60%: p = 0.255 (ns), 10% vs 80%: p = 0.077 (ns), 20% vs 40%: p = 0.381 (ns), 20% vs 60%: p = 0.388 (ns), 20% vs 80%: p = 0.130 (ns),<br>40% vs 60%: p = 0.467 (ns), 40% vs 80%: p = 0.262 (ns),<br>60% vs 80%: p = 0.181 (ns) |
| Ex 9l: JAM2$^{L1}$ contrast (stationary) | 0%: 0.025 ± 0.019<br>5%: -0.003 ± 0.012<br>10%: -0.002 ± 0.010<br>20%: 0.017 ± 0.017<br>40%: -0.017 ± 0.012<br>60%: -0.023 ± 0.010<br>80%: -0.037 ± 0.013 | Mixed Linear Model Regression:<br>0% vs 5%: p = 0.083 (ns), **0% vs 10%: p = 0.045 (*),**<br>0% vs 20%: p = 0.402 (ns), **0% vs 40%: p = 0.002 (**),**<br>**0% vs 60%: p = 0.0007 (***), 0% vs 80%: p < 0.0001 (****),**<br>5% vs 10%: p = 0.579 (ns), 5% vs 20%: p = 0.198 (ns),<br>**5% vs 40%: p = 0.006 (**), 5% vs 60%: p = 0.002 (**), 5% vs 80%: p < 0.0001 (****),** 10% vs 20%: p = 0.070 (ns), **10% vs 40%: p = 0.028 (*), 10% vs 60%: p = 0.009 (**),**<br>**10% vs 80%: p < 0.0001 (****), 20% vs 40%: p = 0.0003 (***), 20% vs 60%: p < 0.0001 (****), 20% vs 80%: p < 0.0001 (****),** 40% vs 60%: p = 0.661 (ns),<br>**40% vs 80%: p = 0.005 (**),**<br>**60% vs 80%: p = 0.021 (*)** | Hierarchical Bootstrap:<br>0% vs 5%: p = 0.100 (ns), 0% vs 10%: p = 0.091 (ns),<br>0% vs 20%: p = 0.335 (ns), **0% vs 40%: p = 0.024 (*),**<br>**0% vs 60%: p = 0.014 (*), 0% vs 80%: p = 0.005 (**),**<br>5% vs 10%: p = 0.394 (ns), 5% vs 20%: p = 0.268 (ns),<br>5% vs 40%: p = 0.060 (ns), **5% vs 60%: p = 0.044 (*),**<br>**5% vs 80%: p = 0.005 (**),** 10% vs 20%: p = 0.200 (ns), 10% vs 40%: p = 0.089 (ns), 10% vs 60%: p = 0.053 (ns), **10% vs 80%: p = 0.004 (**), 20% vs 40%: p = 0.046 (*), 20% vs 60%: p = 0.023 (*), 20% vs 80%: p = 0.002 (**),** 40% vs 60%: p = 0.361 (ns), 40% vs 80%: p = 0.053 (ns)<br>60% vs 80%: p = 0.146 (ns) |
| Ex 9p: Ratio of contrast preference | Ratio preferring high contrasts vs low contrasts:<br>JAM2$^{L23}$: 0.516 vs 0.484 ± 0.051<br>JAM2$^{L1}$: 0.283 vs 0.717 ± 0.070<br>VIP$^{L23}$: 0.140 vs 0.860 ± 0.083<br>SST$^{L23}$: 0.917 vs 0.083 ± 0.053<br><br>*JAM$^{L23}$: N = 9, n = 120 neurons*<br>*JAM2$^{L1}$: N = 11, n = 144 neurons*<br>*VIP$^{L23}$: N = 4, n = 83 neurons*<br>*SST$^{L23}$: N = 6, n = 56 neurons* | Mann Whitney test:<br>JAM2$^{L23}$: p = 0.713 (ns)<br>JAM2$^{L1}$: p = 0.0003 (***)<br>VIP$^{L23}$: p = 0.0286 (*)<br>SST$^{L23}$: p = 0.0022 (**) | |

**Table 6.1 - related to Extended Data 10 - Figure 6**

| | Mean ± SEM | P value | |
|---|---|---|---|
| Ex 10b: SST->JAM2(LAMP5+ cINs) | L1: 306 ± 67.55<br>L23: 212.8 ± 37.42<br>L4: 318.6 ± 52.75<br>L5: 313.8 ± 48.78<br>L6: 212.8 ± 28.21<br>PYN$^{L23}$: 861.7 ± 97.8<br><br>N = 7, LAMP5$^{L1}$ = 12; LAMP5$^{L23}$ = 12; LAMP5$^{L4}$ = 9; LAMP5$^{L5}$ = 12; LAMP5$^{L6}$ = 8; PYN$^{L23}$ = 13 cells. | Kruskal-Wallis test: p < 0.0001 | Uncorrected Dunn's test:<br>L1 vs. L23: p = 0.2174 (ns)<br>L1 vs. L4: p = 0.6962 (ns)<br>L1 vs. L5: p = 0.6979 (ns)<br>L1 vs. L6: p = 0.5333 (ns)<br>L1 vs. PYN$^{L23}$: p < 0.0001 (****)<br>L23 vs. L4: p = 0.1254 (ns)<br>L23 vs. L5: p = 0.1049 (ns)<br>L23 vs. L6: p = 0.6310 (ns)<br>L23 vs. PYN$^{L23}$: p < 0.0001 (****)<br>L4 vs. L5: p = 0.9751 (ns)<br>L4 vs. L6: p = 0.3474 (ns)<br>L4 vs. PYN$^{L23}$: p = 0.0016 (**)<br>L5 vs. L6: p = 0.3320 (ns)<br>L5 vs. PYN$^{L23}$: p = 0.0005 (***)<br><br>L6 vs. PYN$^{L23}$: p < 0.0001 (****) |
| Ex 10c: PV->JAM2(LAMP5+ cINs) | L1: 5.527 ± 2.882<br>L23: 385.1 ± 68.96<br>L4: 555.4 ± 52.33 | Kruskal-Wallis test: p < 0.0001 | Uncorrected Dunn's test:<br>L1 vs. L23: p < 0.0001 (****)<br>L1 vs. L4: p < 0.0001 (****) |

| | | | |
|---|---|---|---|
| | L5: 222.6 ± 35.37<br>L6: 182.5 ± 21.58<br>$PYN^{L23}$: 786.4 ± 109.1<br><br>N = 4, $LAMP5^{L1}$ = 11; $LAMP5^{L23}$ = 11; $LAMP5^{L4}$ = 12; $LAMP5^{L5}$ = 11; $LAMP5^{L6}$ = 11; $PYN^{L23}$ = 11 cells. | | L1 vs. L5: p = 0.0092 (**)<br>L1 vs. L6: p = 0.0396 (*)<br>L1 vs. $PYN^{L23}$: p < 0.0001 (****)<br>L23 vs. L4: p = 0.155 (ns)<br>L23 vs. L5: p = 0.1644 (ns)<br>L23 vs. L6: p = 0.0527 (ns)<br>L23 vs. $PYN^{L23}$: p = 0.0527 (ns)<br>L4 vs. L5: p = 0.0045 (**)<br>L4 vs. L6: p = 0.0007 (***)<br>L4 vs. $PYN^{L23}$: p = 0.5773 (ns)<br>L5 vs. L6: p = 0.5841 (ns)<br>L5 vs. $PYN^{L23}$: p = 0.0009 (***)<br>L6 vs. $PYN^{L23}$: p = 0.0001 (***) |
| Ex 10d: VIP->JAM2(LAMP5+ cINs) | L1: 0.65 ± 0.4914<br>L23: 17.07 ± 5.026<br>L4: 14.5 ± 3.973<br>L5: 25.25 ± 11.57<br>L6: 14.43 ± 5.937<br>$SST^{L45}$: 232.1 ± 49.32<br><br>N = 7, $LAMP5^{L1}$ = 8; $LAMP5^{L23}$ = 12; $LAMP5^{L4}$ = 6; $LAMP5^{L5}$ = 11; $LAMP5^{L6}$ = 9; $SST^{L45}$ = 12 cells. | Kruskal-Wallis test: p < 0.0001 | Uncorrected Dunn's test:<br>L1 vs. L23: p = 0.0067 (**)<br>L1 vs. L4: p = 0.0223 (*)<br>L1 vs. L5: p = 0.0175 (*)<br>L1 vs. L6: p = 0.0367 (*)<br>L1 vs. $SST^{L45}$: p < 0.0001 (****)<br>L23 vs. L4: p = 0.9961 (ns)<br>L23 vs. L5: p = 0.751 (ns)<br>L23 vs. L6: p = 0.6149 (ns)<br>L23 vs. $SST^{L45}$: p = 0.0005 (***)<br>L4 vs. L5: p = 0.7979 (ns)<br>L4 vs. L6: p = 0.6773 (ns)<br>L4 vs. $SST^{L45}$: p = 0.0047 (**)<br>L5 vs. L6: p = 0.8424 (ns)<br>L5 vs. $SST^{L45}$: p = 0.0002 (***)<br>L6 vs. $SST^{L45}$: p = 0.0002 (***) |
